# Bias correction for inverse variance weighting Mendelian randomization

**DOI:** 10.1101/2021.03.26.437168

**Authors:** Ninon Mounier, Zoltán Kutalik

**Affiliations:** University Center for Primary Care and Public Health, University of Lausanne, Lausanne, 1010, Switzerland; Swiss Institute of Bioinformatics, Lausanne, 1015, Switzerland; Department of Computational Biology, University of Lausanne, Lausanne, 1015, Switzerland

**Keywords:** Mendelian randomization, Weak instrument bias, Winner’s curse, Sample overlap

## Abstract

Inverse-variance weighted two-sample Mendelian randomization (IVW-MR) is the most widely used approach that utilizes genome-wide association studies (GWAS) summary statistics to infer the existence and the strength of the causal effect between an exposure and an outcome. Estimates from this approach can be subject to different biases due to the use of weak instruments and winner’s curse, which can change as a function of the overlap between the exposure and outcome samples.

We developed a method (`MRlap`) that simultaneously considers weak instrument bias and winner’s curse, while accounting for potential sample overlap. Assuming spike-and-slab genomic architecture and leveraging LD-score regression and other techniques, we could analytically derive, reliably estimate, and hence correct for the bias of IVW-MR using association summary statistics only.

We tested our approach using simulated data for a wide range of realistic settings. In all the explored scenarios, our correction reduced the bias, in some situations by as much as 30 folds. Additionally, our results are consistent with the fact that the strength of the biases will decrease as the sample size increases and we also showed that the overall bias is also dependent on the genetic architecture of the exposure, and traits with low heritability and/or high polygenicity are more strongly affected. Applying `MRlap` to obesity-related exposures revealed significant differences between IVW-based and corrected effects, both for non-overlapping and fully overlapping samples.

Our method not only reduces bias in causal effect estimation but also enables the use of much larger GWAS sample sizes, by allowing for potentially overlapping samples.

## 1 Introduction

Mendelian randomization (MR) is a method that uses genetic variants (typically single nucleotide polymorphisms, SNPs) as instrumental variables (IVs) to infer the existence and the strength of the causal effect between an exposure and an outcome [1]. In particular, two-sample summary data MR [2], which requires solely genome-wide association study (GWAS) summary statistics, has become increasingly popular. The reason for this is that in the last decade, GWASs have drastically increased in sample size [3] and the resulting summary statistics are often publicly available. This allows not only the identification of genetic variants independently associated with a particular exposure, i.e. IVs, but also the look up of the effect of such variants on a wide range of outcome traits in different samples. Each IV provides an independent estimate for the causal effect and these estimates can then be combined using an fixed effect, inverse variance-weighting (IVW) meta-analysis [2]. MR relies on three main assumptions (Figure 1): (1) Relevance – IVs must be robustly associated with the exposure. (2) Exchangeability – IVs must not be associated with any confounder of the exposure-outcome relationship. (3) Exclusion restriction – IVs must be independent of the outcome conditional on the exposure and all confounders of the exposure-outcome relationship.

**Figure 1:**
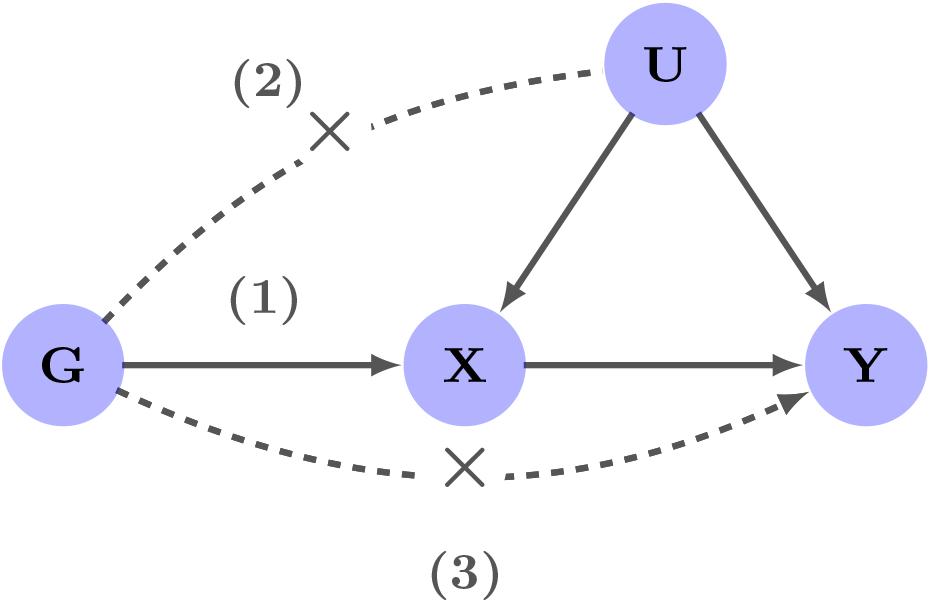
Main assumptions of Mendelian randomization. (1) Relevance – instrumental variables (IVs), denoted by *G*, are strongly associated with the exposure. (2) Exchangeability – *G* is not associated with any confounder of the exposure-outcome relationship. (3) Exclusion restriction – *G* is independent of the outcome conditional on the exposure and all confounders of the exposure-outcome relationship (i.e. the only path between the IVs and the outcome is via the exposure).

Additionally, two-sample MR methods expect the exposure and the outcome GWAS summary statistics to be obtained from independent samples. Sample overlap acts as a modifier for two well-known sources of bias in MR, weak instrument bias and winner’s curse. Two-sample MR estimates obtained from non-overlapping samples are known to be biased towards the null. When using overlapping samples, the causal effect estimate can be biased towards the observational correlation, which includes the correlation induced by confounders [4]. Most MR methods assume that SNP-exposure effects are measured without noise (NO Measurement Error, NOME assumption) [5]. This simplification leads to regression-dilution bias, which increases as the instruments get weaker. For this reason, the bias introduced by the NOME assumption is referred to as weak instrument bias and it becomes more and more severe as the average variance of the exposure explained by the IVs decreases [6]. When combined with sample overlap, the effect of weak instrument bias will move towards the observational correlation [7,8]. Using IVs strongly associated with the exposure and/or increasing the sample size can mitigate weak instrument bias [7]. Although the exact multiplicative bias due to the NOME assumption can be expressed analytically (proportional to the inverse of the F-statistic), the estimator for the multiplicative constant has typically high variance and works poorly in practice [5]. The simulation-extrapolation based SIMEX method proved to yield more robust corrections both for IVW [9] and MR-Egger estimates [5]. More recently, methods that incorporate measurement errors, such as MR-RAPS [10] or dIVW [11], have been proposed to tackle weak instrument bias. In addition, MR estimates are subject to winner’s curse, which occurs when the same sample is used to select IVs and estimate their effect on the exposure. In such case, the observed IV effect on the exposure is not an unbiased estimator for its true effect and is likely to be overestimated (in absolute value). This would affect the causal effect estimate (underestimation in non-overlapping sample and bias towards the observational correlation in fully overlapping samples) [8]. Using a third independent sample to select instruments [12], and therefore avoid winner’s curse, is a valid solution, but summary statistics from such additional samples are rarely available. Based on the expectation of truncated normal distribution [13], a correction can be applied for the SNP-exposure effect sizes. However, the additional estimator variance such correction entails can outweigh the benefit of the reduced bias, which can be mitigated by directly maximising the conditional likelihood [14]. Still, all these methods account for winner’s curse for a single SNP, but do not model the bias induced by the IV selection process from millions of potential markers, which is far more complex and depends on the underlying genetic architecture of the exposure. While previous approaches aimed at tackling one bias at a time, the intricate way these different sources of biases interplay with each other remains poorly understood, and there is currently no method that simultaneously handles them. To fill this gap, we propose a new method called `MRlap`, which is a summary statistics-based MR framework that simultaneously takes into account weak instrument bias and winner’s curse, while accounting for potential sample overlap, which modifies these biases.

Another major source of bias in MR is pleiotropy. In the presence of uncorrelated pleiotropy (reducing the exclusion restriction to the InSIDE assumption [15]), the causal effect estimate from IVW-MR is still consistent. Correlated pleiotropy (which can be induced by the existence of a genetic confounder acting on both the exposure and the outcome), however, can lead to the violation of the second assumption and more severe biases. Some approaches can be used to relax this assumption by assuming that at least 50% of the instruments are valid [16] or that non-pleiotropic instruments are the most frequent [17]. Since our `MRlap` approach does not explicitly tackle pleiotropy, we compared the impact of different sources of biases (pleiotropy, weak instrument bias, winner’s curse in the presence of potential sample overlap) on IVW-MR, MR-RAPS, dIVW, weighted median, weighted mode and `MRlap`.

In this paper, we will first introduce a two-sample MR framework that simultaneously takes into account weak instrument bias and winner’s curse, while accounting for potential sample overlap and its effect as a modifier of these biases, in order to obtain a corrected causal effect estimate. We will then test our approach and compare the proposed correction of the IVW-MR causal effect estimate against its uncorrected counterpart (and some pleiotropy-robust methods) using a wide range of simulation settings, including scenarios with pleiotropy. Finally, to demonstrate its utility, we will apply our approach to obesity-related traits using UK Biobank [18] (UKBB) data.

## 2 Results

### 2.1 Overview of the method

We propose a two-sample MR framework that takes into account two sources of bias: weak instrument bias and winner’s curse, while simultaneously accounting for potential sample overlap and its effect as a modifier of these biases. We analytically derived the expectation of the observed effect for IVW-based MR estimate:

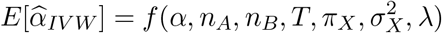

which depends on the true causal effect size (*α*), the sample sizes of the exposure and outcome GWASs (*n_A_, n_B_*), the threshold used to select IVs (*T*), the cross-trait LDSC intercept (λ, which depends on the degree of the sample overlap, the true causal effect and the strength of the confounder) and the genetic architecture of the exposure (characterised by the polygenicity *π_X_* and per-variant heritability 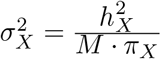, M being the number of potential IVs). All parameters (except *α*) are either known or can be estimated from the data. This allows us to adjust the IVW causal effect estimate with the aim of making it unbiased.

### 2.2 Simulations

Simulation results under our standard settings show a large discrepancy of the IVW-based causal effects estimated using different degrees of sample overlap, while the corrected effects are more closely aligned with the true causal effects (Figure 2 - A, Table S2). We observe a 10% overestimation of the causal effect for fully overlapping samples, and a 15% underestimation of the causal effect for non-overlapping samples, due to winner’s curse and weak instrument bias both biasing the estimate towards the null. As expected from (9), the bias is larger when using less stringent thresholds T. For all the thresholds tested, the ratio of the between-group and the within-groups variances is larger (up to 26 times) for IVW-based effects than for corrected effects (Table S3), highlighting the differences in IVW-based effects when estimated using different degrees of overlaps. The fact that the within-group variance is 1.3 times higher for the corrected effects compared to the uncorrected counterpart is due to the slightly increased variance of the bias-corrected estimator. The RMSE of the IVW-based effects is very dependent on the degree of overlap (being larger for non-overlapping and fully overlapping samples) while the RMSE of the corrected effects is consistent across varying degrees of overlap and up to 1.75 times lower for non-overlapping samples (Figure 2 - B). The corrected effects also yield a much better coverage, close to 95% for all degrees of overlap (Figure 2 - C). The corrected effects are significantly different from the IVW-based effects for all overlaps values except 50% and all thresholds (Table S2). The absolute bias of the IVW-based effects goes up to 0.033 while for the corrected effect it is smaller than 0.012 for all overlaps and thresholds.

**Figure 2:**
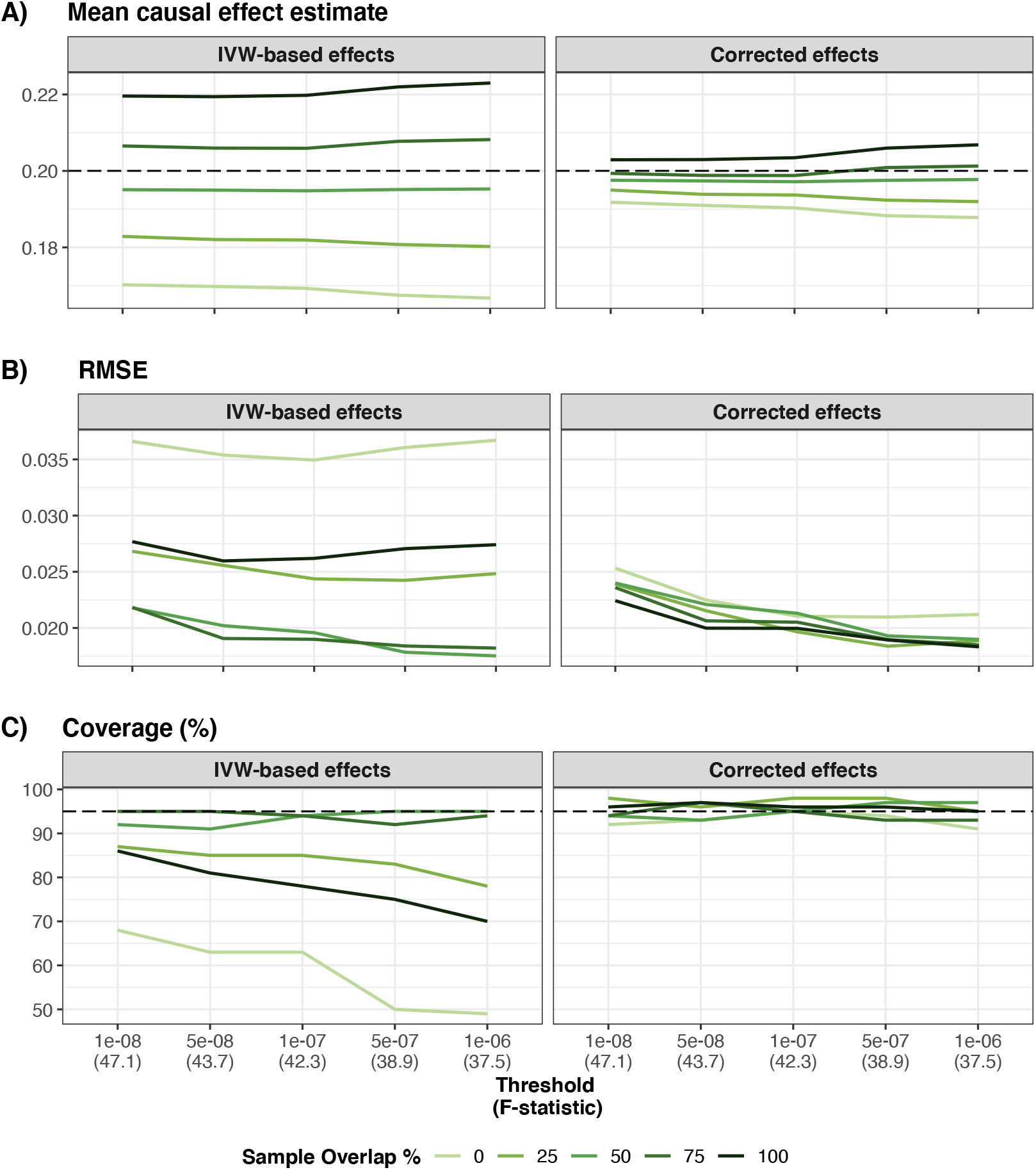
Simulation results for standard settings. *n_A_* = *n_B_* = 20, 000, *π_x_* = 0.001, 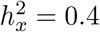, *κ_x_* = 0.3, *κ_y_* = 0.5, α = 0.2 Panel A) shows the mean IVW-based and corrected effect for each overlap and threshold obtained from 100 simulations (the dashed line represents the true causal effect). Panel B) shows the mean RMSE obtained for IVW-based and corrected effect for each overlap and threshold. Panel C) shows the coverage of the 95% confidence interval for IVW-based and corrected effect for each overlap and threshold.

The bias of IVW-based effects depends on the strength of the confounder. If the confounder is weak, IVW-based effects are mostly biased towards the null for low overlaps (Figure S5, Table S4, Table S5). When we simulated a stronger confounder, the bias of the IVW-based effects for fully overlapping samples increased (Figure S6, Table S6, Table S7). In both cases, the corrected and the IVW-based effects are significantly different for almost all overlaps and thresholds, and the corrected effects are substantially less biased than the IVW-based effects (Table S4, Table S6).

When the confounder effect (*ρ*:= *κ_x_* · *κ_y_*) and the causal effect (*α*) are acting in opposite directions, the results are particularly interesting because winner’s curse and weak instrument bias are biasing the results towards the null regardless of the sample overlap degree (Figure 3 - A, Table S8). In this case, IVW-based effects are more similar across the different degrees of sample overlap tested, but all are underestimating the true causal effect. For this reason, we do not observe a less striking decrease in the heterogeneity of the estimates across different sample overlaps upon correction (ratio of the between- and the within-group variance is 5 times larger for IVW-based effects), but still the correction reduces RMSE and bias, as well as leads to a better coverage, for all overlaps and thresholds (Figure 3, Table S9). In this scenario, IVW- based and corrected effects significantly differ for all overlaps and thresholds, with an average underestimation of 13% for IVW-based effects (Table S9).

**Figure 3:**
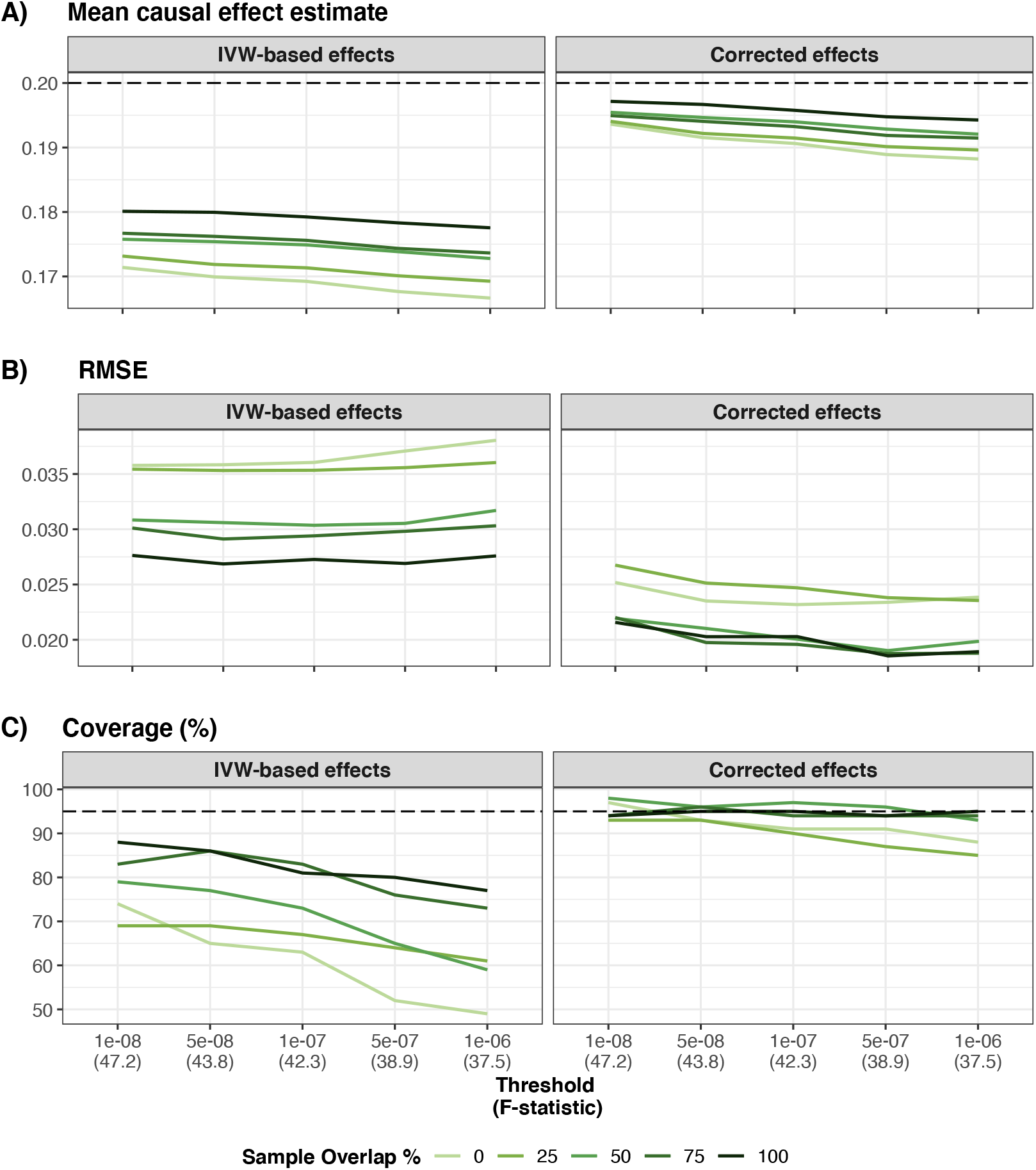
Simulation results for a scenario with a negative confounder. *n_A_* = *n_B_* = 20, 000, *π_x_* = 0.001, 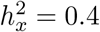, *κ_x_* = −0.3, *κ_y_* = 0.5, α = 0.2 Panel A) shows the mean IVW-based and corrected effect for each overlap and threshold obtained from 100 simulations (the dashed line represents the true causal effect). Panel B) shows the mean RMSE obtained for IVW-based and corrected effect for each overlap and threshold. Panel C) shows the coverage of the 95% confidence interval for IVW-based and corrected effect for each overlap and threshold.

In the absence of a true causal relationship between the exposure and the outcome, IVW-based effects from non-overlapping samples are unbiased. However, for fully overlapping samples, the IVW-based effects are biased towards the confounder-induced correlation (Figure S7, Table S10, Table S11). We showed that for large overlap percentages (≥ 50%), the corrected effects are significantly different from the IVW-based effects (60% smaller), and they are less biased for all overlaps and thresholds. Moreover, while the false positive rate (at 5% level) for IVW-based effects using non-overlapping samples is below 5%, it is much larger (between 20 and 40% depending on the threshold) when using fully overlapping samples. The correction proposed here provides a much better control of false positive rate for all degrees of sample overlap (Figure S8).

Simulating binary, instead of continuous, exposure led to very similar results. The absolute bias was in average 2.3 times larger for IVW-based effects than for corrected effects (Table S12) and the causal effect estimates were much more consistent across varying degrees of overlap after correction (Figure S9, Table S13). Results obtained using more realistic parameters in terms of sample sizes, genetic architecture and causal effect strength show a similar pattern. We observe an important bias of IVW-based causal effects, mostly when estimated from nonoverlapping (22% underestimation) or fully overlapping samples (30% overestimation), that is strongly reduced when using our correction (Figure S10, Table S14, Table S15). Corrected effects significantly differ from IVW-based effects for the most extreme overlaps values (0%, 75%, 100%) for which corrected effects are on average 5 times less biased than IVW-based effects.

We compared corrected effects obtained using the full (overlapping) sample to IVW-based effects obtained by splitting it into two halves to avoid sample overlap. We showed that the IVW estimators were 3.27 times more biased than those from `MRlap` and the latter reduced the estimator variance by more than three-fold (3.43) thanks to the elevated sample size. In addition, the corrected effects estimates had better coverage (88% vs 72%) and higher power (100% vs 94%). Thus we have demonstrated that applying `MRlap` to the full (overlapping) sample is a better strategy than ensuring no sample overlap and applying the IVW estimator, because it is less biased and has considerably lower variance (Table S16).

We also investigated the extent of the bias, for both IVW-based and corrected effects, in scenarios with either uncorrelated pleiotropy (Table S17, Figure S11, Table S18), moderate correlated pleiotropy (Table S19, Figure S12, Table S20) and strong correlated pleiotropy (Table S21, Figure S13, Table S22). Results obtained in presence of uncorrelated pleiotropy are very similar to the ones obtained for the standard settings, with slightly larger within group variances. MR-RAPS and dIVW however did not perform particularly well, even in the standard settings scenario, potentially because the instruments are relatively strong and their estimates are therefore mostly affected by winner’s curse. Using a third sample to select instruments and avoid winner’s curse was out of the scope of the paper, but we did try to use alternative selection approaches for these methods. First, we used a less stringent selection threshold (p-value threshold of 1e — 04, Figure S14) and we also ignored the selection step (only pruning the SNPs to obtain independent instruments, but not using any selection threshold, (Figure S14) to reduce the impact of winner’s curse, as suggested in Ye *et al* [11]. Surprisingly, the causal effect estimates for both methods were more strongly biased than when using a p-value threshold of 5e — 08. In presence of correlated pleiotropy, we observe a bias towards the ratio of the genetic confounder effects on ***Y*** and ***X*** (0.75 for moderate correlated pleiotropy and 1.4 for strong correlated pleiotropy) for both IVW-based and corrected effects for all sample overlap degrees. This can be explained by the fact that there is now a third source of bias, correlated pleiotropy, affecting the causal effect estimates. This source of bias is independent of sample overlap, and corrected effect estimates are able to recover consistent causal effect estimates across varying degrees of overlap, corresponding to the sum of the true causal effect and of the bias induced by correlated pleiotropy. When comparing our results to results obtained using pleiotropy-robust methods (Figure 4), we observe that for non-overlapping samples, Weighted Median, Weighted Mode and MR-RAPS estimates are able to recover the value of the IVW-based estimate from the standard settings scenarios (i.e. a downward biased causal effect estimate). However, for large degrees of sample overlap (≥ 50%), these approaches are biased towards the observational correlation, and their causal effect estimates are strongly overlap-dependent.

**Figure 4:**
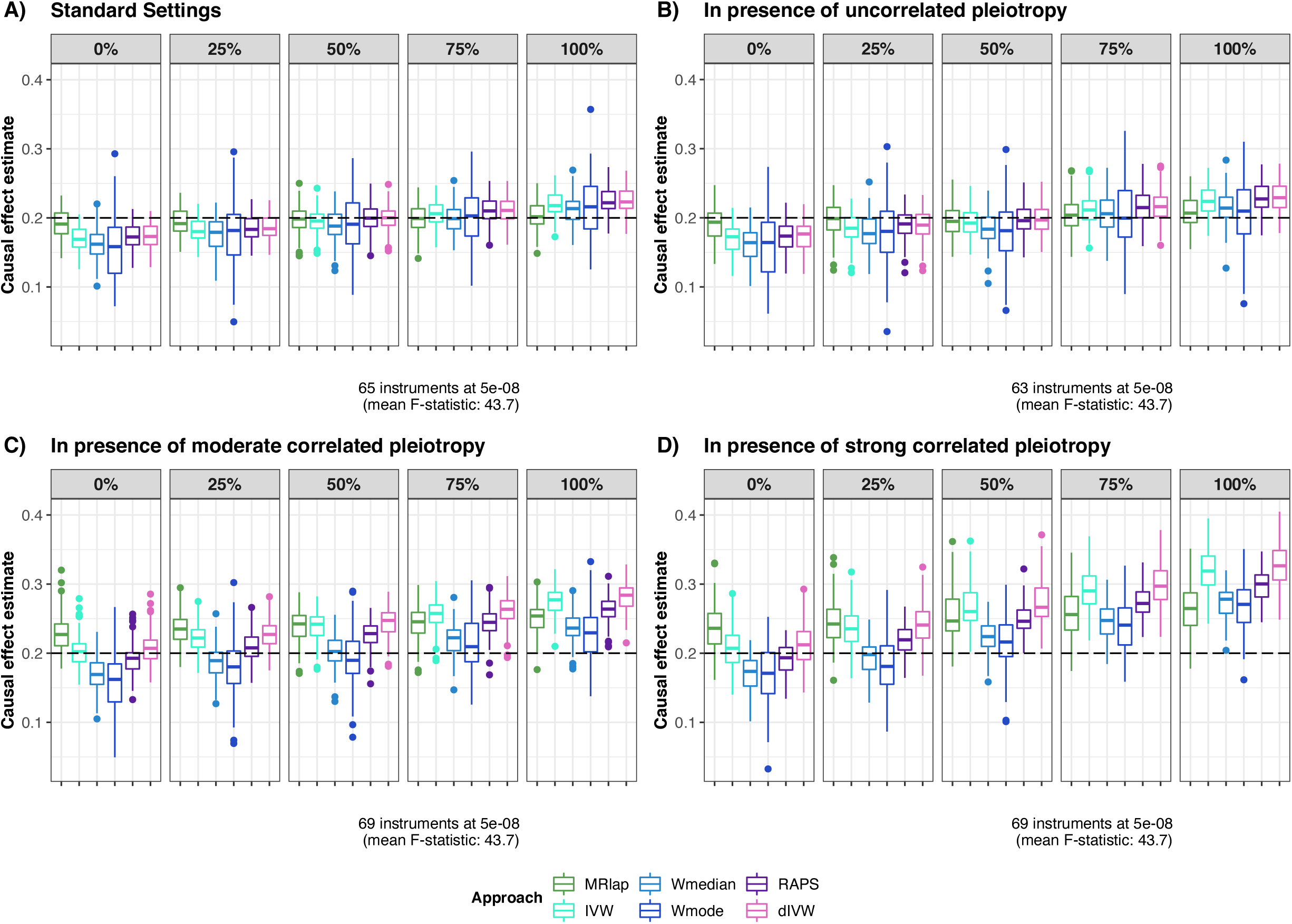
Comparison of different MR approaches. Causal effects estimates were obtained from 100 simulations using 6 different methods (MRlap in green, IVW in turquoise blue, Weighted Median in light blue, Weighted Mode in dark blue, MR-RAPS in purple and diVW in pink). The dashed line represents the true causal effect. Panel A) shows results for the standard settings scenario (no pleiotropy). Panel B) shows results in presence of uncorrelated pleiotropy. Panel C) shows results in presence of moderate correlated pleiotropy. Panel D) shows results in presence of strong correlated pleiotropy. The average number of instruments and mean F-statistic (at 5e-08) are indicated for each scenario.

### 2.3 Application to UKBB

We tested our method on UKBB obesity-related exposures using a similar approach and splitting the full dataset into samples of varying degrees of overlap. We started by estimating the causal effect of BMI on BMI and the causal effect of SBP on SBP as these are expected to be equal to 1. In this case, we only looked at non-overlapping samples and compared the IVW-based and the corrected effects to the true expected effect (Figure 5, Table S23 and S24). IVW-based effects are biased towards the null for all thresholds (95% confidence intervals do not include 1, coverage between 0 and 18%), with the bias being stronger for less stringent thresholds. Corrected effects however were less biased (at the cost of a slightly higher variance) and non-significantly different from 1 for all thresholds (coverage of between 65 and 96%), illustrating the importance of correcting for weak-instrument bias and winner’s curse even in the absence of sample overlap.

**Figure 5:**
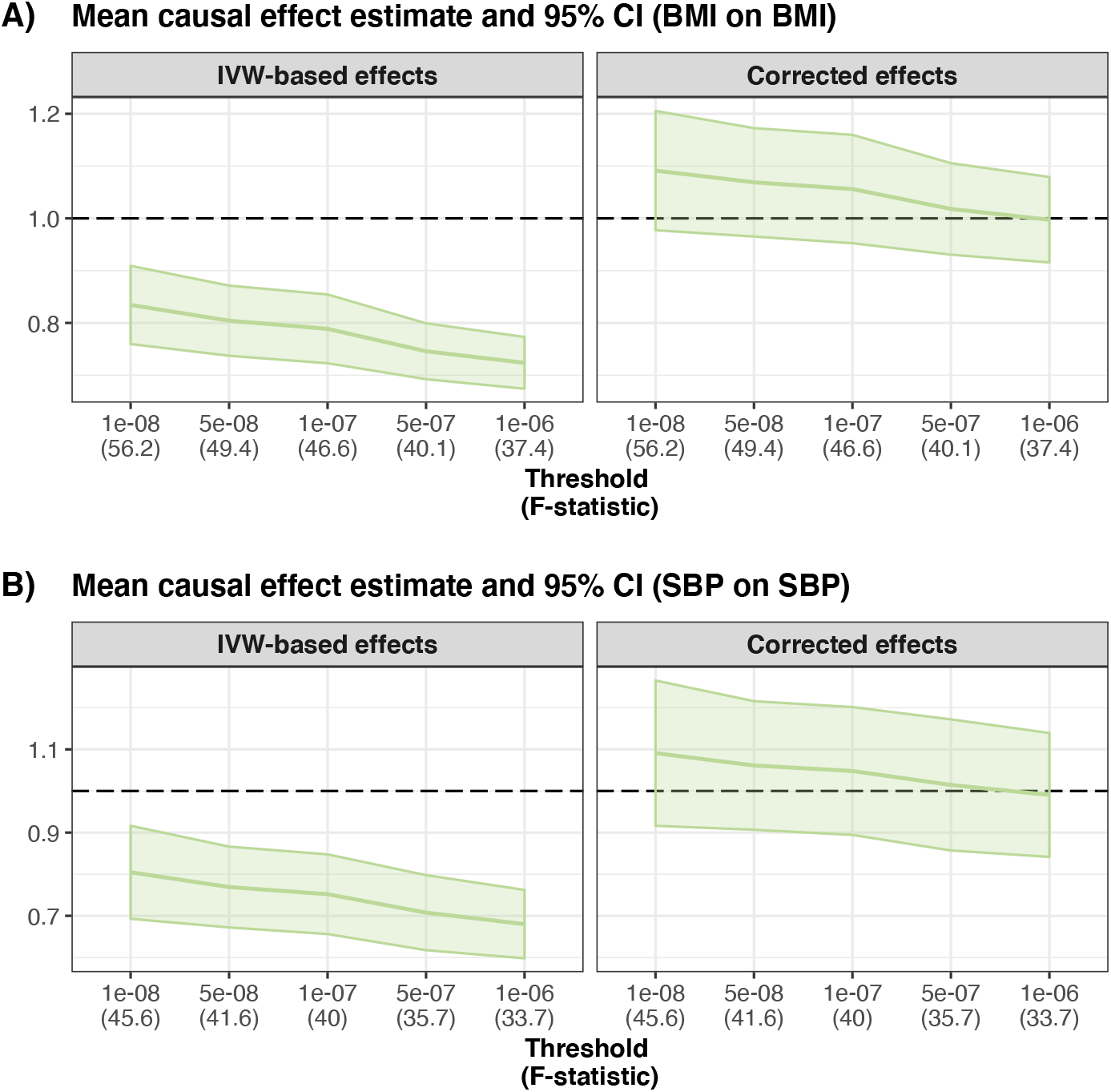
Effect of BMI on BMI and effect of SBP on SBP. This figure shows the mean IVW-based and corrected effect (and 95% confidence interval) for each threshold obtained from 100 different sampled datasets, using non-overlapping samples. The true causal effect is expected to be 1, as represented by the dashed line. Panel A) corresponds to the effect estimates of BMI on BMI. Panel B) corresponds to the effect estimates of SBP on SBP.

When looking at the IVW-based effect of BMI on SBP (Figure 6 - A), we observed that the estimates obtained using different p-value thresholds vary considerably, independently of sample overlap. Even though we expect an increase in bias when reducing the threshold used, as shown in simulations, here we see that for non-overlapping sample the IVW-based effect is larger for less stringent thresholds. This is inconsistent with winner’s curse and weak instrument bias that lean the estimate towards the null and we would expect the IVW-based effects for less stringent thresholds to be smaller (in absolute value). Hence, we believe that this phenomenon is not related to any of the biases discussed here and is due to other reasons such as the existence of multiple causal effects depending on exposure sub-type or the presence of a heritable confounder (see Discussion). Here we will focus on the results obtained using a p-value threshold of 5e-8.

**Figure 6:**
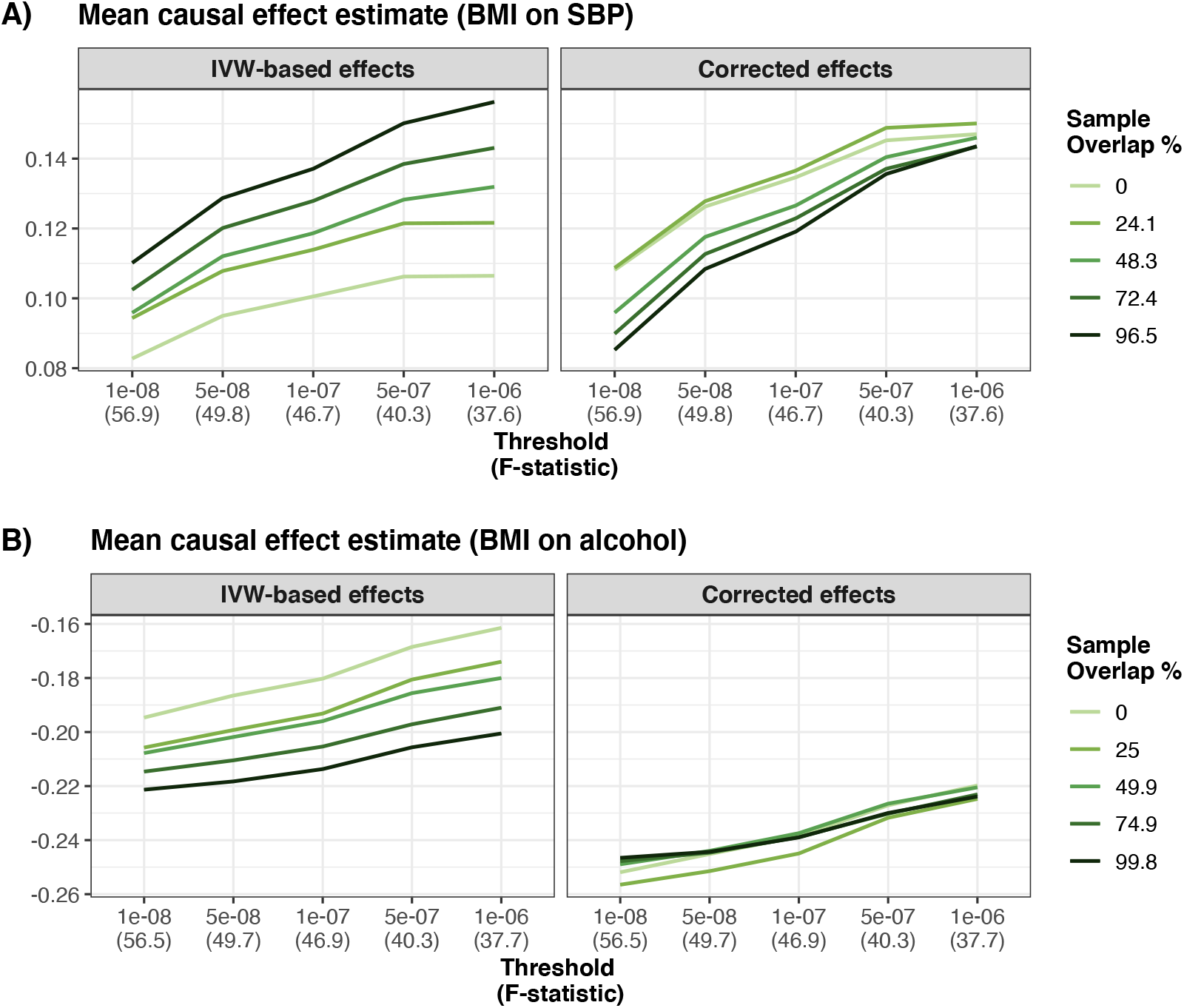
Effect of BMI on SBP and effect of BMI on alcohol intake frequency. This figure shows the mean IVW-based and corrected effect for each overlap and threshold obtained from 100 different sampled datasets. Panel A) corresponds to the effect estimates of BMI on SBP. Panel B) corresponds to the effect estimates of BMI on alcohol.

When using IVs reaching genome-wide significance, the IVW-based effects range between 0.095, for non-overlapping samples, and 0.129, for fully overlapping samples (Table S24). After correction, the range of the estimated effect is about 2 times smaller (0.108 ∓ 0.128). The better agreement of corrected effects across overlaps can be seen by looking at the ratio between the between groups and the within groups variance which is reduced 4-fold upon correction (Table S26). For non-overlapping samples, the difference between IVW-based and corrected effect is significant (*p*_diff_ = 0.0049), and standard two-sample MR underestimates the causal effect by about 25% compared to the corrected effect (the IVW-based effect is 0.095 while the corrected effect is 0.126). For fully overlapping samples, there is no significant difference between IVW-based and corrected effects. We observed a similar pattern when investigating the effect of BMI on smoking (Figure S16, Table S26, Table S28) where the largest bias occurs for non-overlapping samples. The IVW-based causal effect at p=5e-8 is 0.130 for non-overlapping samples while results after correction point towards a causal effect of 0.170 (underestimation of 24% - significant difference between IVW-based and corrected effects, *p*_diff_ = 0.0137). We do not see a significant difference between IVW-based and corrected effects for larger overlap values, but it is important to note that in this case, because of the impact of missing data on our design, the largest possible overlap was only 48.5%.

Our results implicate the existence of an environmental confounder biasing the causal effect estimate of BMI on alcohol intake frequency. At p=5e-8, IVW-based effects range between −0.187 for non-overlapping samples and −0.218 for fully overlapping samples, whereas the corrected effects are larger (−0.244 to −0.2515) (Figure 6 - B, Table S28). The between-group variance is much smaller for the corrected effects (Table S30). The difference between IVW-based and corrected effects is significant for all overlaps and the corrected effects being stronger than the IVW-based ones hints at the existence of an environmental confounder having a concordant effect on BMI and alcohol intake frequency, biasing all estimates towards the null (as shown in simulations, Figure 3). While we could not identify any plausible confounder of this relationship, its existence is supported by the fact that the observational correlation between BMI and alcohol intake frequency (−0.13 among the 379,530 genetically British individuals in the UKBB) is weaker than the standardised causal effect.

## 3 Discussion

We developed a method that models the entire instrument selection process, sample overlap and exposure effect estimation error. As a result, our approach reduces winner’s curse and weak instrument bias, even when the degree of overlap is unknown. Descriptive work have recently focused on a single relationship, investigating the effect of winner’s curse in the estimation of the causal effect of BMI on coronary artery disease, including comparison of the effect estimates for non-overlapping and fully overlapping samples [19]. To the best of our knowledge, while some empirical work describing the interplay between these biases has been done [20], no other method can tackle all the aforementioned biases while using summary statistics from exposure and outcome GWASs with arbitrary sample overlap. We tested our approach using a wide range of simulations scenarios: varying the strength of the causal effect, the strength of the confounder effect, sample sizes for the exposure and the outcome, as well as the genetic architecture of the exposure and demonstrated that both estimates for non-overlapping and fully overlapping samples can be biased. The direction and the magnitude of the bias depends on sample overlap and is strongly influenced by the effect of the confounder. When the confounder and the causal effect are acting in the same direction, observed effects are overestimated for fully-overlapping samples and underestimated for non-overlapping samples. However, when they are acting in opposite directions, these are underestimated for all overlaps because the direction of the biases is towards the null for any degree of sample overlap. We also showed that in the absence of a causal effect, results from overlapping samples would be biased, potentially leading to elevated type I error.

The correction we proposed worked remarkably well under all scenarios, and allows to drastically reduce the bias. For standard settings for example, we observed a 15% overestimation for fully overlapping samples and 10% underestimation for non-overlapping samples, that were respectively reduced to a 5% overestimation and a 2% underestimation after correction. We also found significant differences between IVW-based and corrected effects for fully-overlapping samples under all scenarios. For non-overlapping samples, IVW-based and corrected effects were significantly different under all scenarios expect in the case of the absence of causal effect. The decreased bias lead to an better coverage of the 95% confidence interval, but the correction also comes with an increased variance. Still, in all of our simulation scenarios the correction yielded reduced estimation error (RMSE) for at least one, if not all, sample overlap degrees. Moreover, while the RMSE of IVW-based effects strongly depends on the degree of overlap (because the bias is overlap-dependent), the RMSE of corrected effects is very similar for all overlaps. In the simulations, we mostly compared our corrected effect to its IVW-based counterpart, and only considered methods that can deal with weak instruments, such as MR-RAPS [10] and dIVW [11], for a small number of scenarios. Comparisons with these approaches are not optimal, as they require a third sample to be robust to winner’s curse, and are expected to perform better using a larger set of weaker instruments. We only investigated `MRlap` performance for thresholds smaller than 1e — 06 since using less stringent thresholds will increase the chances of using IVs that would violate the relevance assumption. We believe that the thresholds that have been used, both for simulations and real data analyses are realistic and that more lenient thresholds are unlikely to be used for IVW-MR in practice.

For real data, we used a sampling strategy to compare results obtained using varying degrees of sample overlap. We first focused on ‘same-trait’ (BMI on BMI and SBP on SBP) analyses for which the true causal effect is expected to be 1, using non-overlapping samples. IVW-based effects were strongly biased towards the null (between 18% and 30% depending on the threshold) and it is important to note that the 95% confidence intervals did not overlap with 1 for any of the traits. Corrected effect were less biased and their confidence intervals were overlapping with one (partly because of the lower precision of the estimator). We observed a slight over-correction for most stringent thresholds that could be due to potential violations of our assumptions regarding the genetic architecture of the exposure (spike-and-slab distribution). For the three other relationships we looked at, strong discrepancies were observed for low degrees overlap, with significant differences between IVW-based and corrected effects. This means that standard two-sample MR settings often lead to an underestimation of the true causal effect, that can be corrected using our approach. We also demonstrated that while most studies are extremely keen on avoiding any sample overlap while performing two-sample MR analysis fearing potential bias, the bias is often much less substantial for higher degrees of sample overlap. Among our many examples, we found that the IVW-MR estimate for the effect of BMI on alcohol intake frequency using the fully-overlapping samples is biased by a confounder. In this case, the confounder and the causal effect were acting in opposite direction, leading to an underestimation of the IVW-based effect for all overlaps. We have also highlighted that there is an important heterogeneity in causal effect estimates that vary with the IV selection threshold, due to heterogeneity in the estimates between the groups of genetic variants used for different thresholds. This can happen if there is a strong phenotypic heterogeneity in the exposure, in which case different groups of IVs could be affecting the exposure through different pathways [21]. Alternatively, in the presence of a genetic confounder, IVs picked up at a less stringent thresholds may be associated to a confounder, hence violating the second assumption of MR. Such phenomenon is out of the scope of our paper. In such case, IVW two-sample MR estimates would be biased, and more sophisticated approaches either specifically accounting for this genetic confounding (CAUSE [22], LHC-MR [23]) or others allowing for multiple causal effects (MR-Clust [21]) would be needed.

Our approach has its own limitations. As IVW-MR estimates, our corrected effect estimates will also be biased in case of the existence of a genetic confounder through which some of the selected instruments are primarily acting on the exposure, as shown in our simulations with correlated pleiotropy. In addition, our analytical derivation hinges on a genetic architecture of the exposure, namely assuming a spike-and-slab distribution of the multivariable effect sizes. Although this is a widely used and confirmed polygenic model, deviations from it could reduce the efficiency of our bias correction. It is also important to note that our work focused on continuous traits, and our approach would only work using case-control designs if the sample overlap degree do not differ between cases and controls. A simplification of the simulated model is that we assumed only a single confounder, but the bias estimation does not depend on this assumption. Finally, we have not explicitly modelled the local LD in the IV selection process, whereby a small Winner’s curse bias may be introduced when selecting the SNP with the strongest effect (at a given locus) as the IV.

Nowadays, samples from large biobanks are often used to estimate SNP effect sizes for both the exposure and the outcome, and hence it will be less and less possible to ensure that the two samples used are not overlapping. Thus, the need for non-overlapping samples forces researchers to use summary statistics from reduced sample size. While our results have shown that the biases usually do not strongly reduce power or change the clinical conclusions, which is line with results from Jiang et *al* [19], not accounting for these biases can still dramatically decrease coverage. For this reason, estimating the corrected effect using our approach (implemented in an R-package to facilitate its use) can be performed as a sensitivity analysis: if the corrected effect does not significantly differ from the IVW-based effect, then the IVW-MR estimate can be safely used (with the advantage of having lower variance). However, if there is a significant difference, corrected effects should be preferred as they are be less biased, independently of the sample overlap.

## 4 Methods

### 4.1 Expectation of the causal effect estimate

Let *X* and *Y* denote two random variables representing two complex traits. Genotype data is denoted by *G* and its jth column by ***g**_j_* (columns representing genetic variants, rows representing individuals). To simplify notation we assume that E [*X*] = E[*Y*] = E[*G*] = 0 and Var(*X*) = Var(*Y*) = Var(*G*) = 1. Let us assume that *X* is observed in sample *A* of sample size *n_A_, Y* is observed in sample *B* of sample size *n_B_* with an overlap of *n*_*A*⋂*B*_ individuals between the two samples. The vector of realisations of *Z^C^* is denoted by ***z***^*C*^ for all variables (*Z* = *X, Y, G, g*) and samples (*C* = *A,B,A* ⋂ *B*). Let us assume the following models

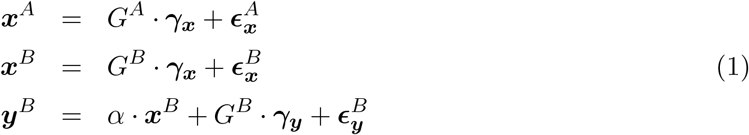

where *γ_x_* are the effect sizes of the genetic variants on X, *γ_y_* are their pleiotropic effects on Y. Assuming that there is a single environmental confounder *U* acting linearly on both traits (as used for simulations) the error term can split into two parts: 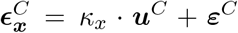 and 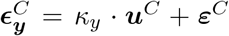, where *κ_x_* and *κ_y_* refer to the effect of U on X and Y respectively, *ε^c^* is independent of the confounder and C can take the values *A, B* or *A* ⋂ *B* as above.

Under the INSIDE assumption [15] (INstrument Strength Independent of Direct Effect, i.e. horizontal pleiotropic effects are independent of the direct effect), Cov (*γ_x_*, *γ_y_*) = 0 and E [*γ_y_*] = 0. We denote *ρ*: = Cov (*ϵ_x_, ϵ_y_*) = *κ_x_* · *κ_y_*. It corresponds to the part of the observational correlation (*r*) due to a (non-genetic) confounder (*r* = *ρ* + *α*). Note that genetic confounding as well as reverse causal effect are also affecting observational correlation, but as long as the instruments used for MR are not associated with the confounder nor the outcome, their effect would be captured by *ρ*.

The univariable effect size estimates from GWASs for genetic variant *j* are as follows

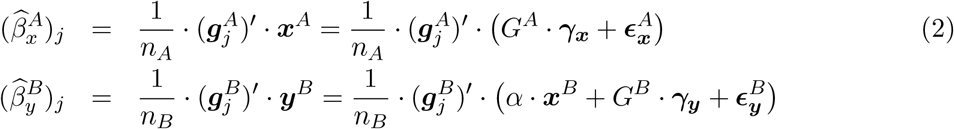

where the genotype data for genetic variant *j* for individuals in sample *A* is denoted by 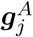.

We intend to use MR to estimate the causal effect of *X* on *Y*. We will use *m* linkage disequilibrium (LD)-independent genetic variants as IVs. Let us now consider the fixed-effect inverse-variance weighting meta-analysis for the ratio estimates for the causal effect α. Each IV *j* provides a ratio estimate

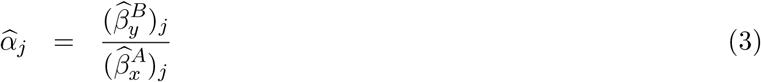

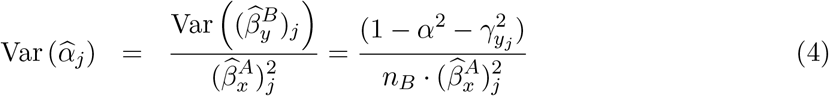

Hence the weights (*w_j_*) of IV *j* for estimating the IVW causal effect are:

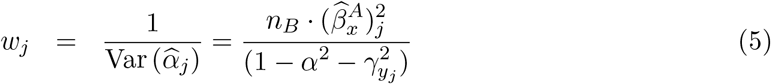

Finally, the estimate can be written in the following form

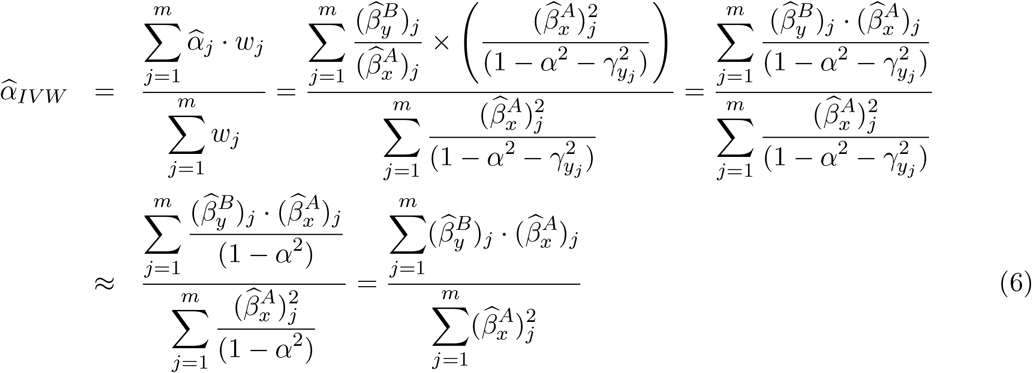

Here, the last approximation is based on the realistic assumption that the individual pleiotropic effect of each SNP is very small.

To account for winner’s Curse, we need to consider the probability of being selected for each genetic variant. Let us consider a threshold *T* (*T* = –*Φ*^-1^(*p*/2) ≈ 5.45 for *p* = 5 × 10^-8^, genomewide significance threshold, for example) and use only IVs with 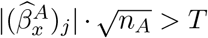. By denoting 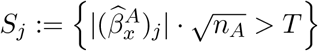, the causal effect estimate (6) changes to

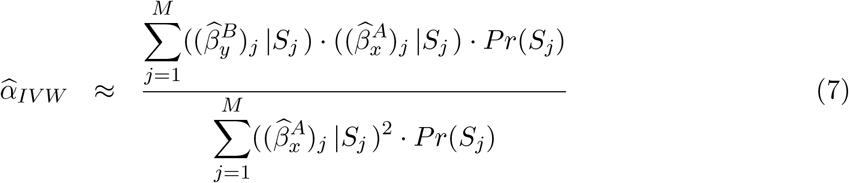

Note that while *m* denoted the number of IVs, *M* represents the number of genome-wide variants from which IVs are selected. By approximating the expectation of a ratio by the ratio of expectations (see Supplementary Section A for details about this assumption), the expectation of the causal effect estimate (7) can be written as

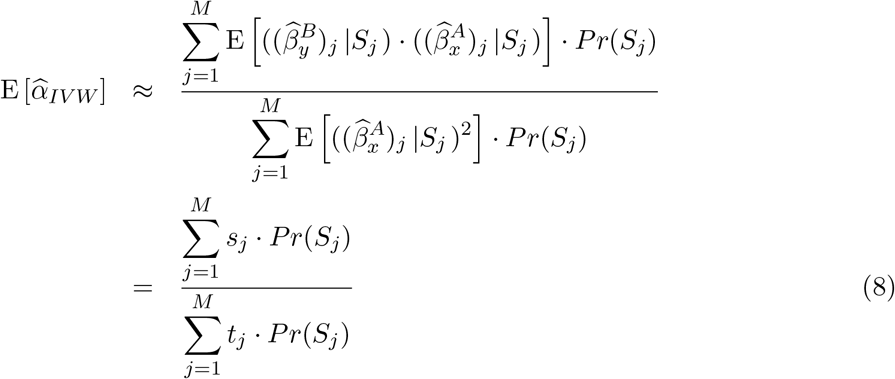

The values of *s_j_*, *t_j_* and *P_r_*(*S_j_*) can be analytically derived (presented in Supplementary Section A), and we show that 8 translates to

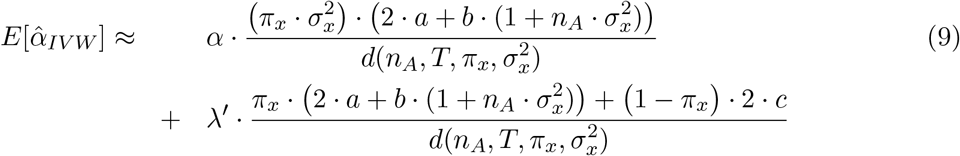

with

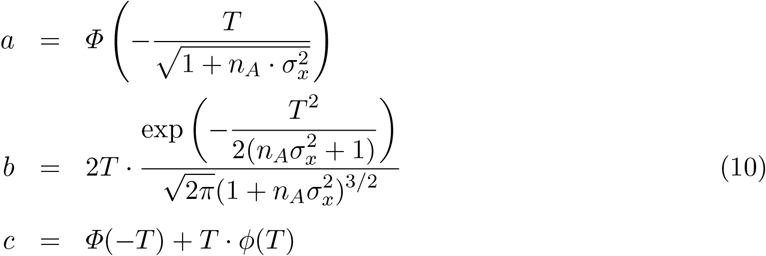

and

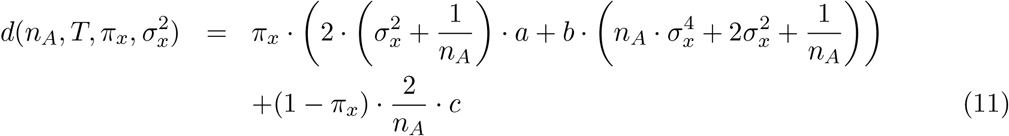

*π_x_* and 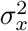 are characteristics of the genetic architecture of trait *X* (respectively, a measure of the polygenicity and the per-variant heritability, see (S35)) [24] and *λ*’ is a quantity closely related to the cross-trait LD score regression (LDSC) intercept [25] 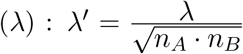. The constants *a, b* and *c* do not depend on the causal effect *α* nor the sample overlap since *n*_*A*⋂*B*_ is only affecting λ’ (see (10) and (S33)). The same is true for the denominator (11).

We can see that in the absence of sample overlap, the second term is equal to 0 and the bias is multiplicative. In this case, 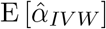 is lower than *α* and the IVW-based effect will be biased towards the null (Figure S1). The only parameters affecting the bias are *π_x_* and 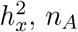 and *T*. When *π_x_* is smaller, or 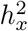 is larger, then IVs have stronger effects, leading to a smaller bias when all other parameters are kept constant. As expected, since these are commonly used approaches to limit weak instrument bias, using a more stringent threshold and/or increasing the exposure sample size reduce the bias.

When there is sample overlap (with % overlap defined as 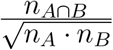, the expression of 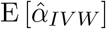 is more complex. The magnitude of the bias will not only depend on the parameters described for non-overlapping samples (Figures S2, S3 and S4), but also on the confounder’s effect (*ρ*) that can affect both the magnitude and the direction of the bias. For example, when the percentage of overlap is relatively low (20%) and *ρ* has the same sign as *α*, then the estimate will be further biased towards the null, whereas a confounder acting in the opposite direction (as the causal effect) will reduce the bias (Figure S2). When the percentage of overlap increases (Figures S3 and S4), the second term starts to dominate, and the bias direction strongly depends on *ρ*.

All parameters except *α* are known or can be estimated from the data. We assume that sample sizes for *X* and *Y* (respectively *n_A_* and *n_B_*) are known, as well as the threshold used to select IVs (*T*). Parameter *λ*’ can be estimated from cross-trait LDSC [25]. *π_x_* and 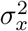 are estimated by matching the denominator of this formula, 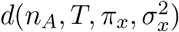, to the denominator of (8) (see Supplementary Section B for details).

From (9), we can derive a corrected effect for the causal effect

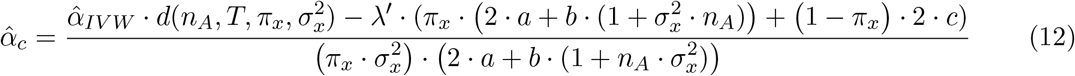

We also derived the standard error of the corrected effect as well as the covariance between IVW-based and corrected effects in Supplementary Section C. Note that the formula proposed by Burgess *et al.* [4] (Supplementary Section D) under the null is a special case of ours when all instruments are selected based on external data (i.e. there is no winner’s curse).

This approach has been implemented in an R-package (`MRlap`), using existing functions from the Two-Sample MR R-package [26] and the LDSC implementation from the GenomicSEM R- package [27]. All analyses presented in this paper have been performed using version 0.0.2. IVs were selected with for different T threshold and independent IVs were identified using distancepruning (500kb, that is equivalent to use LD-pruning with a LD cutoff of 0). For the LDSC analyses we used the 1000G LD-scores [28].

### 4.2 Simulations

We used UKBB [18] genotypic data and restricted our analyses to unrelated individuals of British ancestry (identified using genomic principal components) and HapMap3 genetic variants [29] (*M* ≈ 1,150,000) to simulate phenotypic data. From this set of 379,530 individuals, we first sampled the exposure dataset (*n_A_* individuals) and 5 different outcome datasets of sample size *n_B_*, with an overlap with the exposure dataset varying (from no overlap to full overlap, increasing in increments of 25%). Next, causal SNPs for the exposure were randomly drawn from the set of 1,150,000 genetic variants, based on the polygenicity of *X* (*π_x_*) and their effects were simulated using the heritability of 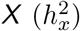 as follows

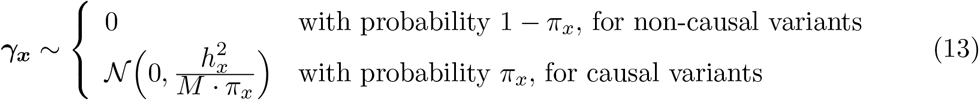

For simplicity, we assumed that there were no direct genetic effects on the outcome. Then, phenotypic data for *X* and *Y* were simulated for all individuals included in the exposure or in any of the outcome samples, taking into account the effect of the confounder *U* on *X* and *Y* (respectively *κ_x_* and *κ_y_*) and the causal effect of *X* on *Y* (*α*), using the following design

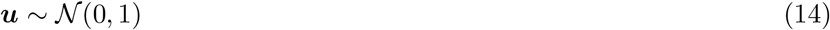

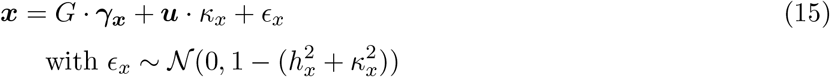

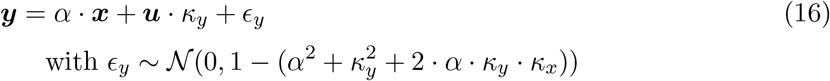

Note that this design ensures that both *X* and *Y* have a zero mean and a variance of 1. Slightly different designs, with the same property, were used to simulate phenotypic data when the exposure was binary (case-control) and in the presence of uncorrelated or correlated pleiotropy (Supplementary Section E). A GWAS was performed for each sample (one for the exposure and five GWASs for the outcome, one for each sample overlap) using BGENIE [30]. We then applied MRlap to these GWAS summary statistics to obtain the IVW-based and the corrected effect estimates.

For each parameter setting we tested, 100 datasets were simulated. Our standard parameter settings consisted of simulating data for *n_A_* = 20,000 and *n_B_* = 20,000 individuals. *X* was simulated with moderate polygenicity and large heritability (*π_x_* = 0.001 and 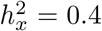). *U* had a moderate effect on both *X* and *Y* (*κ_x_* = 0.3, *κ_y_* = 0.5), leading to a correlation (*ρ*) of 0.15 induced by the confounder. We varied the size of the causal effect of *X* on *Y* from null (*α* = 0) to moderate (*α* = 0.2).

In addition to these standard settings, we explored various other parameter values. We investigated the effect of a confounder acting in an opposite direction (*κ_x_* = —0.3 and *κ_y_* = 0.5) and tested different strengths for the confounding factor (weaker: *κ_x_* = 0.15 and *κ_y_* = 0.3, and stronger: *κ_x_* = 0.5 and *κ_y_* = 0.8). We also considered a case-control design, where the exposure was first simulated on the liability scale before being converted to the observed scale for our analyses. For this setting we used larger sample sizes (*n_A_* = 100,000, *n_B_* = 100, 000) and a prevalence of 0.1 to define cases and controls. We explored a scenario with more realistic parameters: larger sample sizes (*n_A_* = 100, 000, *n_B_* = 100, 000), increased polygenicity (*π_x_* = 0.005), lower heritability 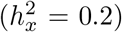 and a smaller causal effect (*α* = 0.1). Finally, we simulated data in presence of both uncorrelated and correlated pleiotropy. For the uncorrelated pleiotropy, we used the standard settings parameters and added direct genetic effects on *Y* (*π_y_* = 0.002 and 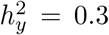). 60% of the SNPs having a direct effect on *X* were also directly affecting *Y* and their direct effects on each trait were uncorrelated. To simulate data in presence of correlated pleiotropy, we added to the model a genetic confounder (*U_g_*) acting both on *X* and *Y*, with respective effects *q_x_* and *q_y_*. First, we modelled a genetic confounder that was highly polygenic and fairly heritable (*π_u_* = 0.0001 and 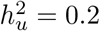), with moderate effects on the two traits (*q_x_* = 0.4 and *q_y_* = 0.3). Then we slightly increased the confounder’s polygenicity and, in order to make the genetic confounding effect stronger, we increased its heritability and its effects on *X* and *Y* (*π_u_* = 0.0005, 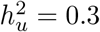, *q_x_* = 0.5 and *q_y_* = 0.7).

For each scenario, IVW-based and corrected causal effects were compared for different degrees of sample overlap and different instrument selection thresholds. Results quality was assessed using root-mean square error (RMSE), coverage and power.

For a given instrument selection threshold, we obtained 500 causal effect estimates: one for each of the 100 simulated data sets for each of the five degrees of sample overlap (ranging from 0 to 100%). Causal effect estimates should ideally not depend on the extent of overlap between the exposure and outcome samples. To quantify the extent to which this holds, we grouped estimates according to which sample overlap degree they came from and compared the between group variance relative to the within group variance of the estimates. A method that is robust to overlap between the exposure and outcome samples will have small between group variance relative to the variance of the estimator (characterised by the within group variance).

Finally, we tested for differences between IVW-based and corrected effects using the following test statistic

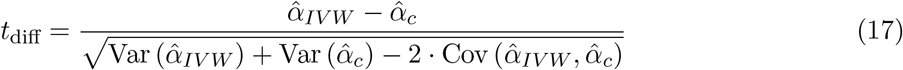

In addition to those scenarios, we wanted to assess the potential gain, in terms of bias and variance, arising from the possibility of performing analyses using the full UKBB sample instead of having to split it into two halves to avoid sample overlap. To do so, we simulated data using the following parameters: large polygenicity and moderate heritability (*π_x_* = 0.01 and 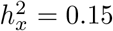), moderate confounder effect (*κ_x_* = 0.3, *κ_y_* = 0.5), and a fairly small causal effect (*α* = 0.1). In this case, we only compared the IVW-based causal effect estimates from non-overlapping samples (*n_A_* = 180,000, *n_B_* = 180,000, roughly half of UKBB sample size) and the corrected causal effect estimates from fully overlapping samples (*n_A_* = 360,000, *n_B_* = 360,000) by measuring the bias, the variance and the RMSE.

Finally, we also compared our `MRlap` results to those of pleiotropy-robust approaches, such as weighted median [16], weighted mode [17], MR-RAPS [10] and dIVW [19], both under the standard settings and in all scenarios with pleiotropy, for all degrees of sample overlap. For all methods, we use a p-value threshold of 5 × 10^-8^ to select IVs, and additional analyses were performed using MR-RAPS and dIVW with a threshold of 10^-4^ and without selection.

### 4.3 Application to UKBB

To assess the effect of sample overlap on real data, we used a design very similar to the one used for simulations. We used both genotypic and phenotypic data from UKBB [18] and restricted our analyses to the same subsets of individuals and genetic variants. From this set of individuals, we first sampled the exposure dataset (100,000 individuals) and 5 different outcome datasets (100,000 individuals), where the overlap with the exposure dataset varied (from no overlap to full overlap, increasing in increments of 25% - in the case of unequal sample sizes, the percentage of overlap for the samples proportionally increased from 0 to the maximum value attainable given the difference in sample sizes). Note that we always used the full set of individuals to create the 100,000-individuals samples, so the percentage of missing data in each sample will be the same as in the UKBB. Therefore, the total number of individuals with phenotypic data (effective sample size) will vary depending on the traits. First, we performed ‘same-trait’ analyses to estimate the causal effect of body mass index (BMI) on itself, and the causal effect of systolic blood pressure (SBP) on itself, using only the non-overlapping samples. In addition, for all degrees of sample overlap, we assessed the effect of BMI on SBP, on the number of cigarettes previously smoked daily (smoking) and on alcohol intake frequency (alcohol). For smoking, answers coded as “-10” or “-1” were considered as missing. For alcohol, answers were recoded to correspond to an increased intake frequency, and answers coded as “-3” were considered as missing. Details about the pairs of traits analysed are available in Table S1.

Phenotypic data was normalised (inverse-normal quantile transformed) and subsequently adjusted for the following covariates: sex, age, age× age and the first 40 principal components. Similarly to what we did for simulations, a GWAS was performed for each sample (the exposure dataset and the five outcome datasets, one for each sample overlap) using BGENIE [30]. We then applied `MRlap` to these GWAS summary statistics to obtain the IVW-based and the corrected effect estimates. In addition, since in this case a reverse causal effect (from the outcome on the exposure) could exist, we filtered out variants that were significantly more strongly associated with the outcome than with the exposure in order to remove potentially invalid IVs (Steiger filter).

We repeated this sampling approach a 100 times. For each repetition, IVW-based and corrected causal effects were compared using the within groups and between groups variances and we tested for differences between the IVW-based and the corrected effects using (17).

## Acknowledgements

The authors thank Chiara Auwerx, Liza Darrous, Sven Erik Ojavee, Marion Patxot, Eleonora Porcu, Marie Sadler and Tabea Schoeler for the helpful discussions and constructive comments. Computations have been performed on the HPC cluster of the Lausanne University Hospital.

This research has been conducted using the UK Biobank Resource under Application Number 16389.

## Author Contributions

Zoltaán Kutalik designed and supervised the project. Zoltaán Kutalik and Ninon Mounier developed the methods. Ninon Mounier performed the analyses. Ninon Mounier and Zoltáan Kutalik wrote the manuscript.

