## Supplementary Material for "Bias correction for inverse variance weighting Mendelian randomization"

#### Supplementary Materials

A. Derivation of the corrected effect estimate
B. Estimation of the standard error of the corrected effect estimate and of the covariance between IVW-based and corrected effects
C. Derivation of the corrected effect estimate
D. Simplification under the null
E. Additional simulation designs
References

Supplementary Figures
Supplementary Tables

[1]University Center for Primary Care and Public Health, University of Lausanne, 1010, Switzerland

[2]Swiss Institute of Bioinformatics, Lausanne, 1015, Switzerland

[3]Department of Computational Biology, University of Lausanne, Lausanne, 1015, Switzerland

### A. Derivation of the corrected effect estimate

Let $X$ and $Y$ denote two random variables representing two complex traits. Genotype data is denoted by $G$ and its $j$th column by $\boldsymbol{g}_j$ (columns representing genetic variants, rows representing individuals). To simplify notation we assume that $\mathrm{E}[X] = \mathrm{E}[Y] = \mathrm{E}[G] = 0$ and $\mathrm{Var}(X) = \mathrm{Var}(Y) = \mathrm{Var}(G) = 1$. Let us assume that $X$ is observed in sample $A$ of sample size $n_A$, $Y$ is observed in sample $B$ of sample size $n_B$ with an overlap of $n_{A\cap B}$ individuals between the two samples. The vector of realisations of $Z^C$ is denoted by $\boldsymbol{z}^C$ for all variables ($Z = X, Y, G, g$) and samples ($C = A, B, A \cap B$). Let us assume the following models

$$\begin{aligned} \boldsymbol{x}^A &= G^A \cdot \boldsymbol{\gamma_x} + \boldsymbol{\epsilon}_{\boldsymbol{x}}^A \\ \boldsymbol{x}^B &= G^B \cdot \boldsymbol{\gamma_x} + \boldsymbol{\epsilon}_{\boldsymbol{x}}^B \\ \boldsymbol{y}^B &= \alpha \cdot \boldsymbol{x}^B + G^B \cdot \boldsymbol{\gamma_y} + \boldsymbol{\epsilon}_{\boldsymbol{y}}^B \end{aligned} \tag{S1}$$

where $\boldsymbol{\gamma_x}$ are the effect sizes of the genetic variants on $X$, $\boldsymbol{\gamma_y}$ are their pleiotropic effects on $Y$. Assuming that there is a single environmental confounder $U$ acting linearly on both traits (as used for simulations) the error term can split into two parts : $\boldsymbol{\epsilon}_{\boldsymbol{x}}^C = \kappa_x \cdot \boldsymbol{u}^C + \boldsymbol{\varepsilon}^C$ and $\boldsymbol{\epsilon}_{\boldsymbol{y}}^C = \kappa_y \cdot \boldsymbol{u}^C + \boldsymbol{\varepsilon}^C$, where $\kappa_x$ and $\kappa_y$ refer to the effect of $U$ on $X$ and $Y$ respectively, $\varepsilon^C$ is independent of the confounder and $C$ can take the values $A, B$ or $A \cap B$ as above. Note that in the derivations below, we do not assume that the correlation between $\boldsymbol{\epsilon}_{\boldsymbol{x}}^C$ and $\boldsymbol{\epsilon}_{\boldsymbol{y}}^B$ is due to a single confounder, we simply split the error terms into correlated and uncorrelated parts.

Under the INSIDE assumption [1] (INstrument Strength Independent of Direct Effect, i.e. horizontal pleiotropic effects are independent of the direct effect), $\mathrm{Cov}(\boldsymbol{\gamma_x}, \boldsymbol{\gamma_y}) = 0$ and $\mathrm{E}[\boldsymbol{\gamma_y}] = 0$. We denote $\rho := \mathrm{Cov}(\boldsymbol{\epsilon_x}, \boldsymbol{\epsilon_y}) = \kappa_x \cdot \kappa_y$. It corresponds to the part of the observational correlation ($r$) due to a (non-genetic) confounder ($r = \rho + \alpha$). Note that genetic confounding as well as reverse causal effect are also affecting observational correlation, but as long as the instruments used for MR are not associated with the confounder nor the outcome, their effect would be captured by $\rho$.

#### a. Inverse Variance Weighted estimate

We intend to use MR to estimate the causal effect of $X$ on $Y$. We will use $m$ linkage disequilibrium (LD)-independent genetic variants as IVs. The GWAS effect size estimates for genetic variant $j$ are as follows

$$\begin{aligned} (\widehat{\beta}_x^A)_j &= \frac{1}{n_A} \cdot (\boldsymbol{g}_j^A)' \cdot \boldsymbol{x}^A = \frac{1}{n_A} \cdot (\boldsymbol{g}_j^A)' \cdot \left(G^A \cdot \boldsymbol{\gamma_x} + \boldsymbol{\epsilon}_{\boldsymbol{x}}^A\right) \\ (\widehat{\beta}_x^B)_j &= \frac{1}{n_B} \cdot (\boldsymbol{g}_j^B)' \cdot \boldsymbol{x}^B = \frac{1}{n_B} \cdot (\boldsymbol{g}_j^B)' \cdot \left(G^B \cdot \boldsymbol{\gamma_x} + \boldsymbol{\epsilon}_{\boldsymbol{x}}^B\right) \\ (\widehat{\beta}_y^B)_j &= \frac{1}{n_B} \cdot (\boldsymbol{g}_j^B)' \cdot \boldsymbol{y}^B = \frac{1}{n_B} \cdot (\boldsymbol{g}_j^B)' \cdot \left(\alpha \cdot \boldsymbol{x}^B + G^B \cdot \boldsymbol{\gamma_y} + \boldsymbol{\epsilon}_{\boldsymbol{y}}^B\right) \end{aligned} \tag{S2}$$

where the genotype data for genetic variant $j$ for individuals in sample $A$ is denoted by $\boldsymbol{g}_j^A$. To simplify the equations, we introduce the following notations

$$\begin{aligned}
\tau_{x_j}^A &:= \frac{1}{n_A} \cdot (\boldsymbol{g}_j^A)' \cdot \left((G^A)_{-j} \cdot \boldsymbol{\gamma}_{\boldsymbol{x}-j} + \boldsymbol{\epsilon}_{\boldsymbol{x}}^A\right) \\
\tau_{x_j}^B &:= \frac{1}{n_B} \cdot (\boldsymbol{g}_j^B)' \cdot \left((G^B)_{-j} \cdot \boldsymbol{\gamma}_{\boldsymbol{x}-j} + \boldsymbol{\epsilon}_{\boldsymbol{x}}^B\right) \\
\tau_{y_j}^B &:= \frac{1}{n_B} \cdot (\boldsymbol{g}_j^B)' \cdot \left((G^B)_{-j} \cdot \boldsymbol{\gamma}_{\boldsymbol{y}-j} + \boldsymbol{\epsilon}_{\boldsymbol{y}}^B\right)
\end{aligned} \tag{S3}$$

where $(\cdot)_{-j}$ subscripts refer to the full set of data except the $j$th element (column in case of matrices). Now we can reformulate the effect size equation (S2) as follows

$$\begin{aligned}
(\widehat{\beta}_x^A)_j &= \gamma_{x_j} + \tau_{x_j}^A \\
(\widehat{\beta}_x^B)_j &= \gamma_{x_j} + \tau_{x_j}^B \\
(\widehat{\beta}_y^B)_j &= \alpha \cdot (\gamma_{x_j} + \tau_{x_j}^B) + \gamma_{y_j} + \tau_{y_j}^B
\end{aligned} \tag{S4}$$

In the following, we will work out the first two moments of the $\tau$ variables. First, since $\boldsymbol{g}_j$ is orthogonal to $G_{-j}$ and all $\epsilon$ variables, the expectation of all $\tau$ variables is zero. Their variances can be calculated as follows

$$\begin{aligned}
\mathrm{Var}\left(\tau_{x_j}^A\right) &= \mathrm{Var}\left(\frac{1}{n_A} \cdot (\boldsymbol{g}_j^A)' \cdot \left((G^A)_{-j}\boldsymbol{\gamma}_{\boldsymbol{x}-j} + \boldsymbol{\epsilon}_{\boldsymbol{x}}^A\right)\right) \\
&= \frac{1}{n_A} \cdot \mathrm{Var}\left(g_j\right) \cdot \mathrm{Var}\left(G_{-j}\boldsymbol{\gamma}_{\boldsymbol{x}-j} + \boldsymbol{\epsilon}_{\boldsymbol{x}}^A\right) \\
&= \frac{1-\gamma_{x_j}^2}{n_A}
\end{aligned} \tag{S5}$$

$$\begin{aligned}
\mathrm{Var}\left(\tau_{x_j}^B\right) &= \mathrm{Var}\left(\frac{1}{n_B} \cdot (\boldsymbol{g}_j^B)' \cdot \left((G^B)_{-j}\boldsymbol{\gamma}_{\boldsymbol{x}-j} + \boldsymbol{\epsilon}_{\boldsymbol{x}}^B\right)\right) \\
&= \frac{1}{n_B} \cdot \mathrm{Var}\left(g_j\right) \cdot \mathrm{Var}\left(G_{-j}\boldsymbol{\gamma}_{\boldsymbol{x}-j} + \boldsymbol{\epsilon}_{\boldsymbol{x}}^B\right) \\
&= \frac{1-\gamma_{x_j}^2}{n_B}
\end{aligned} \tag{S6}$$

$$\begin{aligned}
\mathrm{Var}\left(\tau_{y_j}^B\right) &= \mathrm{Var}\left(\frac{1}{n_B} \cdot (\boldsymbol{g}_j^B)' \cdot \left((G^B)_{-j} \cdot \boldsymbol{\gamma}_{\boldsymbol{y}-j} + \boldsymbol{\epsilon}_{\boldsymbol{y}}^B\right)\right) \\
&= \frac{1}{n_B} \cdot \mathrm{Var}\left(g_j\right) \cdot \mathrm{Var}\left(G_{-j}\boldsymbol{\gamma}_{\boldsymbol{y}-j} + \boldsymbol{\epsilon}_{\boldsymbol{y}}^B\right) \\
&= \frac{1-\alpha^2-\gamma_{y_j}^2}{n_B}
\end{aligned} \tag{S7}$$

Expanding the expressions for the $\tau$s, (S3) becomes

$$\begin{aligned}
\tau_{x_j}^A &= \frac{1}{n_A} \cdot \left\{(\boldsymbol{g}_j^{A\setminus B})' \cdot \left((G^{A\setminus B})_{-j}\boldsymbol{\gamma}_{\boldsymbol{x}-j} + \boldsymbol{\epsilon}_{\boldsymbol{x}}^{A\setminus B}\right) + (\boldsymbol{g}_j^{A\cap B})' \cdot \left((G^{A\cap B})_{-j}\boldsymbol{\gamma}_{\boldsymbol{x}-j} + \boldsymbol{\epsilon}_{\boldsymbol{x}}^{A\cap B}\right)\right\} \\
\tau_{x_j}^B &= \frac{1}{n_B} \cdot \left\{(\boldsymbol{g}_j^{B\setminus A})' \cdot \left((G^{B\setminus A})_{-j}\boldsymbol{\gamma}_{\boldsymbol{x}-j} + \boldsymbol{\epsilon}_{\boldsymbol{x}}^{B\setminus A}\right) + (\boldsymbol{g}_j^{A\cap B})' \cdot \left((G^{A\cap B})_{-j}\boldsymbol{\gamma}_{\boldsymbol{x}-j} + \boldsymbol{\epsilon}_{\boldsymbol{x}}^{A\cap B}\right)\right\}
\end{aligned} \tag{S8}$$

$$\tau^B_{y_j} = \frac{1}{n_B} \cdot \left\{ (\boldsymbol{g}_j^{B\setminus A})' \cdot \left( (G^{B\setminus A})_{-j} \boldsymbol{\gamma}_{\boldsymbol{y}-j} + \boldsymbol{\epsilon}_{\boldsymbol{y}}^{B\setminus A} \right) + (\boldsymbol{g}_j^{A\cap B})' \cdot \left( (G^{A\cap B})_{-j} \boldsymbol{\gamma}_{\boldsymbol{y}-j} + \boldsymbol{\epsilon}_{\boldsymbol{y}}^{A\cap B} \right) \right\}$$

Exploiting the fact that the covariance between quantities derived from non-overlapping samples is zero, it enables us to work out the pairwise covariances as follows

$$\begin{aligned}
\mathrm{Cov}\left(\tau^A_{x_j}, \tau^B_{x_j}\right) &= \mathrm{E}\Bigg[ \frac{1}{n_A} \cdot \left\{ (\boldsymbol{g}_j^{A\setminus B})' \cdot \left( (G^{A\setminus B})_{-j} \boldsymbol{\gamma}_{\boldsymbol{x}-j} + \boldsymbol{\epsilon}_{\boldsymbol{x}}^{A\setminus B} \right) + (\boldsymbol{g}_j^{A\cap B})' \cdot \left( (G^{A\cap B})_{-j} \boldsymbol{\gamma}_{\boldsymbol{x}-j} + \boldsymbol{\epsilon}_{\boldsymbol{x}}^{A\cap B} \right) \right\} \times \\
&\quad \frac{1}{n_B} \cdot \left\{ (\boldsymbol{g}_j^{B\setminus A})' \cdot \left( (G^{B\setminus A})_{-j} \boldsymbol{\gamma}_{\boldsymbol{x}-j} + \boldsymbol{\epsilon}_{\boldsymbol{x}}^{B\setminus A} \right) + (\boldsymbol{g}_j^{A\cap B})' \cdot \left( (G^{A\cap B})_{-j} \boldsymbol{\gamma}_{\boldsymbol{x}-j} + \boldsymbol{\epsilon}_{\boldsymbol{x}}^{A\cap B} \right) \right\} \Bigg] \\
&= \frac{1}{n_A \cdot n_B} \cdot \mathrm{E}\left[ \left\{ (\boldsymbol{g}_j^{A\cap B})' \cdot \left( (G^{A\cap B})_{-j} \boldsymbol{\gamma}_{\boldsymbol{x}-j} + \boldsymbol{\epsilon}_{\boldsymbol{x}}^{A\cap B} \right) \right\}^2 \right] \\
&= \frac{n_{A\cap B}}{n_A \cdot n_B} \cdot \mathrm{Var}\left(\boldsymbol{g}_j\right) \cdot \mathrm{Var}\left( (G^{A\cap B})_{-j} \boldsymbol{\gamma}_{\boldsymbol{x}-j} + \boldsymbol{\epsilon}_{\boldsymbol{x}}^{A\cap B} \right) \qquad \text{(S9)} \\
&= \frac{n_{A\cap B}}{n_A \cdot n_B} \cdot (1 - \gamma^2_{x_j})
\end{aligned}$$

Similarly for $\mathrm{Cov}\left(\tau^A_{x_j}, \tau^B_{y_j}\right)$, using the INSIDE assumption, i.e. that $\boldsymbol{\gamma}_{\boldsymbol{x}}$ and $\boldsymbol{\gamma}_{\boldsymbol{y}}$ are uncorrelated, we have

$$\begin{aligned}
\mathrm{Cov}\left(\tau^A_{x_j}, \tau^B_{y_j}\right) &= \mathrm{E}\Bigg[ \frac{1}{n_A} \cdot \left\{ (\boldsymbol{g}_j^{A\setminus B})' \cdot \left( (G^{A\setminus B})_{-j} \boldsymbol{\gamma}_{\boldsymbol{x}-j} + \boldsymbol{\epsilon}_{\boldsymbol{x}}^{A\setminus B} \right) + (\boldsymbol{g}_j^{A\cap B})' \cdot \left( (G^{A\cap B})_{-j} \boldsymbol{\gamma}_{\boldsymbol{x}-j} + \boldsymbol{\epsilon}_{\boldsymbol{x}}^{A\cap B} \right) \right\} \times \\
&\quad \frac{1}{n_B} \cdot \left\{ (\boldsymbol{g}_j^{B\setminus A})' \cdot \left( (G^{B\setminus A})_{-j} \boldsymbol{\gamma}_{\boldsymbol{y}-j} + \boldsymbol{\epsilon}_{\boldsymbol{y}}^{B\setminus A} \right) + (\boldsymbol{g}_j^{A\cap B})' \cdot \left( (G^{A\cap B})_{-j} \boldsymbol{\gamma}_{\boldsymbol{y}-j} + \boldsymbol{\epsilon}_{\boldsymbol{y}}^{A\cap B} \right) \right\} \Bigg] \\
&= \frac{1}{n_A \cdot n_B} \cdot \mathrm{E}\left[ \left\{ (\boldsymbol{g}_j^{A\cap B})' \cdot \left( (G^{A\cap B})_{-j} \boldsymbol{\gamma}_{\boldsymbol{x}-j} + \boldsymbol{\epsilon}_{\boldsymbol{x}}^{A\cap B} \right) \right\} \times \right. \\
&\quad \left. \left\{ (\boldsymbol{g}_j^{A\cap B})' \cdot \left( (G^{A\cap B})_{-j} \boldsymbol{\gamma}_{\boldsymbol{y}-j} + \boldsymbol{\epsilon}_{\boldsymbol{y}}^{A\cap B} \right) \right\} \right] \\
&= \frac{n_{A\cap B}}{n_A \cdot n_B} \cdot \mathrm{Var}\left(\boldsymbol{g}_j\right) \cdot \mathrm{E}\left[ \boldsymbol{\epsilon}_{\boldsymbol{x}} \cdot \boldsymbol{\epsilon}_{\boldsymbol{y}} \right] \qquad \text{(S10)} \\
&= \frac{n_{A\cap B}}{n_A \cdot n_B} \cdot \rho
\end{aligned}$$

We are now in position to compute the covariance between $(\widehat{\beta}^A_x)_j$ and $(\widehat{\beta}^B_y)_j$

$$\begin{aligned}
\mathrm{Cov}\left( (\widehat{\beta}^A_x)_j, (\widehat{\beta}^B_y)_j \right) &= \mathrm{E}\left[ ((\widehat{\beta}^A_x)_j - \gamma_{x_j}) \cdot ((\widehat{\beta}^B_y)_j - \alpha \cdot \gamma_{x_j} - \gamma_{y_j}) \right] \\
&= \mathrm{E}\left[ \tau^A_{x_j} \cdot (\alpha \cdot \tau^B_{x_j} + \tau^B_{y_j}) \right] \\
&= \alpha \cdot \mathrm{Cov}\left(\tau^A_{x_j}, \tau^B_{x_j}\right) + \mathrm{Cov}\left(\tau^A_{x_j}, \tau^B_{y_j}\right) \qquad \text{(S11)} \\
&= \frac{n_{A\cap B}}{n_A \cdot n_B} \cdot (\alpha \cdot (1 - \gamma^2_{x_j}) + \rho)
\end{aligned}$$

Let us now consider the fixed-effect inverse-variance weighting meta-analysis for the ratio estimates for the causal effect $\alpha$. Each IV $j$ provides a ratio estimate

$$\widehat{\alpha}_j = \frac{(\widehat{\beta}^B_y)_j}{(\widehat{\beta}^A_x)_j} \qquad \text{(S12)}$$

$$\mathrm{Var}\left(\widehat{\alpha}_j\right) = \frac{\mathrm{Var}\left( (\widehat{\beta}^B_y)_j \right)}{(\widehat{\beta}^A_x)^2_j} = \frac{(1 - \alpha^2 - \gamma^2_{y_j})}{n_B \cdot (\widehat{\beta}^A_x)^2_j} \qquad \text{(S13)}$$

Hence the weights $(w_j)$ of IV $j$ for estimating the IVW causal effect are

$$w_j \quad = \quad \frac{1}{\mathrm{Var}\left(\widehat{\alpha}_j\right)} = \frac{n_B \cdot (\widehat{\beta}_x^A)_j^2}{(1-\alpha^2-\gamma_{y_j}^2)} \tag{S14}$$

Finally, the estimate can be written in the following form

$$\begin{aligned}\widehat{\alpha}_{IVW} \quad &= \quad \frac{\sum_{k=1}^{m} \widehat{\alpha}_j \cdot w_j}{\sum_{k=1}^{m} w_j} = \frac{\sum_{j=1}^{m} \frac{(\widehat{\beta}_y^B)_j}{(\widehat{\beta}_x^A)_j} \times \left(\frac{(\widehat{\beta}_x^A)_j^2}{(1-\alpha^2-\gamma_{y_j}^2)}\right)}{\sum_{k=1}^{m} \frac{(\widehat{\beta}_x^A)_j^2}{(1-\alpha^2-\gamma_{y_j}^2)}} = \frac{\sum_{k=1}^{m} \frac{(\widehat{\beta}_y^B)_j \cdot (\widehat{\beta}_x^A)_j}{(1-\alpha^2-\gamma_{y_j}^2)}}{\sum_{k=1}^{m} \frac{(\widehat{\beta}_x^A)_j^2}{(1-\alpha^2-\gamma_{y_j}^2)}} \\ &\approx \quad \frac{\sum_{j=1}^{m} \frac{(\widehat{\beta}_y^B)_j \cdot (\widehat{\beta}_x^A)_j}{(1-\alpha^2)}}{\sum_{k=1}^{m} \frac{(\widehat{\beta}_x^A)_j^2}{(1-\alpha^2)}} = \frac{\sum_{k=1}^{m} (\widehat{\beta}_y^B)_j \cdot (\widehat{\beta}_x^A)_j}{\sum_{k=1}^{m} (\widehat{\beta}_x^A)_j^2}\end{aligned} \tag{S15}$$

Here, the last approximation is based on the realistic assumption that the individual pleiotropic effect of each SNP is very small.

#### b. Expectation of the estimate

In reality we select IVs based on their estimated test statistic, typically $|(\widehat{\beta}_x^A)_j| \cdot \sqrt{n_A} > T$ with $T = -\Phi^{-1}(5 \times 10^{-8}/2) \approx 5.45$, representing the genome-wide significance threshold. By denoting $S_j := \left\{|(\widehat{\beta}_x^A)_j| \cdot \sqrt{n_A} > T\right\}$, the causal effect estimate (S15) changes to

$$\widehat{\alpha}_{IVW} \quad \approx \quad \frac{\sum_{j=1}^{M} ((\widehat{\beta}_y^B)_j\,|S_j\,) \cdot ((\widehat{\beta}_x^A)_j\,|S_j\,) \cdot Pr(S_j)}{\sum_{j=1}^{M} ((\widehat{\beta}_x^A)_j\,|S_j\,)^2 \cdot Pr(S_j)} \tag{S16}$$

Note that while $m$ denoted the number of IVs, $M$ represents the number of genome-wide markers from which IVs are selected. By approximating the expectation of a ratio by the ratio of expectations, the expectation of the causal effect estimate (S16) can be written as

$$\mathrm{E}\left[\hat{\alpha}_{IVW}\right] \quad \approx \quad \frac{\sum_{j=1}^{m} \mathrm{E}\left[\left((\alpha \cdot \gamma_{x_j} + \gamma_{y_j}) + \left(\alpha \cdot \tau_{x_j}^B + \tau_{y_j}^B\,|S_j\right)\right) \cdot \left(\gamma_{x_j} + \left(\tau_{x_j}^A\,|S_j\right)\right)\right] \cdot Pr(S_j)}{\sum_{j=1}^{m} \mathrm{E}\left[\left(\gamma_{x_j} + \left(\tau_{x_j}^A\,|S_j\right)\right)^2\right] \cdot Pr(S_j)}$$

$$= \frac{\sum_{j=1}^{m} s_j \cdot Pr(S_j)}{\sum_{j=1}^{m} t_j \cdot Pr(S_j)} \tag{S17}$$

To ensure that the expectation of the ratio could reasonably be approximated by the ratio of expectations in this case, we compared the expectation of the ratio to the ratio of the expectations for both simulated and real data (Figures S17 and S18). We observed a very good agreement between the two and therefore proceeded with this approximation.

Let us expand the $j$th term of the numerator $s_j$

$$\begin{aligned} s_j &= \left(\alpha \cdot \gamma_{x_j}^2 + \gamma_{y_j} \cdot \gamma_{x_j}\right) + \left(\alpha \cdot \gamma_{x_j} + \gamma_{y_j}\right) \cdot \mathrm{E}\left[\tau_{x_j}^A \,|S_j\right] + (\alpha \cdot \gamma_{x_j}) \cdot \mathrm{E}\left[\tau_{x_j}^B \,|S_j\right] \\ &+ \gamma_{x_j} \cdot \mathrm{E}\left[\tau_{y_j}^B \,|S_j\right] + \alpha \cdot \mathrm{E}\left[\tau_{x_j}^B \cdot \tau_{x_j}^A \,|S_j\right] + \mathrm{E}\left[\tau_{y_j}^B \cdot \tau_{x_j}^A \,|S_j\right] \end{aligned} \tag{S18}$$

Similarly for $t_j$ we have

$$t_j = \gamma_{x_j}^2 + 2\gamma_{x_j} \cdot \mathrm{E}\left[\tau_{x_j}^A \,|S_j\right] + \mathrm{E}\left[(\tau_{x_j}^A)^2 \,|S_j\right] \tag{S19}$$

In the following sections we will compute each term of $s_j$ and $t_j$.

- **Switching from** $\mathrm{E}\left[Z = z|S_j\right] \cdot Pr(S_j)$ **to** $\mathrm{E}\left[Z = z|S_j^c\right] \cdot Pr(S_j^c)$

If we define the complement of $S_j$ as $S_j^c := \left\{ -\frac{T}{\sqrt{n_A}} - \gamma_{x_j} \leq \tau_{x_j}^A \leq \frac{T}{\sqrt{n_A}} - \gamma_{x_j} \right\}$, any random variable $Z$ conditional on $S_j$ can be rewritten as follows

$$\begin{aligned} Pr(Z = z|S_j) \cdot Pr(S_j) &= Pr(\{Z = z\} \cap S_j) = Pr(Z = z) - Pr(\{Z = z\} \cap S_j^c) \\ &= Pr(Z = z) - Pr(\{Z = z\}|S_j^c) \cdot Pr(S_j^c) \end{aligned} \tag{S20}$$

Hence:

$$\begin{aligned} \mathrm{E}\left[Z|S_j\right] \cdot Pr(S_j) &= \int_{-\infty}^{\infty} z \cdot Pr(Z = z|S_j) \cdot Pr(S_j) dz \\ &= \int_{-\infty}^{\infty} z \cdot \left(Pr(Z = z) - Pr(\{Z = z\}|S_j^c) \cdot Pr(S_j^c)\right) dz \\ &= \mathrm{E}\left[Z\right] - \mathrm{E}\left[Z|S_j^c\right] \cdot Pr(S_j^c) \end{aligned} \tag{S21}$$

We first calculate $Pr(S_j^c)$ using integration by substitution ($f(t) = t\sqrt{n_A}$), which will be necessary for all further computations

$$\begin{aligned} h_j &:= Pr(S_j^c) = \int_{-\frac{T}{\sqrt{n_A}} - \gamma_{x_j}}^{\frac{T}{\sqrt{n_A}} - \gamma_{x_j}} \frac{1}{\sqrt{2\pi \cdot n_A^{-1}}} \cdot \exp\left(-\frac{u^2}{2 \cdot n_A^{-1}}\right) du \\ &= \Phi(T - \sqrt{n_A}\gamma_{x_j}) - \Phi(-T - \sqrt{n_A}\gamma_{x_j}) \end{aligned} \tag{S22}$$

with $\Phi$ being the standard normal cumulative distribution function. In the following we will substitute $\tau_x^A, \tau_x^B, \tau_y^B, (\tau_x^B \cdot \tau_x^A)$ and $(\tau_y^B \cdot \tau_x^A)$ for $Z$ in (S21) and compute each conditional expectation. Note that when no thresholding is applied ($T = 0$) we have $h_j = 0$.

- **Computation of** $\mathrm{E}\left[\tau_{x_j}^A \,|S_j\right] \cdot Pr(S_j)$

$$\mathrm{E}\left[\tau_{x_j}^A \,|S_j\right] \cdot Pr(S_j) \;=\; \left(\mathrm{E}\left[\tau_{x_j}^A\right] - \mathrm{E}\left[\tau_{x_j}^A|S_j^c\right] \cdot h_j\right) = -h_j \cdot \mathrm{E}\left[\tau_{x_j}^A|S_j^c\right] \tag{S23}$$

Using the properties of the truncated normal distribution we get

$$\begin{aligned}\mathrm{E}\left[\tau_{x_j}^A \,|S_j\right] \cdot Pr(S_j) &= -h \cdot \frac{1}{\sqrt{n_A}} \cdot \frac{\phi(-T-\gamma_{x_j}\sqrt{n_A}) - \phi(T-\gamma_{x_j}\sqrt{n_A}))}{\Phi(T-\gamma_{x_j}\sqrt{n_A}) - \Phi(-T-\gamma_{x_j}\sqrt{n_A})} \\ &= \frac{1}{\sqrt{n_A}} \cdot (\phi(T-\gamma_{x_j}\sqrt{n_A}) - \phi(-T-\gamma_{x_j}\sqrt{n_A}))\end{aligned} \tag{S24}$$

with $\phi$ being the standard normal probability density function.

- **Computation of** $\mathrm{E}\left[(\tau_{x_j}^A)^2 \,|S_j\right] \cdot Pr(S_j)$

$$\mathrm{E}\left[(\tau_{x_j}^A)^2 \,|S_j\right] \cdot Pr(S_j) \;=\; \left(\mathrm{E}\left[\tau_{x_j}^A)^2\right] - \mathrm{E}\left[(\tau_{x_j}^A)^2|S_j^c\right] \cdot h_j\right) = \frac{1}{n_A} - h \cdot \mathrm{E}\left[(\tau_{x_j}^A)^2|S_j^c\right] \tag{S25}$$

Using the properties of the truncated normal distribution we get

$$\begin{aligned}&\mathrm{E}\left[(\tau_{x_j}^A)^2 \,|S_j\right] \cdot Pr(S_j) = \\ &= \frac{1}{n_A} - h_j \cdot \frac{1}{n_A} \cdot \left(1 + \frac{(-T-\gamma_{x_j}\sqrt{n_A}) \cdot \phi(-T-\gamma_{x_j}\sqrt{n_A}) - (T-\gamma_{x_j}\sqrt{n_A}) \cdot \phi(T-\gamma_{x_j}\sqrt{n_A}))}{\Phi(T-\gamma_{x_j}\sqrt{n_A}) - \Phi(-T-\gamma_{x_j}\sqrt{n_A})}\right) \\ &= \frac{1}{n_A}(1-h_j) - \frac{1}{n_A} \cdot \left((-T-\gamma_{x_j}\sqrt{n_A}) \cdot \phi(-T-\gamma_{x_j}\sqrt{n_A}) - (T-\gamma_{x_j}\sqrt{n_A}) \cdot \phi(T-\gamma_{x_j}\sqrt{n_A}))\right)\end{aligned} \tag{S26}$$

- **Computation of** $\mathrm{E}\left[\tau_{x_j}^B \,|S_j\right] \cdot Pr(S_j)$

We can split the error term $\tau_{x_j}^B$ into $\tau_{x_j}^A$ dependent and independent parts. We have shown above that $\mathrm{Cov}\left(\tau_{x_j}^A, \tau_{x_j}^B\right) = (1-\gamma_{x_j}^2) \cdot n_{A\cap B}/(n_A \cdot n_B)$ and $\mathrm{Var}\left(\tau_{x_j}^A\right) = (1-\gamma_{x_j}^2)/n_A$, thus we have

$$\begin{aligned}\tau_{x_j}^B &= \tau_{x_j}^A \cdot \frac{\mathrm{Cov}\left(\tau_{x_j}^A, \tau_{x_j}^B\right)}{\mathrm{Var}\left(\tau_{x_j}^A\right)} + \eta_x^B \quad \text{with} \quad \mathrm{Cov}\left(\eta_x^B, \tau_{x_j}^A\right) = 0 \\ &= \tau_{x_j}^A \cdot \frac{n_{A\cap B}}{n_B} + \eta_x^B\end{aligned} \tag{S27}$$

This allows us to utilise the formula for $\mathrm{E}\left[\tau_{x_j}^A \,|S_j\right] \cdot Pr(S_j)$ to derive $\mathrm{E}\left[\tau_{x_j}^B \,|S_j\right] \cdot Pr(S_j)$ as follows

$$\mathrm{E}\left[\tau_{x_j}^B \,|S_j\right] \cdot Pr(S_j) \;=\; \frac{n_{A\cap B}}{n_B} \cdot \mathrm{E}\left[\tau_{x_j}^A \,|S\right] \cdot Pr(S_j) + \mathrm{E}\left[\eta_x^B \,|S_j\right]$$

$$= \frac{n_{A\cap B}}{n_B} \cdot \mathrm{E}\left[\tau^A_{x_j} \,|S_j\right] \cdot Pr(S_j) \tag{S28}$$

- **Computation of** $\mathrm{E}\left[\tau^B_{x_j} \cdot \tau^A_{x_j} \,|S\right] \cdot Pr(S_j)$

Using the split sample notation helps us to compute the expectation of the $\left(\tau^B_{x_j} \cdot \tau^A_{x_j} \,|S_j\right)$ term analogously to how we did for $\left(\tau^B_{x_j} \,|S_j\right)$

$$\begin{aligned}
\mathrm{E}\left[\tau^B_{x_j} \cdot \tau^A_{x_j} \,|S_j\right] \cdot Pr(S_j) &= \mathrm{E}\left[(\tau^A_{x_j} \cdot \frac{n_{A\cap B}}{n_B} + \eta^B_x) \cdot \tau^A_{x_j} \,|S_j\right] \cdot Pr(S_j) \\
&= \frac{n_{A\cap B}}{n_B} \mathrm{E}\left[(\tau^A_{x_j})^2 \,|S_j\right] \cdot Pr(S_j) \qquad \text{(S29)}
\end{aligned}$$

- **Computation of** $\mathrm{E}\left[\tau^B_{y_j} \,|S_j\right] \cdot Pr(S_j)$ **and** $\mathrm{E}\left[\tau^B_{y_j} \cdot \tau^A_{x_j} \,|S_j\right] \cdot Pr(S_j)$

The decomposing $\tau^B_{y_j} = \rho\tau^B_{x_j} + \eta^B$ with $\mathrm{Cov}\left(\eta^B, \tau^B_{x_j}\right) = 0$ allows us to trace back this computation to that of $\mathrm{E}\left[\tau^B_{x_j} \,|S_j\right]$ and $\mathrm{E}\left[\tau^A_{x_j} \cdot \tau^B_{x_j} \,|S_j\right]$, respectively

$$\mathrm{E}\left[\tau^B_{y_j} \,|S_j\right] \cdot Pr(S_j) = \rho \cdot \mathrm{E}\left[\tau^B_{x_j} \,|S_j\right] \cdot Pr(S_j) \tag{S30}$$

$$\mathrm{E}\left[\tau^B_{y_j} \cdot \tau^A_{x_j} \,|S_j\right] \cdot Pr(S_j) = \rho \cdot \mathrm{E}\left[\tau^B_{x_j} \cdot \tau^A_{x_j} \,|S_j\right] \cdot Pr(S_j) \tag{S31}$$

- **Evaluation of** $s_j$

We are now in position to evaluate the expression for $s_j$

$$\begin{aligned}
s_j &= \left(\alpha \cdot \gamma^2_{x_j} + \gamma_{y_j} \cdot \gamma_{x_j}\right) + \left(\alpha \cdot \gamma_{x_j} + \gamma_{y_j}\right) \cdot \mathrm{E}\left[\tau^A_{x_j} \,|S_j\right] + (\alpha \cdot \gamma_{x_j}) \cdot \mathrm{E}\left[\tau^B_{x_j} \,|S_j\right] + \gamma_{x_j} \cdot \mathrm{E}\left[\tau^B_{y_j} \,|S_j\right] \\
&+ \alpha \cdot \mathrm{E}\left[\tau^B_{x_j} \cdot \tau^A_{x_j} \,|S_j\right] + \mathrm{E}\left[\tau^B_{y_j} \cdot \tau^A_{x_j} \,|S_j\right] \\
&= \left(\alpha \cdot \gamma^2_{x_j} + \gamma_{y_j} \cdot \gamma_{x_j}\right) + \left(\alpha \cdot \gamma_{x_j} + \gamma_{y_j}\right) \cdot \mathrm{E}\left[\tau^A_{x_j} \,|S_j\right] + (\alpha \cdot \gamma_{x_j}) \cdot \frac{n_{A\cap B}}{n_B} \cdot \mathrm{E}\left[\tau^A_{x_j} \,|S_j\right] \\
&+ \gamma_{x_j} \cdot \rho \cdot \frac{n_{A\cap B}}{n_B} \cdot \mathrm{E}\left[\tau^A_{x_j} \,|S_j\right] + \alpha \cdot \frac{n_{A\cap B}}{n_B} \cdot \mathrm{E}\left[(\tau^A_{x_j})^2 \,|S_j\right] + \rho \cdot \frac{n_{A\cap B}}{n_B} \cdot \mathrm{E}\left[(\tau^A_{x_j})^2 \,|S_j\right] \qquad \text{(S32)} \\
&= \left(\alpha \cdot \gamma^2_{x_j} + \gamma_{y_j} \cdot \gamma_{x_j}\right) + \left(\alpha \cdot \gamma_{x_j} + \gamma_{y_j} + (\alpha \cdot \gamma_{x_j}) \cdot \frac{n_{A\cap B}}{n_B} + \gamma_{x_j} \cdot \rho \cdot \frac{n_{A\cap B}}{n_B}\right) \cdot \mathrm{E}\left[\tau^A_{x_j} \,|S_j\right] \\
&+ (\alpha + \rho) \cdot \frac{n_{A\cap B}}{n_B} \cdot \mathrm{E}\left[(\tau^A_{x_j})^2 \,|S_j\right]
\end{aligned}$$

Let $\lambda'$ denote a quantity closely related to the cross-trait LD score regression (LDSC) [2] intercept $(\lambda)$

$$\lambda' = (\alpha + \rho) \cdot \frac{n_{A\cap B}}{n_A \cdot n_B} = \frac{\lambda}{\sqrt{n_A \cdot n_B}} \tag{S33}$$

The expression for $s_j$ (S32) can then be turned into

$$s_j \cdot Pr(S_j) = \left(\alpha \cdot \gamma^2_{x_j} + \gamma_{y_j} \cdot \gamma_{x_j}\right) \cdot (1 - h_j) + \left(\alpha \cdot \gamma_{x_j} + \gamma_{y_j} + \gamma_{x_j} \cdot \lambda' \cdot n_A\right) \cdot \mathrm{E}\left[\tau^A_{x_j} \,|S_j\right] \cdot Pr(S_j)$$

$$
\begin{aligned}
&+ \quad \lambda' \cdot n_A \cdot \mathrm{E}\left[(\tau_{x_j}^A)^2 \,|S_j\right] \cdot Pr(S_j) \\
&= \quad \left(\alpha \cdot \gamma_{x_j}^2 + \gamma_{y_j} \cdot \gamma_{x_j}\right) \cdot (1-h_j) \\
&+ \quad \left(\alpha \cdot \gamma_{x_j} + \gamma_{y_j} + \gamma_{x_j} \cdot \lambda' \cdot n_A\right) \cdot \frac{1}{\sqrt{n_A}} \cdot \left(\phi(T - \gamma_{x_j}\sqrt{n_A}) - \phi(-T - \gamma_{x_j}\sqrt{n_A})\right) \\
&+ \quad \lambda' \cdot n_A \cdot \frac{1}{n_A}(1-h_j) \qquad \text{(S34)} \\
&+ \quad \lambda' \cdot n_A \cdot \frac{1}{n_A} \cdot \left((T - \gamma_{x_j}\sqrt{n_A}) \cdot \phi(T - \gamma_{x_j}\sqrt{n_A}) - (-T - \gamma_{x_j}\sqrt{n_A}) \cdot \phi(-T - \gamma_{x_j}\sqrt{n_A})\right) \\
&= \quad \left(\alpha \cdot \gamma_{x_j}^2 + \gamma_{y_j} \cdot \gamma_{x_j}\right) \cdot (1-h_j) \\
&+ \quad \left(\frac{\alpha\gamma_{x_j} + \gamma_{y_j} + \lambda' \cdot \sqrt{n_A} \cdot T}{\sqrt{n_A}}\right) \cdot \phi(T - \gamma_{x_j} \cdot \sqrt{n_A}) \\
&- \quad \left(\frac{\alpha\gamma_{x_j} + \gamma_{y_j} - \lambda' \cdot \sqrt{n_A} \cdot T}{\sqrt{n_A}}\right) \cdot \phi(-T - \gamma_{x_j} \cdot \sqrt{n_A}) \\
&+ \quad \lambda'(1-h_j)
\end{aligned}
$$

In the following, we will assume the following popular [3] genetic architecture for $X$ :

$$
\boldsymbol{\gamma_x} \quad = \quad \boldsymbol{\zeta_x} \odot \boldsymbol{v_x} \qquad \text{with} \quad \boldsymbol{\zeta_x} \sim \mathcal{B}(1, \pi_x) \quad \text{and} \quad \boldsymbol{v_x} \sim \mathcal{N}(0, \sigma_x^2) \qquad \text{(S35)}
$$

where $\pi_x$ and $\sigma_x^2$ are characteristics of the genetic architecture of trait $X$ (respectively, a measure of the polygenicity and the per-variant heritability).
It allows us to split the sums for zero effect genetic variants and non-zero effect genetic variants.
When $\gamma_{x_j} = 0$ the expression simplifies to

$$
\sum_{j \in \mathcal{M}_0} s_j \cdot Pr(S_j) \quad = \quad M \cdot (1 - \pi_x) \cdot 2\lambda' \cdot (T \cdot \phi(T) + \Phi(-T)) \qquad \text{(S36)}
$$

For non-zero effects we have:

$$
\begin{aligned}
\frac{\sum_{j \in \mathcal{M}_1} s_j \cdot Pr(S_j)}{\pi_x \cdot M} \quad &= \quad \int_{-\infty}^{\infty} \frac{1}{\sigma_x} \cdot \phi(\gamma_x/\sigma_x) \cdot \left(\alpha \cdot \gamma_x^2 + \gamma_{y_j} \cdot \gamma_x + \lambda'\right) \\
&\times \quad (1 - \Phi(T - \sqrt{n_A}\gamma_x) + \Phi(-T - \sqrt{n_A}\gamma_x))\, d\gamma_x \\
&+ \quad \int_{-\infty}^{\infty} \frac{1}{\sigma_x} \cdot \phi(\gamma_x/\sigma_x) \\
&\times \quad \left[\left(\frac{\alpha\gamma_x + \gamma + \lambda'\sqrt{n_A}T}{\sqrt{n_A}}\right) \cdot \phi(T - \gamma_x\sqrt{n_A})\right. \\
&- \quad \left.\left(\frac{\alpha\gamma_x + \gamma - \lambda'\sqrt{n_A}T}{\sqrt{n_A}}\right) \cdot \phi(-T - \gamma_x\sqrt{n_A})\right] d\gamma_x \\
&= \quad 2 \cdot \int_{\infty}^{\infty} \left(\alpha \cdot \gamma_x^2 + \lambda'\right) \cdot \frac{1}{\sigma_x} \cdot \phi(\gamma_x/\sigma_x) \cdot \Phi(-T + \sqrt{n_A}\gamma_x) d\gamma_x \\
&+ \quad \frac{\exp\left(-\frac{1}{2} \cdot \frac{T^2}{1 + \sigma_x^2 \cdot n_A}\right) \cdot \sqrt{\frac{2}{\pi}} \cdot T \cdot (\alpha \cdot \sigma_x^2 + \lambda' \cdot (1 + \sigma_x^2 \cdot n_A))}{(1 + \sigma_x^2 \cdot n_A)^{(3/2)}} \qquad \text{(S37)}
\end{aligned}
$$

$$
\begin{aligned}
&= 2\lambda' \cdot \Phi\left(-\frac{T}{\sqrt{1+n_A\cdot\sigma_x^2}}\right) \\
&+ 2\alpha \cdot \int_{\infty}^{\infty} \gamma_x^2 \cdot \frac{1}{\sigma_x} \cdot \phi(\gamma_x/\sigma_x) \cdot \Phi(-T+\sqrt{n_A}\gamma_x) d\gamma_x \\
&+ \frac{\exp\left(-\frac{1}{2}\cdot\frac{T^2}{1+\sigma_x^2\cdot n_A}\right)\cdot\sqrt{\frac{2}{\pi}}\cdot T\cdot(\alpha\cdot\sigma_x^2+\lambda'\cdot(1+\sigma_x^2\cdot n_A))}{(1+\sigma_x^2\cdot n_A)^{(3/2)}}
\end{aligned}
$$

The remaining integral can be solved as follows

$$
\begin{aligned}
\frac{\partial}{\partial x}\left(x\cdot\frac{\phi(x/c)}{c}\cdot\Phi(a+b\cdot x)\right) &= \frac{\phi(x/c)}{c}\cdot\Phi(a+b\cdot x) + x\cdot\left(\frac{1}{c}\cdot\frac{-x}{c^2}\cdot\phi(x/c)\right)\cdot\Phi(a+b\cdot x) \\
&+ x\cdot\frac{\phi(x/c)}{c}\cdot(b\cdot\phi(a+b\cdot x))
\end{aligned}
\tag{S38}
$$

By integrating both sides w.r.t. $x$ from $-\infty$ to $\infty$ we get

$$
\begin{aligned}
\left[x\cdot\frac{\phi(x/c)}{c}\cdot\Phi(a+b\cdot x)\right]_{-\infty}^{\infty} &= \int_{-\infty}^{\infty}\frac{\phi(x/c)}{c}\cdot\Phi(a+b\cdot x)dx \\
&+ \int_{-\infty}^{\infty} x\cdot\left(\frac{1}{c}\cdot\frac{-x}{c^2}\cdot\phi(x/c)\right)\cdot\Phi(a+b\cdot x)dx \\
&+ \int_{-\infty}^{\infty} x\cdot\frac{\phi(x/c)}{c}\cdot(b\cdot\phi(a+b\cdot x))\,dx
\end{aligned}
\tag{S39}
$$

It is easy to see that the quantity on the left hand side is zero (both limits are zero). We use the following two well-known integral identities

$$
\int_{-\infty}^{\infty}\phi(x)\cdot\Phi(a+b\cdot x)dx = \Phi\left(\frac{a}{\sqrt{1+b^2}}\right) \tag{S40}
$$

$$
\int_{-\infty}^{\infty} x\cdot\phi(x)\cdot\phi(a+b\cdot x)dx = -\frac{ab\cdot\exp\left(-\frac{a^2}{2(b^2+1)}\right)}{\sqrt{2\pi}(1+b^2)^{3/2}} \tag{S41}
$$

Therefore the above equation (S39) simplifies to

$$
\begin{aligned}
0 &= \Phi\left(\frac{a}{\sqrt{1+(bc)^2}}\right) \\
&- \frac{1}{c^3}\cdot\int_{-\infty}^{\infty} x^2\cdot\phi(x/c)\cdot\Phi(a+b\cdot x)dx \\
&- (bc)\cdot\frac{abc\cdot\exp\left(-\frac{a^2}{2((bc)^2+1)}\right)}{\sqrt{2\pi}(1+(bc)^2)^{3/2}}
\end{aligned}
\tag{S42}
$$

Thus we have

$$
\int_{-\infty}^{\infty} x^2\cdot\frac{\phi(x/c)}{c}\cdot\Phi(a+b\cdot x)dx = c^2\cdot\Phi\left(\frac{a}{\sqrt{1+(bc)^2}}\right)
$$

$$
- \quad a \cdot b^2 \cdot c^4 \cdot \frac{\exp\left(-\frac{a^2}{2((bc)^2+1)}\right)}{\sqrt{2\pi}(1+(bc)^2)^{3/2}} \tag{S43}
$$

Substituting $a = -T, b = \sqrt{n_A}, c = \sigma_x$ yields:

$$
\begin{aligned}
\int_{-\infty}^{\infty} x^2 \cdot \frac{\phi(x/\sigma_x)}{\sigma_x} \cdot \Phi(-T + \sqrt{n_A} \cdot x)dx &= \sigma_x^2 \cdot \Phi\left(\frac{-T}{\sqrt{1+n_A\sigma_x^2}}\right) \\
&+ T \cdot n_A \cdot \sigma_x^4 \cdot \frac{\exp\left(-\frac{T^2}{2(n_A\sigma_x^2+1)}\right)}{\sqrt{2\pi}(1+n_A\sigma_x^2)^{3/2}}
\end{aligned} \tag{S44}
$$

Finally, we can provide a closed form expression for $\sum_j s_j \cdot Pr(S_j)$ as follows

$$
\begin{aligned}
\frac{\sum\limits_{j\in\mathcal{M}_1} s_j \cdot Pr(S_j)}{\pi_x \cdot M} &= 2\lambda' \cdot \Phi\left(-\frac{T}{\sqrt{1+n_A \cdot \sigma_x^2}}\right) \\
&+ 2\alpha \cdot \left(\sigma_x^2 \cdot \Phi\left(\frac{-T}{\sqrt{1+n_A\sigma_x^2}}\right) + T \cdot n_A \cdot \sigma_x^4 \cdot \frac{\exp\left(-\frac{T^2}{2(n_A\sigma_x^2+1)}\right)}{\sqrt{2\pi}(1+n_A\sigma_x^2)^{3/2}}\right) \\
&+ \frac{\exp\left(-\frac{1}{2}\cdot\frac{T^2}{1+\sigma_x^2 \cdot n_A}\right) \cdot \sqrt{\frac{2}{\pi}} \cdot T \cdot (\alpha \cdot \sigma_x^2 + \lambda' \cdot (1+\sigma_x^2 \cdot n_A))}{(1+\sigma_x^2 \cdot n_A)^{(3/2)}} \qquad \text{(S45)} \\
&= 2(\alpha\sigma_x^2 + \lambda') \cdot \Phi\left(-\frac{T}{\sqrt{1+n_A \cdot \sigma_x^2}}\right) \\
&+ 2T \cdot \frac{\exp\left(-\frac{T^2}{2(n_A\sigma_x^2+1)}\right)}{\sqrt{2\pi}(1+n_A\sigma_x^2)^{3/2}} \cdot \left(\alpha \cdot n_A \cdot \sigma_x^4 + \alpha\sigma_x^2 + \lambda' \cdot (1+\sigma_x^2 \cdot n_A)\right)
\end{aligned}
$$

- **Evaluation of $t_j$**

Similarly, we can compute $t_j \cdot Pr(S_j)$

$$
\begin{aligned}
t_j \cdot Pr(S_j) &= \gamma_{x_j}^2 \cdot (1-h) + 2\gamma_{x_j} \cdot \mathrm{E}\left[\tau_{x_j}^A \,|S_j\right] \cdot Pr(S_j) + \mathrm{E}\left[(\tau_{x_j}^A)^2 \,|S_j\right] \cdot Pr(S_j) \\
&= \gamma_{x_j}^2 \cdot (1-h) + 2\gamma_{x_j} \cdot \frac{1}{\sqrt{n_A}} \cdot (\phi(T - \gamma_{x_j}\sqrt{n_A}) - \phi(-T - \gamma_{x_j}\sqrt{n_A})) \\
&+ \frac{1}{n_A}(1-h) - \frac{1}{n_A} \cdot \big((-T - \gamma_{x_j}\sqrt{n_A}) \cdot \phi(-T - \gamma_{x_j}\sqrt{n_A}) \qquad \text{(S46)} \\
&- (T - \gamma_{x_j}\sqrt{n_A}) \cdot \phi(T - \gamma_{x_j}\sqrt{n_A})\big) \\
&= \left(\gamma_{x_j}^2 + \frac{1}{n_A}\right) \cdot (1-h) \\
&+ \frac{1}{n_A} \cdot \big((T + \gamma_{x_j}\sqrt{n_A}) \cdot \phi(T - \gamma_{x_j}\sqrt{n_A}) + (T - \gamma_{x_j}\sqrt{n_A}) \cdot \phi(-T - \gamma_{x_j}\sqrt{n_A})\big)
\end{aligned}
$$

As before, when $\gamma_{x_j} = 0$ the expression simplifies to

$$\sum_{j \in \mathcal{M}_0} t_j \cdot Pr(S_j) \quad = \quad M \cdot (1 - \pi_x) \cdot \frac{2}{n_A} \cdot (\Phi(-T) + T \cdot \phi(T)) \tag{S47}$$

For non-zero effects we have

$$\begin{aligned}
\frac{\sum_{j \in \mathcal{M}_1} t_j \cdot Pr(S_j)}{\pi_x \cdot M} &= \int_{\infty}^{\infty} (t|\gamma_x) \cdot \frac{1}{\sigma_x} \cdot \phi(\gamma_x/\sigma_x) d\gamma_x \\
&= \int_{\infty}^{\infty} \frac{1}{\sigma_x} \cdot \phi(\gamma_x/\sigma_x) \cdot \left(\gamma_x^2 + \frac{1}{n_A}\right) \cdot \left(1 - \Phi(T - \sqrt{n_A}\gamma_x) + \Phi(-T - \sqrt{n_A}\gamma_x)\right) d\gamma_x \\
&+ \frac{1}{n_A} \cdot \int_{\infty}^{\infty} \left((T + \gamma_{x_j}\sqrt{n_A}) \cdot \phi(T - \gamma_{x_j}\sqrt{n_A}) + (T - \gamma_{x_j}\sqrt{n_A}) \cdot \phi(-T - \gamma_{x_j}\sqrt{n_A})\right) \\
&\times \frac{1}{\sigma_x} \cdot \phi(\gamma_x/\sigma_x) d\gamma_x \\
&= 2 \cdot \int_{\infty}^{\infty} \frac{1}{\sigma_x} \cdot \phi(\gamma_x/\sigma_x) \cdot \left(\gamma_x^2 + \frac{1}{n_A}\right) \cdot \Phi(-T + \sqrt{n_A}\gamma_x) d\gamma_x \\
&+ \frac{1}{n_A} \cdot \frac{\exp\left(-\frac{1}{2} \cdot \frac{T^2}{1 + \sigma_x^2 \cdot n_A}\right) \cdot \sqrt{\frac{2}{\pi}} \cdot T \cdot (2\sigma_x^2 \cdot n_A + 1)}{(1 + \sigma_x^2 \cdot n_A)^{(3/2)}} \\
&= 2 \cdot \frac{1}{n_A} \cdot \Phi\left(-\frac{T}{\sqrt{1 + n_A \cdot \sigma_x^2}}\right) \\
&+ 2 \cdot \int_{\infty}^{\infty} \frac{1}{\sigma_x} \cdot \phi(\gamma_x/\sigma_x) \cdot \gamma_x^2 \cdot \Phi(-T + \sqrt{n_A}\gamma_x) d\gamma_x \\
&+ \frac{1}{n_A} \cdot \frac{\exp\left(-\frac{1}{2} \cdot \frac{T^2}{1 + \sigma_x^2 \cdot n_A}\right) \cdot \sqrt{\frac{2}{\pi}} \cdot T \cdot (2\sigma_x^2 \cdot n_A + 1)}{(1 + \sigma_x^2 \cdot n_A)^{(3/2)}}
\end{aligned} \tag{S48}$$

Plugging in the expression for the integral we already computed for $s_j$ (S44) we have

$$\begin{aligned}
\frac{\sum_{j \in \mathcal{M}_1} t_j \cdot Pr(S_j)}{\pi_x \cdot M} &= 2 \cdot \frac{1}{n_A} \cdot \Phi\left(-\frac{T}{\sqrt{1 + n_A \cdot \sigma_x^2}}\right) \\
&+ 2 \cdot \left(\sigma_x^2 \cdot \Phi\left(\frac{-T}{\sqrt{1 + n_A\sigma_x^2}}\right) + T \cdot n_A \cdot \sigma_x^4 \cdot \frac{\exp\left(-\frac{T^2}{2(n_A\sigma_x^2 + 1)}\right)}{\sqrt{2\pi}(1 + n_A\sigma_x^2)^{3/2}}\right) \\
&+ \frac{1}{n_A} \cdot \frac{\exp\left(-\frac{1}{2} \cdot \frac{T^2}{1 + \sigma_x^2 \cdot n_A}\right) \cdot \sqrt{\frac{2}{\pi}} \cdot T \cdot (2\sigma_x^2 \cdot n_A + 1)}{(1 + \sigma_x^2 \cdot n_A)^{(3/2)}} \\
&= 2 \cdot \left(\sigma_x^2 + \frac{1}{n_A}\right) \cdot \Phi\left(-\frac{T}{\sqrt{1 + n_A \cdot \sigma_x^2}}\right)
\end{aligned} \tag{S49}$$

$$+ \quad 2T \cdot \left( n_A \cdot \sigma_x^4 + 2\sigma_x^2 + \frac{1}{n_A} \right) \cdot \frac{\exp\left( -\frac{T^2}{2(n_A\sigma_x^2+1)} \right)}{\sqrt{2\pi}(1+n_A\sigma_x^2)^{3/2}}$$

- **The final formula**

To simplify the notations, let's define

$$\begin{aligned} a &= \Phi\left( -\frac{T}{\sqrt{1+n_A\cdot\sigma_x^2}} \right) \\ b &= 2T \cdot \frac{\exp\left( -\frac{T^2}{2(n_A\sigma_x^2+1)} \right)}{\sqrt{2\pi}(1+n_A\sigma_x^2)^{3/2}} \\ c &= \Phi(-T) + T\cdot\phi(T) \end{aligned} \tag{S50}$$

This leads us to the expectation of the causal effect estimation, $E[\hat{\alpha}_{IVW}]$

$$\begin{aligned} & \frac{\pi_x \cdot \left(2(\alpha\sigma_x^2+\lambda')\cdot a + b\cdot\left(\alpha\cdot n_A\cdot\sigma_x^4 + \alpha\sigma_x^2 + \lambda'\cdot(1+\sigma_x^2\cdot n_A)\right)\right) + (1-\pi_x)\cdot 2\lambda'\cdot c}{\pi_x\cdot\left(2\cdot\left(\sigma_x^2+\frac{1}{n_A}\right)\cdot a + b\cdot\left(n_A\cdot\sigma_x^4+2\sigma_x^2+\frac{1}{n_A}\right)\right) + (1-\pi_x)\cdot\frac{2}{n_A}\cdot c} \\ = \; & \alpha\cdot\frac{\left(\pi_x\cdot\sigma_x^2\right)\cdot\left(2\cdot a + b\cdot(1+n_A\cdot\sigma_x^2)\right)}{d(n_A,T,\pi_X,\sigma_x^2)} \\ & +\lambda'\cdot\frac{\pi_x\cdot\left(2\cdot a+b\cdot(1+n_A\cdot\sigma_x^2)\right) + \left(1-\pi_x\right)\cdot 2\cdot c}{d(n_A,T,\pi_x,\sigma_x^2)} \end{aligned} \tag{S51}$$

with

$$\begin{aligned} d(n_A,T,\pi_x,\sigma_x^2) &= \pi_x\cdot\left(2\cdot\left(\sigma_x^2+\frac{1}{n_A}\right)\cdot a + b\cdot\left(n_A\cdot\sigma_x^4+2\sigma_x^2+\frac{1}{n_A}\right)\right) \\ & \quad +(1-\pi_x)\cdot\frac{2}{n_A}\cdot c \end{aligned} \tag{S52}$$

To estimate this expectation, one needs to first obtain the per-(active)variant-heritability ($\sigma_x^2$) and polygenicity ($\pi_x$) of the exposure (See Supplementary section B). Note that the exposure heritability is simply $\pi_x \cdot M \cdot \sigma_x^2$. Then a cross-trait LDSC would inform us about the value of $\lambda'$. Finally, the threshold $T$ is decided in the MR analysis.

#### c. Estimation of the corrected effect

From (S51) , we can derive a corrected effect ($\hat{\alpha}_c$) for the causal effect

$$\hat{\alpha}_c = \frac{\hat{\alpha}_{IVW}\cdot d(n_A,T,\pi_X,\sigma_X^2) - \lambda'\cdot\left(\pi_X\cdot\left(2\cdot a+b\cdot(1+\sigma_X^2\cdot n_A)\right) + \left(1-\pi_X\right)\cdot 2\cdot c\right)}{\left(\pi_X\cdot\sigma_X^2\right)\cdot\left(2\cdot a+b\cdot(1+n_A\cdot\sigma_X^2)\right)} \tag{S53}$$

While $n_A, n_B$ and $T$ are known parameters, several other quantities in the correction needs to be estimated in the above equation. The LDSC intercept can be obtained via LD score regression, giving us $\widehat{\lambda}$. The estimation of the two remaining parameters $\widehat{\pi}_x$ and $\widehat{\sigma}_x^2$ are described below.

#### B. Estimation of the genetic architecture

Parameters describing the genetic architecture of the exposure, the per-(active)variant-heritability ($\sigma_x^2$) and polygenicity ($\pi_x$), are needed to estimate the corrected effect $\hat{\alpha}_c$ (S53). To estimate those, we are taking advantage of the equivalence between (S15) and (S17). The denominators of these two formulas are equal, leading us to

$$\begin{aligned} \sum_{j=1}^{m} (\widehat{\beta}_x^A)_j^2 &= \sum_{j=1}^{m} t_j \cdot Pr(S_j) \\ \sum_{k=1}^{m} (\widehat{\beta}_x^A)_k^2 &= M \cdot d(n_A, T, \pi_X, \sigma_X^2) \end{aligned} \quad \text{(S54)}$$

The left-hand side of (S54) can be estimated from the data, and $n_A$ and $T$ are known. By plugging in the heritability estimate from LDSC regression ($\widehat{h}_x^2$) and defining $\sigma_x^2 = \frac{h_x^2}{M \cdot \pi_x}$, the only unknown parameter is $\pi_x$. We can then use an optimisation approach to minimise the difference between $\sum_{k=1}^{m} (\widehat{\beta}_x^A)_k^2$ and $M \cdot d(n_A, T, \pi_x, \sigma_x^2)$ in order to estimate $\pi_x$ and the corresponding $\sigma_x^2$. These estimates, $\widehat{\pi}_x$ and $\widehat{\sigma}_x^2$ can then be plugged into (S53) estimate the corrected effects.

#### C. Estimation of the standard error of the corrected effect estimate and of the covariance between IVW-based and corrected effects

We chose a sampling strategy (parametric bootstrap) to estimate the variance of the corrected causal effect, computed according to (S53). As a first step, we simulate each parameter included in the formula $s$ times. $n_A$, $n_B$, $T$ are known and can be directly used. We start by simulating $s$ IVW-based effects to account for the variability of $\hat{\alpha}_{IVW}$

$$\hat{\boldsymbol{\alpha}}_{1...s} \sim \mathcal{N}\big(\hat{\alpha}_{IVW}, \mathrm{Var}\left(\hat{\alpha}_{IVW}\right)\big) \tag{S55}$$

We also simulate $s$ cross-trait LDsc intercepts and heritability estimates

$$\widehat{\boldsymbol{\lambda}}_{1...s} \sim \mathcal{N}\big(\widehat{\lambda}, \mathrm{Var}\left(\widehat{\lambda}\right)\big) \tag{S56}$$

$$\left(\widehat{\boldsymbol{h}^{\mathbf{2}}_{\boldsymbol{x}}}\right)_{1...s} \sim \mathcal{N}\big(\widehat{h^2_x}, \mathrm{Var}\left(h^2_x\right)\big) \tag{S57}$$

To account for the variability of $\widehat{\pi}_x$ we simulate $s$ set of IVs. Let us consider $\hat{\boldsymbol{\beta}}^{\boldsymbol{A}}_{\boldsymbol{x}}$ the IVW-based effects of the $m$ IVs on the exposure. The effects of the $s$ set of IVs on $X$ are simulated using a multivariate normal distribution

$$\left(\hat{\boldsymbol{\beta}^{\boldsymbol{A}}_{\boldsymbol{x}}}\right)_{1...s} \sim \mathcal{N}\left(\hat{\boldsymbol{\beta}}^{\boldsymbol{A}}_{\boldsymbol{x}}, \frac{I}{n_A}\right) \tag{S58}$$

For each set of IVs, we use the approach described in Supplementary section B to estimate $(\widehat{\boldsymbol{\pi}}_{\boldsymbol{x}})_{1...s}$ and $\left(\widehat{\boldsymbol{\sigma}}^{\mathbf{2}}_{\boldsymbol{x}}\right)_{1...s}$. Finally, using $\hat{\boldsymbol{\alpha}}_{1...s}$, $\widehat{\boldsymbol{\lambda}}_{1...s}$, $(\widehat{\boldsymbol{\pi}}_{\boldsymbol{x}})_{1...s}$ and $\left(\widehat{\boldsymbol{\sigma}}^{\mathbf{2}}_{\boldsymbol{x}}\right)_{1...s}$ and (S53) we estimate $s$ corrected effects, $(\widehat{\boldsymbol{\alpha}}_{\boldsymbol{c}})_{1...s}$, to approximate the distribution of the corrected effect. The standard error of the corrected effect is estimated by taking the standard deviation of the $s$ corrected effects estimates, i.e. $\widehat{\mathrm{Var}\left(\widehat{\alpha}_c\right)} = \sum_{j=1}^{s}\left((\widehat{\boldsymbol{\alpha}}_{\boldsymbol{c}})_j - \widehat{\alpha}_c\right)^2/(s-1)$.

In addition, we can use the $s$ simulated IVW-based and corrected effects ($\hat{\boldsymbol{\alpha}}_{1...s}$ and $(\widehat{\boldsymbol{\alpha}_{\boldsymbol{c}}})_{1...s}$) to estimate the covariance between them. This covariance estimate can be used to test if there is a difference between IVW-based and corrected effects. Namely, the test statistic we use is

$$\frac{\widehat{\alpha} - \widehat{\alpha}_c}{\sqrt{\mathrm{Var}\left(\widehat{\alpha}\right) + \mathrm{Var}\left(\widehat{\alpha}_c\right) - 2\cdot\mathrm{Cov}\left(\widehat{\alpha}, \widehat{\alpha}_c\right)}} \sim \mathcal{N}(0,1) \tag{S59}$$

Since the number of simulations ($s$) that is used can influence the results - *if too small* - as well as the runtime - *if too large* - we decided to implement a procedure to automatically determine the best $s$ value. To do so, we start with $s = 1,000$ and create 10 random subsets to obtain 10 values for the variance of the corrected effect ($\boldsymbol{var}_{subsets}$) and 10 values for the covariance between the corrected and the IVW-based effect ($\boldsymbol{cov}_{subsets}$). We then estimate the coefficients of variation ($c_{var}$ and $c_{cov}$) for both parameters

$$c_{var} = \frac{sd(\boldsymbol{var}_{subsets})}{mean(\boldsymbol{var}_{subsets})} \tag{S60}$$

$$c_{cov} = \frac{sd(\boldsymbol{cov}_{subsets})}{mean(\boldsymbol{cov}_{subsets})} \tag{S61}$$

$$\tag{S62}$$

If one of the coefficient of variations is larger that 5%, or if the denominator of the test-statistic is undefined (S59), when $\mathrm{Var}\left(\widehat{\alpha}\right) + \mathrm{Var}\left(\widehat{\alpha}_c\right) - 2 \cdot \mathrm{Cov}\left(\widehat{\alpha}, \widehat{\alpha}_c\right) < 0$), we increase the value of $s$ to $s + 1,000$, and calculate again the coefficients of variation using 10 subsets. This is repeated until all criteria (coefficients of variation smaller than 5% and denominator defined) are fulfilled.

We validated this approach by comparing the standard error obtained as described above to the one that can be derived from the 100 simulated datasets (standard settings). For each threshold and each overlap, we calculated the standard error for the corrected effect of each dataset using our approach, and compared the mean of these standard errors to the standard deviation of the corrected effects estimates across the 100 datasets (Figure S19) and observed a very good agreement.

#### D. Simplification under the null

In Burgess *et al.* [4], a formula for the expected bias under the null is proposed. In order to compare our results with the ones from this paper, we had to make a few additional simplifications. In their simulation design, all genetic variants were considered to be causal ($\pi_x = 1$) and were used (no threshold used to select IVs).

When setting $\alpha = 0$ and $\pi_x = 1$ in (S51) we obtain

$$\begin{aligned}
\mathrm{E}\left[\hat{\alpha}_{IVW}\right] &= \lambda' \cdot \frac{\left(2 \cdot a + b \cdot (1 + n_A \cdot \sigma_X^2)\right)}{d(n_A, T, \pi_x, \sigma_x^2)} \\
&= \lambda' \cdot \frac{\left(2 \cdot a + b \cdot (1 + n_A \cdot \sigma_x^2)\right)}{2 \cdot a \cdot (\sigma_x^2 + \frac{1}{n_A}) + b \cdot (n_A \cdot \sigma_x^2 + 2 \cdot \sigma_x^2 + \frac{1}{nA})} \\
&= \lambda' \cdot \frac{\left(2 \cdot a + b \cdot (1 + n_A \cdot \sigma_x^2)\right)}{d(n_A, T, \pi_x, \sigma_x^2)} \\
&= \lambda' \cdot \frac{\left(2 \cdot a + b \cdot (1 + n_A \cdot \sigma_x^2)\right)}{(\sigma_x^2 + \frac{1}{n_A}) \cdot (2 \cdot a + b \cdot (1 + n_A \cdot \sigma_x^2)} \\
&= \frac{\lambda'}{\sigma_x^2 + \frac{1}{n_A}}
\end{aligned} \tag{S63}$$

We can then use (S33) to replace $\lambda'$

$$\mathrm{E}\left[\hat{\alpha}_{IVW}\right] = \rho \cdot \frac{n_{A \cap B}}{n_A \cdot n_B} \cdot \frac{1}{\sigma_x^2 + \frac{1}{n_A}} \tag{S64}$$

The simulation design proposed in their paper (Burgess *et al.* [4], Section 3 - "SIMULATION STUDY—CONTINUOUS OUTCOME") relies on non-standardised genotypic and phenotypic effects. In order to be compared, the expectation of the causal effect estimate we derived needs to be on the same scale and it can be done using the phenotypic variances of $X$ and $Y$ (respectively $\mathrm{Var}(x)$ and $\mathrm{Var}(y)$)

$$\mathrm{E}\left[\hat{\alpha}_{IVW}\right] = \frac{\sqrt{\mathrm{Var}(x)}}{\sqrt{\mathrm{Var}(y)}} \cdot \rho \cdot \frac{n_{A \cap B}}{n_A \cdot n_B} \cdot \frac{1}{\sigma_x^2 + \frac{1}{n_A}} \tag{S65}$$

The expectation of the causal effect estimate proposed by Burgess *et al.* [4] is as follows:

$$\mathrm{E}\left[\hat{\alpha}_{IVW}\right]_B = \text{OLS estimate} \cdot \frac{\%\text{ of overlap}}{100} \cdot \frac{1}{F} \tag{S66}$$

with:

$$\text{OLS estimate} = \frac{\mathrm{Cov}(x, y)}{\mathrm{Var}(x)}$$

$$= \frac{\sqrt{\mathrm{Var}\,(x)}}{\sqrt{\mathrm{Var}\,(y)}} \cdot \rho \text{ since } \rho \text{ is the correlation between } X \text{ and } Y \tag{S67}$$

$$\frac{\%\text{ of overlap}}{100} = \frac{n_{A\cap B}}{n_B} \tag{S68}$$

$$F = \frac{n_A - m - 1}{m} \cdot \frac{R^2}{1 - R^2} \tag{S69}$$

$$(R^2 \text{ being the coefficient of determination})$$

leading to:

$$\mathrm{E}\left[\hat{\alpha}_{IVW}\right]_B = \frac{\sqrt{\mathrm{Var}\,(x)}}{\sqrt{\mathrm{Var}\,(y)}} \cdot \rho \cdot \frac{n_{A\cap B}}{n_B} \cdot \frac{1}{\dfrac{n_A - m - 1}{m} \cdot \dfrac{R^2}{1 - R^2}} \tag{S70}$$

By defining $\sigma_x^2 = \frac{R^2}{m}$, it can be shown that when $n_A >> m$ and $R^2$ is small

$$\frac{1}{n_A} \cdot \frac{1}{\sigma_x^2 + \dfrac{1}{n_A}} \approx \frac{1}{\dfrac{n_A - m - 1}{m} \cdot \dfrac{R^2}{1 - R^2}} \tag{S71}$$

In this case, the expectation of the causal effect estimate we derived (S65) is equivalent to the one proposed by Burgess *et al.* [4] (S70).

We compared causal effect estimate expectations for all the scenarios without a causal effect they presented (Burgess *et al.* [4], Table 3). To do so, we calculated the theoretical values for $\mathrm{Var}\,(x)$, $\mathrm{Var}\,(y)$, OLS estimate and $R^2$ based on the parameters reported in the table and plugged them into (S65) and (S70). The causal effect estimate expectations can be compared to the IVW-based effects they reported. The expectation of the causal effect estimate we derived give results that are very close to the reported effect for all scenarios while the one proposed by Burgess *et al.* [4] suffer from a slight overestimation (Supplementary Table S32). It can be due to the fact that we used the theoretical values of $R^2$ which are lower than the observed ones because of overfitting.

### E. Additional simulation designs

#### a. Using case-control data for the exposure

We used the liability threshold model [5] to simulate binary (case-control) data for the exposure. We first simulated the exposure data on the liability scale ($X^*$) before converting it to the observed scale ($X$).

Causal SNPs for the exposure were randomly drawn from the set of 1,150,000 genetic variants, based on the polygenicity of the exposure ($\pi_x$) and their effects on $X^*$ were simulated using the liability-scale heritability of the exposure ($h_x^2$) as follows

$$\boldsymbol{\gamma_x} \sim \begin{cases} 0 & \text{with probability } 1-\pi_x\text{, for non-causal variants} \\ \mathcal{N}\Big(0, \dfrac{h_x^2}{M \cdot \pi_x}\Big) & \text{with probability } \pi_x\text{, for causal variants} \end{cases} \tag{S72}$$

For simplicity (and without loss of generality), we assumed that there were no direct genetic effects on the outcome. Then, phenotypic data for $X^*$, $X$ and $Y$ were simulated for all individuals included in the exposure or in any of the outcome samples, taking into account the effect of the confounder $U$ on $X^*$ and $Y$ (respectively $\kappa_x$ and $\kappa_y$) and the causal effect of $X$ on $Y$ ($\alpha$), using the following design

$$\boldsymbol{u} \sim \mathcal{N}(0,1) \tag{S73}$$

$$\boldsymbol{x^*} = G \cdot \boldsymbol{\gamma_x} + \boldsymbol{u} \cdot \kappa_x + \epsilon_x \tag{S74}$$

$$\text{with } \epsilon_x \sim \mathcal{N}(0, 1-(h_x^2+\kappa_x^2))$$

$$\boldsymbol{x} \sim \begin{cases} 0 & \text{if } \boldsymbol{x^*} \leq t \\ 1 & \text{if } \boldsymbol{x^*} > t \end{cases} \tag{S75}$$

$$t \text{ was defined using the prevalence, } p,\ t = -\Phi^{-1}(p)$$

$$\boldsymbol{x} \text{ was then scaled to have zero mean and variance of 1}$$

$$\boldsymbol{y} = \alpha \cdot \boldsymbol{x} + \boldsymbol{u} \cdot \kappa_y + \epsilon_y \tag{S76}$$

$$\text{with } \epsilon_y \sim \mathcal{N}(0, 1-(\alpha^2+\kappa_y^2 + 2 \cdot \alpha \cdot \kappa_y \cdot \text{Cov}\,(\boldsymbol{x},\boldsymbol{u})))$$

$$\text{Cov}\,(\boldsymbol{x},\boldsymbol{u}) \text{ was estimated directly from the data}$$

This simulation design also ensures that both $X$ and $Y$ have a zero mean and a variance of 1. Phenotypic data obtained using this design could be further used as described in Section 2.2 of the main text.

#### b. Simulations in presence of uncorrelated pleiotropy

To simulate uncorrelated pleiotropy, we assumed that a large proportion (60%) of the SNPs that were causally affecting the exposure were also causal SNPs for the outcome.

Causal SNPs for the exposure were randomly drawn from the set of 1,150,000 genetic variants, based on the polygenicity of $X$ ($\pi_x$) and their effects were simulated using the heritability of $X$

($h_x^2$) as follows

$$\boldsymbol{\gamma_x} \sim \begin{cases} 0 & \text{with probability } 1-\pi_x \text{, for non-causal variants} \\ \mathcal{N}\Big(0, \dfrac{h_x^2}{M \cdot \pi_x}\Big) & \text{with probability } \pi_x \text{, for causal variants} \end{cases} \tag{S77}$$

Similarly, causal SNPs for the outcome were semi-randomly drawn from our set of 1,500,000 SNPs (depending on the polygenicity of $Y$, $\pi_y$ - we forced 60% of the causal SNPs for the exposure to also be causal SNPs for the outcome) and their effect were simulated (using the heritability of $Y$, $h_y^2$) as follows:

$$\boldsymbol{\gamma_y} \sim \begin{cases} 0 & \text{with probability } 1-\pi_y \text{, for non-causal variants} \\ \mathcal{N}\Big(0, \dfrac{h_y^2}{M \cdot \pi_y}\Big) & \text{with probability } \pi_y \text{, for causal variants} \end{cases} \tag{S78}$$

Then, phenotypic data for $X$ and $Y$ were simulated for all individuals included in the exposure or in any of the outcome samples, taking into account the effect of the confounder $U$ on $X$ and $Y$ (respectively $\kappa_x$ and $\kappa_y$) and the causal effect of $X$ on $Y$ ($\alpha$), using the following design

$$\boldsymbol{u} \sim \mathcal{N}(0,1) \tag{S79}$$

$$\boldsymbol{x} = G \cdot \boldsymbol{\gamma_x} + \boldsymbol{u} \cdot \kappa_x + \epsilon_x \tag{S80}$$

$$\text{with } \epsilon_x \sim \mathcal{N}(0, 1-(h_x^2+\kappa_x^2))$$

$$\boldsymbol{y} = G \cdot \boldsymbol{\gamma_y} + \alpha \cdot \boldsymbol{x} + \boldsymbol{u} \cdot \kappa_y + \epsilon_y \tag{S81}$$

$$\text{with } \epsilon_y \sim \mathcal{N}(0, 1-(h_y^2+\alpha^2+\kappa_y^2+2\cdot\alpha\cdot\kappa_y\cdot\kappa_x))$$

This simulation design also ensures that both $X$ and $Y$ have a zero mean and a variance of 1. Phenotypic data obtained using this design could be further used as described in Section 2.2 of the main text.

#### c. Simulations in presence of correlated pleiotropy

To simulate data in presence of correlated pleiotropy, we added to the model a genetic confounder ($U_g$) acting both on $X$ and $Y$, with respective effects $q_x$ and $q_y$.
Causal SNPs for the exposure were randomly drawn from the set of 1,150,000 genetic variants, based on the polygenicity of the exposure ($\pi_x$) and their effects on $X^*$ were simulated using the liability-scale heritability of the exposure ($h_x^2$) as follows

$$\boldsymbol{\gamma_x} \sim \begin{cases} 0 & \text{with probability } 1-\pi_x \text{, for non-causal variants} \\ \mathcal{N}\Big(0, \dfrac{h_x^2}{M \cdot \pi_x}\Big) & \text{with probability } \pi_x \text{, for causal variants} \end{cases} \tag{S82}$$

Similarly, causal SNPs for the genetic confounder were randomly drown from our set of 1,500,000 SNPs (depending on the polygenicity of $U_g$, $\pi_u$ - causal SNPs for the genetic confounder randomly overlap with causal SNPs for the exposure) and their effect were simulated (using the heritability of $U_g$, $h_u^2$) as follows

$$\boldsymbol{\gamma_u} \sim \begin{cases} 0 & \text{with probability } 1-\pi_u \text{, for non-causal variants} \\ \mathcal{N}\Big(0, \dfrac{h_u^2}{M \cdot \pi_u}\Big) & \text{with probability } \pi_u \text{, for causal variants} \end{cases} \tag{S83}$$

For simplicity, we assumed that there were no direct genetic effects on the outcome. Then, phenotypic data for $X$ and $Y$ were simulated for all individuals included in the exposure or in any of the outcome samples, taking into account the effect of the environmental confounder $U$ on $X$ and $Y$ (respectively $\kappa_x$ and $\kappa_y$), the effect of the genetic confounder $U_g$ on $X$ and $Y$ (respectively $q_x$ and $q_y$) and the causal effect of $X$ on $Y$ ($\alpha$), using the following design:

$$\boldsymbol{u} \sim \mathcal{N}(0,1) \tag{S84}$$

$$\boldsymbol{u_g} = G \cdot \boldsymbol{\gamma_u} + \epsilon_u \tag{S85}$$

$$\text{with } \epsilon_u \sim \mathcal{N}(0, 1-h_u^2)$$

$$\boldsymbol{x} = G \cdot \boldsymbol{\gamma_x} + \boldsymbol{u} \cdot \kappa_x + \boldsymbol{u_g} \cdot q_x + \epsilon_x \tag{S86}$$

$$\text{with } \epsilon_x \sim \mathcal{N}(0, 1-(h_x^2+\kappa_x^2+q_x^2))$$

$$\boldsymbol{y} = \alpha \cdot \boldsymbol{x} + \boldsymbol{u} \cdot \kappa_y + \boldsymbol{u_g} \cdot q_y + \epsilon_y \tag{S87}$$

$$\text{with } \epsilon_y \sim \mathcal{N}(0, 1-(\alpha^2+\kappa_y^2+2\cdot\alpha\cdot\kappa_y\cdot\kappa_x+q_y^2+2\cdot\alpha\cdot q_y\cdot q_x))$$

This simulation design also ensures that both $X$ and $Y$ have a zero mean and a variance of 1. Phenotypic data obtained using this design could be further used as described in Section 2.2 of the main text.

### Supplementary Figures


S1 Effect of the different parameters when overlap = 0% . . . . . . . . . . . 24
S2 Effect of the different parameters when overlap = 20% . . . . . . . . . . 25
S3 Effect of the different parameters when overlap = 80% . . . . . . . . . . 26
S4 Effect of the different parameters when overlap = 100% . . . . . . . . . 27
S5 Simulation results for a scenario with a weaker environmental confounder 29
S6 Simulation results for a scenario with a stronger environmental confounder . . . . . . . . . . . . . . . . . . . . . . . . . . . . . . . . . . . . 31
S7 Simulation results in the absence of a causal effect . . . . . . . . . . . . 32
S8 False positive rate in the absence of a causal effect . . . . . . . . . . . . 33
S9 Simulation results for a case-control design . . . . . . . . . . . . . . . . 34
S10 Simulation results for realistic settings . . . . . . . . . . . . . . . . . . . 36
S11 Simulation results in presence of uncorrelated pleiotropy . . . . . . . . . 38
S12 Simulation results in presence of moderate correlated pleiotropy . . . . 40
S13 Simulation results in presence of strong correlated pleiotropy . . . . . . 41
S14 Comparison of different MR approaches (less stringent selection threshold for MR-RAPS and dIVW) . . . . . . . . . . . . . . . . . . . . . . . . 42
S15 Comparison of different MR approaches (no selection for MR-RAPS and dIVW) . . . . . . . . . . . . . . . . . . . . . . . . . . . . . . . . . . 43
S16 Effect of BMI on smoking . . . . . . . . . . . . . . . . . . . . . . . . . . . 45
S17 Ratio of the expectation (simulated data) . . . . . . . . . . . . . . . . . . 46
S18 Ratio of the expectation (BMI-BMI data) . . . . . . . . . . . . . . . . . . 47
S19 Standard error estimation for the corrected effects . . . . . . . . . . . . . 48


### Supplementary Tables


S1 Description of the pairs of traits analysed . . . . . . . . . . . . . . . . . . 28
S3 Analysis of variance for standard settings . . . . . . . . . . . . . . . . . . 28
S5 Analysis of variance for a scenario with a weaker environmental confounder . . . . . . . . . . . . . . . . . . . . . . . . . . . . . . . . . . . . 30
S7 Analysis of variance for a scenario with a stronger environmental confounder . . . . . . . . . . . . . . . . . . . . . . . . . . . . . . . . . . . . 30
S9 Analysis of variance for a scenario with a negative confounder . . . . . 30
S11 Analysis of variance in the absence of a causal effect . . . . . . . . . . . 33
S13 Analysis of variance for a case-control design . . . . . . . . . . . . . . . 35
S15 Analysis of variance for realistic settings . . . . . . . . . . . . . . . . . . 35
S16 Increasing sample size and power by allowing and correcting for sample overlap . . . . . . . . . . . . . . . . . . . . . . . . . . . . . . . . . . . . 37
S18 Analysis of variance in presence of uncorrelated pleiotropy . . . . . . . . 39
S20 Analysis of variance in presence of moderate correlated pleiotropy . . . 39
S22 Analysis of variance in presence of strong correlated pleiotropy . . . . . 39

S26 **Analysis of variance for the effect of BMI on SBP** . . . . . . . . . . . . . 44
S28 **Analysis of variance for the effect of BMI on smoking** . . . . . . . . . . . 44
S30 **Analysis of variance for the effect of BMI on alcohol** . . . . . . . . . . . . 45

Supplementary tables S2, S4, S6, S8, S10, S12, S14, S17, S19, S21, S23, S24, S25, S27, S29 are too large to be included in this document and are available in an `xlsx` file.

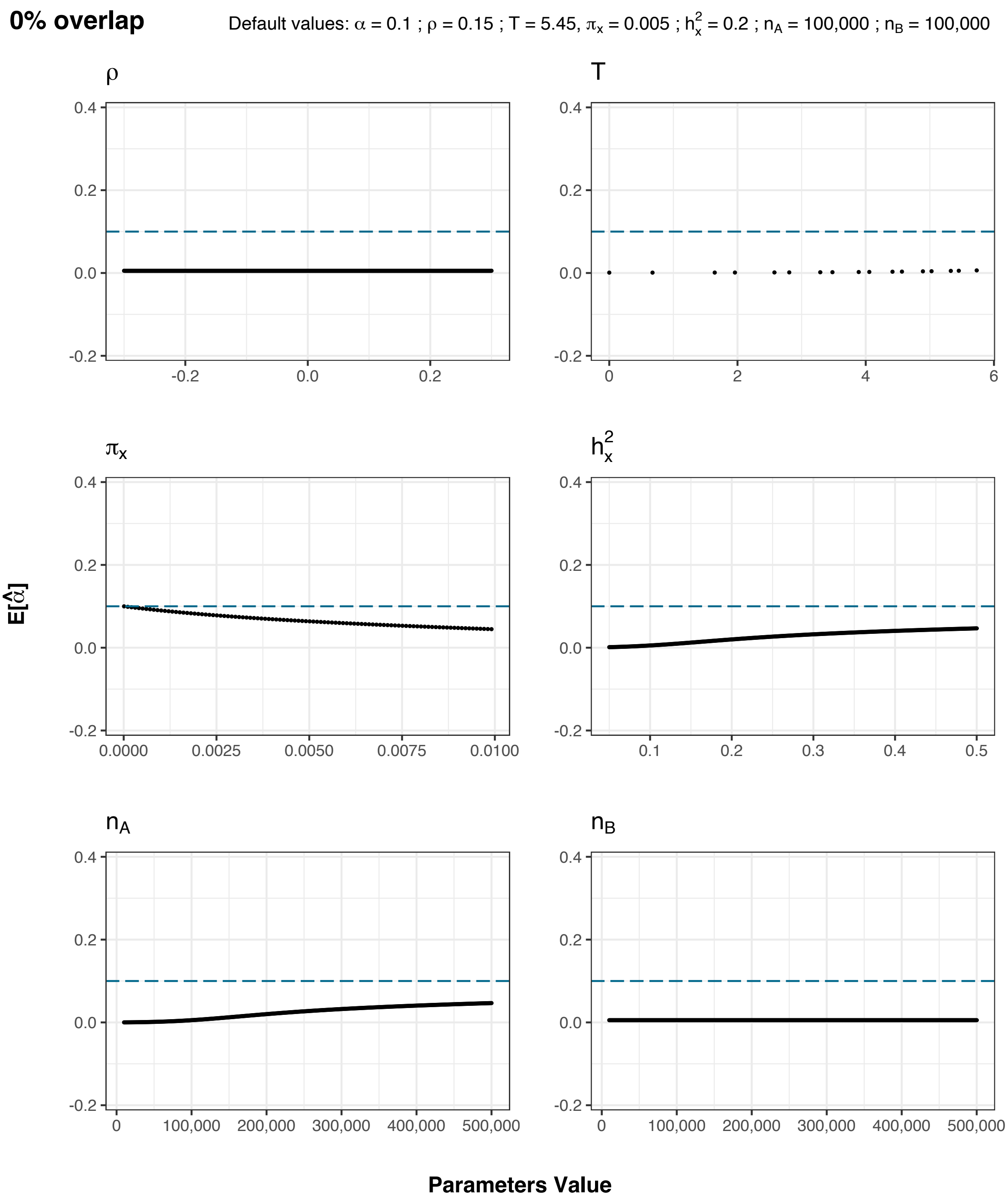


**Figure S1: Effect of the different parameters when overlap = 0%**
For each parameter affecting $E[\hat{\alpha}]$, we tested a wide range of values, while keeping the other parameters constant. The true causal effect value is indicated by the blue dotted line.
Defaults parameters are $n_A = 100,000, n_B = 100,000, \pi_x = 0.005, h^2_x = 0.2, \rho = 0.15, \alpha = 0.1$

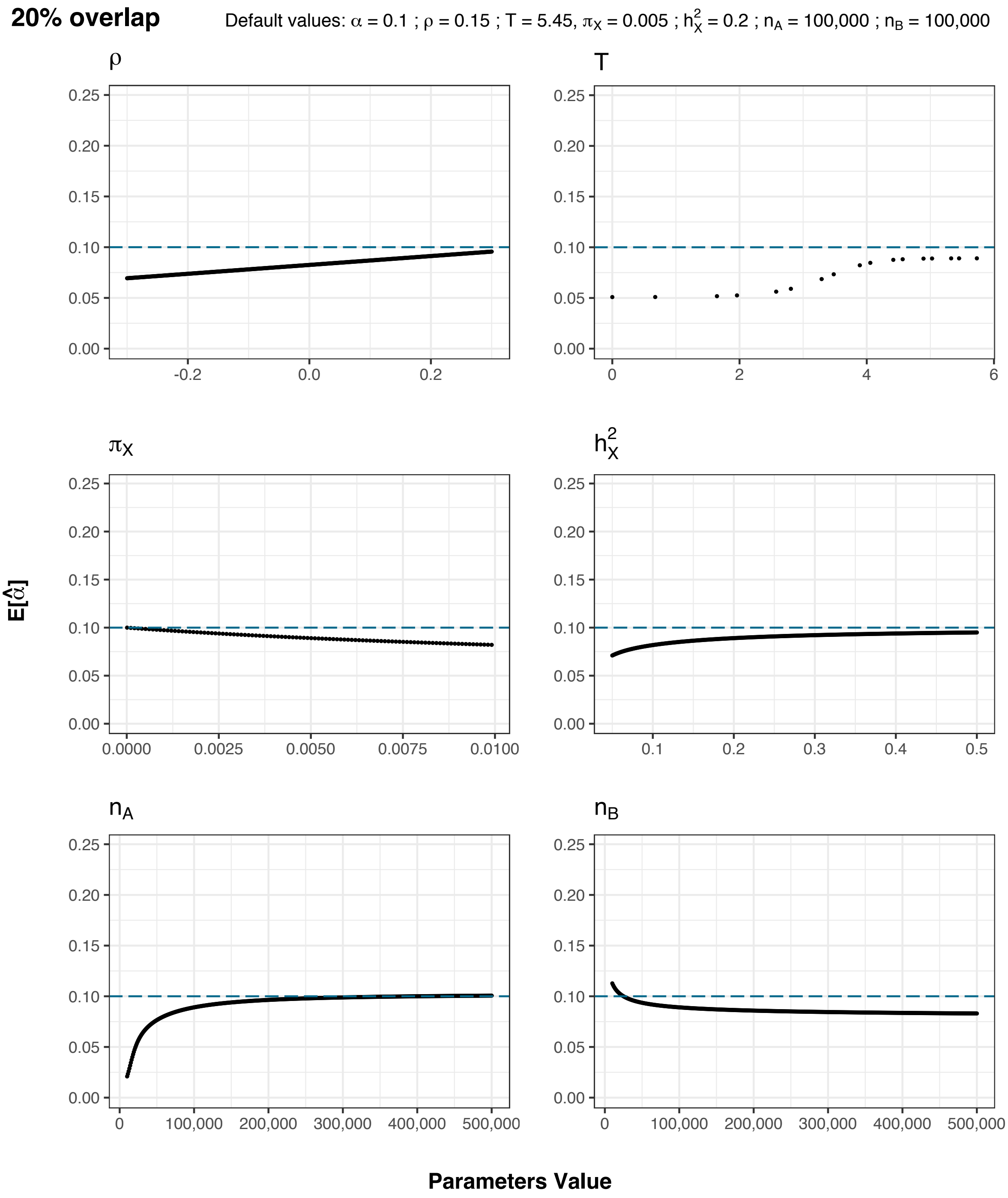


**Figure S2: Effect of the different parameters when overlap = 20%**
For each parameter affecting $E[\hat{\alpha}]$, we tested a wide range of values, while keeping the other parameters constant. The true causal effect value is indicated by the blue dotted line.
Defaults parameters are $n_A = 100,000, n_B = 100,000, \pi_x = 0.005, h_x^2 = 0.2, \rho = 0.15, \alpha = 0.1$

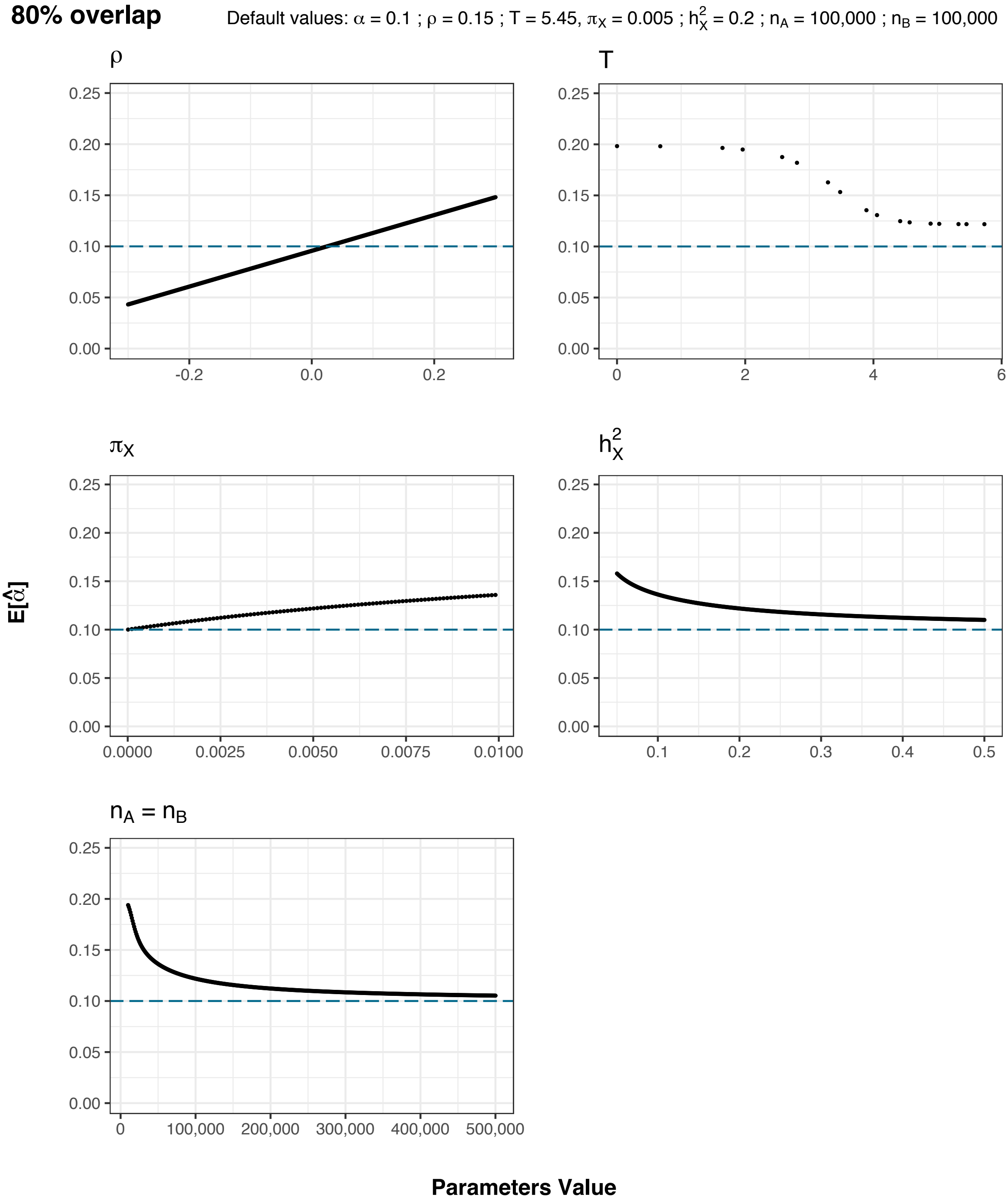


**Figure S3: Effect of the different parameters when overlap = 80%**
For each parameter affecting $E[\hat{\alpha}]$, we tested a wide range of values, while keeping the other parameters constant. The true causal effect value is indicated by the blue dotted line.
Defaults parameters are $n_A = 100,000, n_B = 100,000, \pi_x = 0.005, h_x^2 = 0.2, \rho = 0.15, \alpha = 0.1$
Note that for simplicity, $n_A$ and $n_B$ are assumed to be equal, to ensure that an overlap of 80% can be observed.

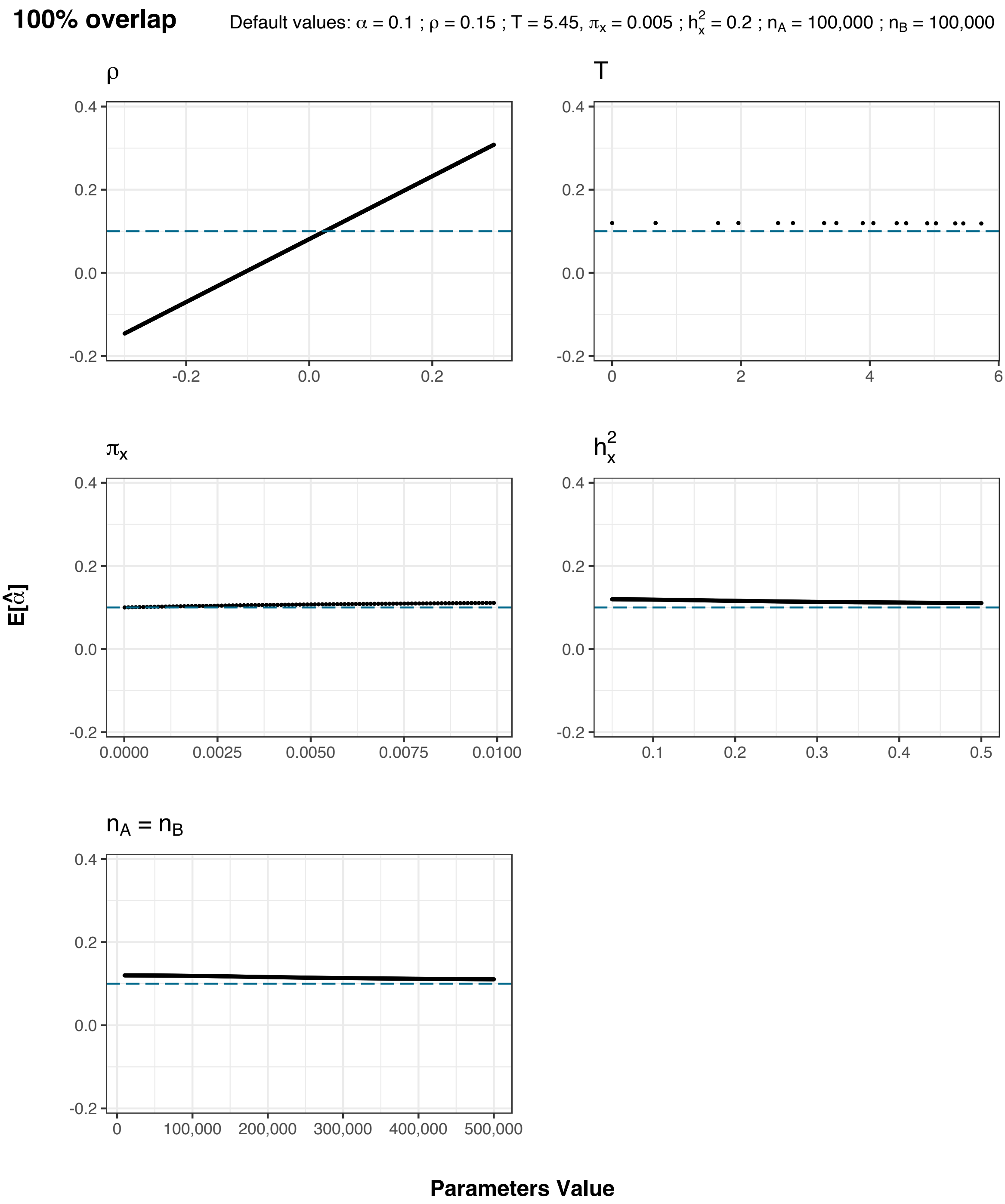


**Figure S4: Effect of the different parameters when overlap = 100%**
For each parameter affecting $E[\hat{\alpha}]$, we tested a wide range of values, while keeping the other parameters constant. The true causal effect value is indicated by the blue dotted line.
Defaults parameters are $n_A = 100,000, n_B = 100,000, \pi_x = 0.005, h_x^2 = 0.2, \rho = 0.15, \alpha = 0.1$
Note that for simplicity, $n_A$ and $n_B$ are assumed to be equal, to ensure that an overlap of 100% can be observed.

| Exposure | Outcome | Field ID Exposure | Field ID Outcome | N Exposure | N Outcome |
|---|---|---|---|---|---|
| BMI | BMI | 21001 | 21001 | 99,493 | 99,491 |
| SBP | SBP | 4080 | 4080 | 93,231 | 93,229 |
| BMI | SBP | 21001 | 4080 | 99,491 | 93,231 |
| BMI | smoking | 21001 | 2887 | 99,490 | 23,507 |
| BMI | alcohol | 21001 | 1558 | 99,493 | 99,808 |

**Table S1: Description of the pairs of traits analysed**
For each pair of trait analysed, we reported the field IDs corresponding to the exposure and the outcome, as well as the mean sample size (across the 100 repetitions, and the 5 overlaps for the outcome - note that for the BMI-BMI analysis, only non-overlapping samples were considered) of the phenotypic data used.

| | | **IVW-based effects** | | | **Corrected effects** | | |
|---|---|---|---|---|---|---|---|
| Threshold | IVs | Within groups | Between groups | Ratio | Within groups | Between groups | Ratio |
| 1e-08 | 51.6 | 0.000433 | 0.0375 | 86.6 | 0.000554 | 0.00178 | 3.22 |
| 5e-08 | 65.4 | 0.000339 | 0.0380 | 112.1 | 0.000434 | 0.00212 | 4.89 |
| 1e-07 | 72.4 | 0.000307 | 0.0391 | 127.3 | 0.000394 | 0.00249 | 6.34 |
| 5e-07 | 92.6 | 0.000259 | 0.0462 | 178.6 | 0.000329 | 0.00483 | 14.67 |
| 1e-06 | 103.0 | 0.000250 | 0.0494 | 197.6 | 0.000319 | 0.00563 | 17.67 |

**Table S3: Analysis of variance for standard settings**
$n_A = n_B = 20,000, \pi_x = 0.001, h_x^2 = 0.4, \kappa_x = 0.3, \kappa_y = 0.5, \alpha = 0.2$
For each threshold, the mean number of instruments used (IVs), the within groups and between group variances, their ratio (between/within) for both IVW-based and corrected effects are reported.

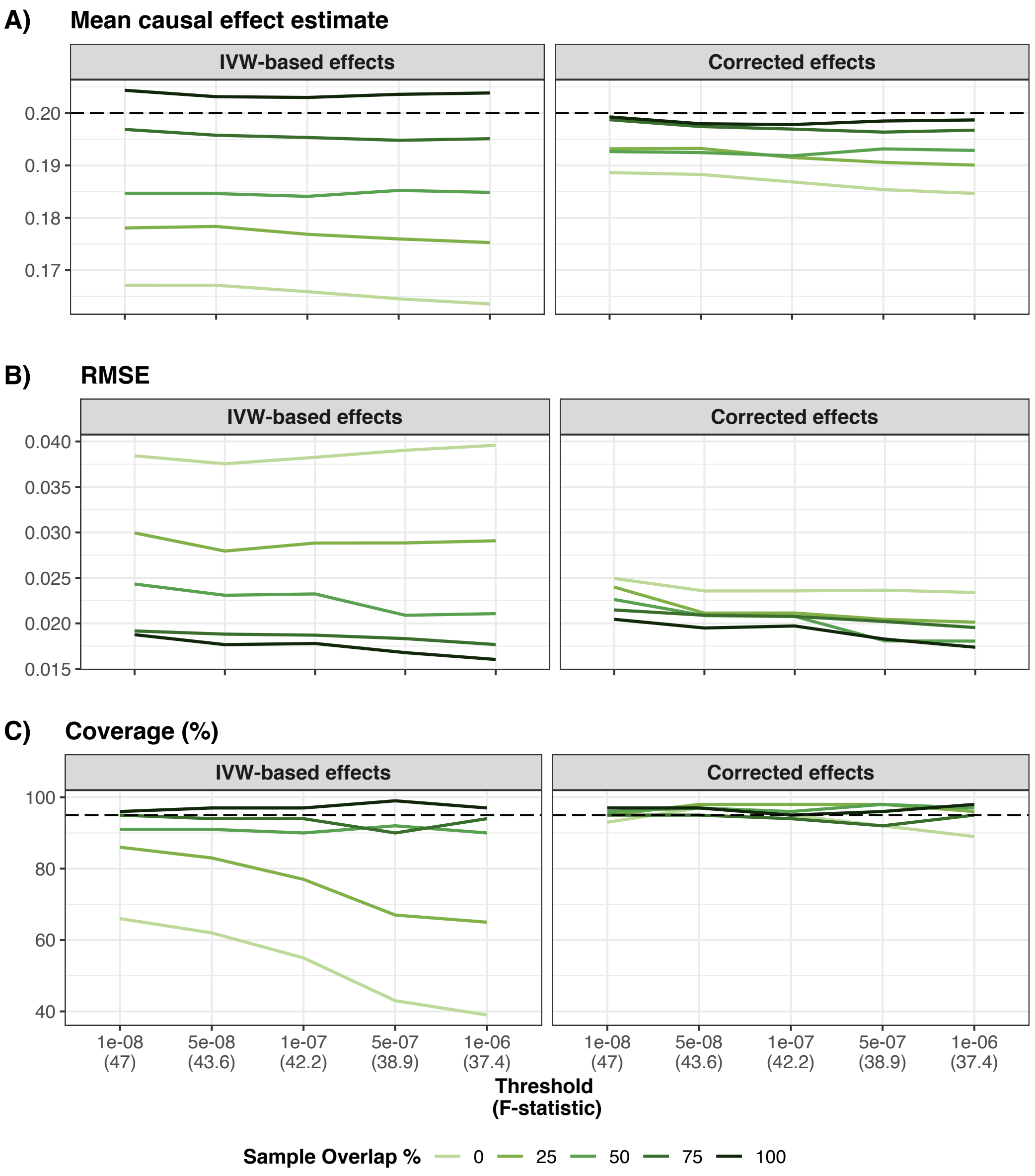


**Figure S5: Simulation results for a scenario with a weaker environmental confounder** $n_A = n_B = 20,000, \pi_x = 0.001, h_x^2 = 0.4, \kappa_x = 0.15, \kappa_y = 0.3, \alpha = 0.2$
Panel A) shows the mean IVW-based and corrected effect for each overlap and threshold obtained from 100 simulations (the dashed line represents the true causal effect). Panel B) shows the mean RMSE obtained for IVW-based and corrected effect for each overlap and threshold. Panel C) shows the coverage of the 95% confidence interval for IVW-based and corrected effect for each overlap and threshold.

| | | IVW-based effects | | | Corrected effects | | |
|---|---|---|---|---|---|---|---|
| Threshold | IVs | Within groups | Between groups | Ratio | Within groups | Between groups | Ratio |
| 1e-08 | 50.6 | 0.000376 | 0.0218 | 58.1 | 0.000476 | 0.00200 | 4.20 |
| 5e-08 | 64.5 | 0.000319 | 0.0201 | 63.0 | 0.000404 | 0.00157 | 3.89 |
| 1e-07 | 71.2 | 0.000307 | 0.0215 | 70.0 | 0.000389 | 0.00200 | 5.12 |
| 5e-07 | 90.8 | 0.000266 | 0.0235 | 88.2 | 0.000340 | 0.00262 | 7.70 |
| 1e-06 | 101.2 | 0.000247 | 0.0252 | 102.2 | 0.000316 | 0.00309 | 9.78 |

**Table S5: Analysis of variance for a scenario with a weaker environmental confounder**
$n_A = n_B = 20,000, \pi_x = 0.001, h_x^2 = 0.4, \kappa_x = 0.15, \kappa_y = 0.3, \alpha = 0.2$
For each threshold, the mean number of instruments used (IVs), the within groups and between group variances, their ratio (between/within) for both IVW-based and corrected effects are reported.

| | | IVW-based effects | | | Corrected effects | | |
|---|---|---|---|---|---|---|---|
| Threshold | IVs | Within groups | Between groups | Ratio | Within groups | Between groups | Ratio |
| 1e-08 | 50.2 | 0.000382 | 0.119 | 312 | 0.000506 | 0.00591 | 11.7 |
| 5e-08 | 64.2 | 0.000320 | 0.125 | 390 | 0.000426 | 0.00815 | 19.1 |
| 1e-07 | 70.8 | 0.000307 | 0.127 | 414 | 0.000412 | 0.00893 | 21.7 |
| 5e-07 | 90.7 | 0.000275 | 0.142 | 516 | 0.000369 | 0.01347 | 36.6 |
| 1e-06 | 100.9 | 0.000258 | 0.151 | 587 | 0.000349 | 0.01559 | 44.7 |

**Table S7: Analysis of variance for a scenario with a stronger environmental confounder**
$n_A = n_B = 20,000, \pi_x = 0.001, h_x^2 = 0.4, \kappa_x = 0.5, \kappa_y = 0.8, \alpha = 0.2$
For each threshold, the mean number of instruments used (IVs), the within groups and between group variances, their ratio (between/within) for both IVW-based and corrected effects are reported.

| | | IVW-based effects | | | Corrected effects | | |
|---|---|---|---|---|---|---|---|
| Threshold | IVs | Within groups | Between groups | Ratio | Within groups | Between groups | Ratio |
| 1e-08 | 51.2 | 0.000423 | 0.00112 | 2.66 | 0.000535 | 0.000191 | 0.356 |
| 5e-08 | 65.4 | 0.000357 | 0.00153 | 4.27 | 0.000449 | 0.000419 | 0.935 |
| 1e-07 | 72.8 | 0.000334 | 0.00151 | 4.52 | 0.000421 | 0.000416 | 0.989 |
| 5e-07 | 92.8 | 0.000290 | 0.00169 | 5.83 | 0.000365 | 0.000524 | 1.436 |
| 1e-06 | 102.8 | 0.000290 | 0.00175 | 6.05 | 0.000367 | 0.000539 | 1.471 |

**Table S9: Analysis of variance for a scenario with a negative confounder**
$n_A = n_B = 20,000, \pi_x = 0.001, h_x^2 = 0.4, \kappa_x = -0.3, \kappa_y = 0.5, \alpha = 0.2$ For each threshold, the mean number of instruments used (IVs), the within groups and between group variances, their ratio (between/within) for both IVW-based and corrected effects are reported.

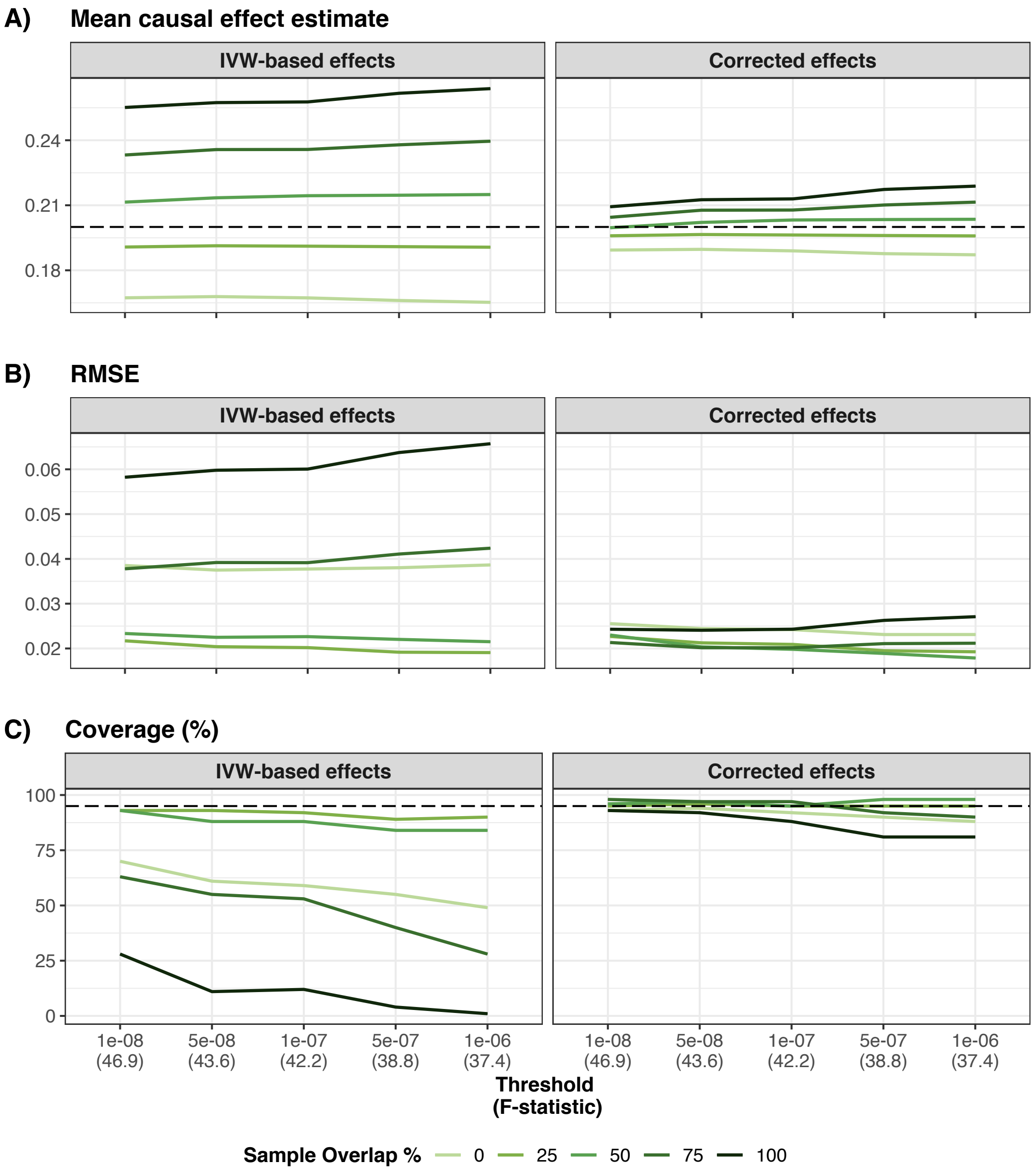


**Figure S6: Simulation results for a scenario with a stronger environmental confounder**

$n_A = n_B = 20,000, \pi_x = 0.001, h_x^2 = 0.4, \kappa_x = 0.5, \kappa_y = 0.8, \alpha = 0.2$

Panel A) shows the mean IVW-based and corrected effect for each overlap and threshold obtained from 100 simulations (the dashed line represents the true causal effect). Panel B) shows the mean RMSE obtained for IVW-based and corrected effect for each overlap and threshold. Panel C) shows the coverage of the 95% confidence interval for IVW-based and corrected effect for each overlap and threshold.

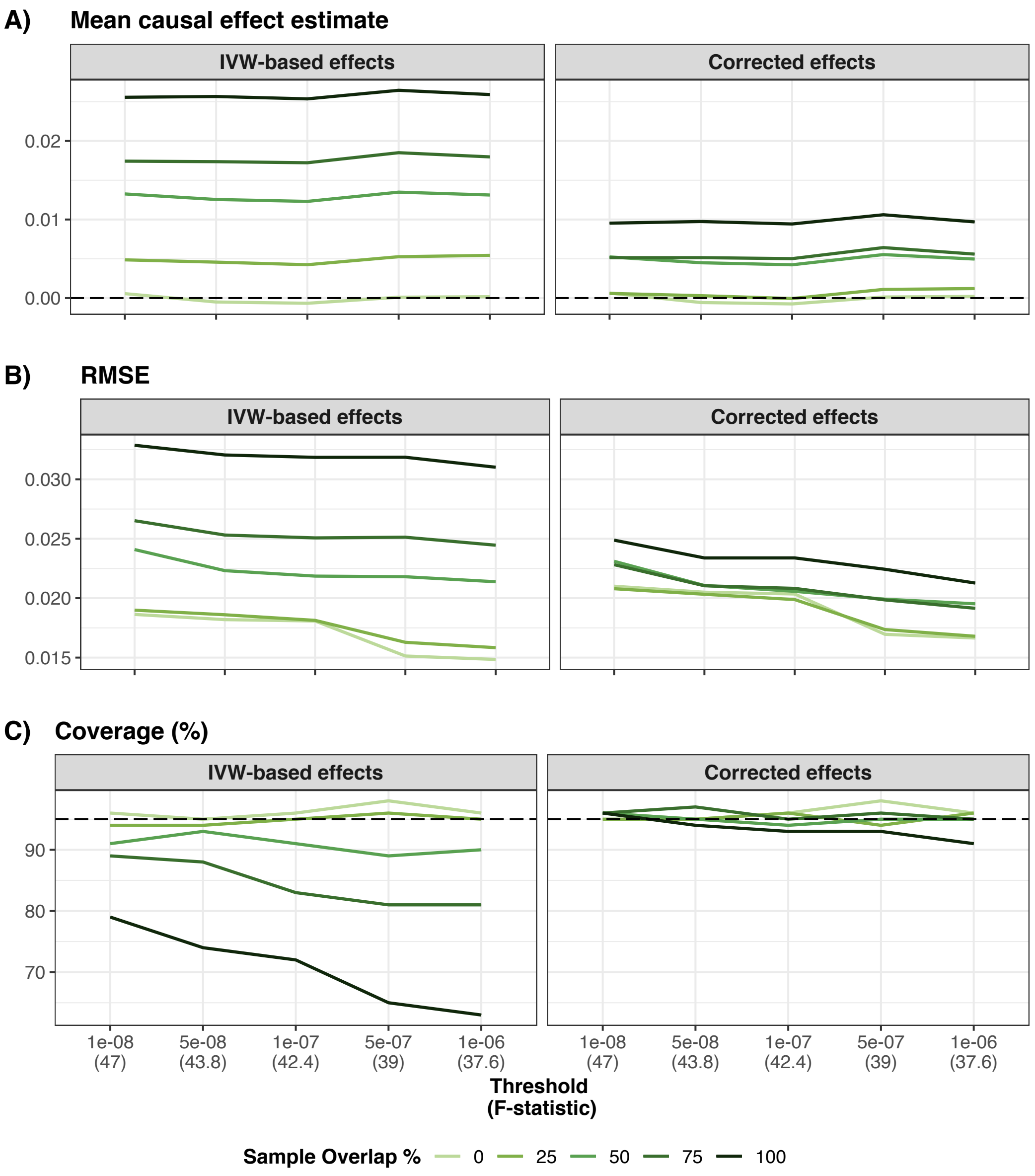


**Figure S7: Simulation results in the absence of a causal effect**
$n_A = n_B = 20,000, \pi_x = 0.001, h_x^2 = 0.4, \kappa_x = 0.3, \kappa_y = 0.5, \alpha = 0$
Panel A) shows the mean IVW-based and corrected effect for each overlap and threshold obtained from 100 simulations (the dashed line represents the true causal effect). Panel B) shows the mean RMSE obtained for IVW-based and corrected effect for each overlap and threshold. Panel C) shows the coverage of the 95% confidence interval for IVW-based and corrected effect for each overlap and threshold.

| | | IVW-based effects | | | Corrected effects | | |
|---|---|---|---|---|---|---|---|
| Threshold | IVs | Within groups | Between groups | Ratio | Within groups | Between groups | Ratio |
| 1e-08 | 52.3 | 0.000387 | 0.0099 | 25.6 | 0.000485 | 0.00140 | 2.89 |
| 5e-08 | 65.4 | 0.000344 | 0.0107 | 31.0 | 0.000429 | 0.00172 | 4.01 |
| 1e-07 | 72.2 | 0.000337 | 0.0106 | 31.6 | 0.000420 | 0.00172 | 4.09 |
| 5e-07 | 92.1 | 0.000276 | 0.0109 | 39.6 | 0.000343 | 0.00181 | 5.28 |
| 1e-06 | 102.4 | 0.000261 | 0.0103 | 39.4 | 0.000325 | 0.00144 | 4.43 |

**Table S11: Analysis of variance in the absence of a causal effect**
$n_A = n_B = 20,000, \pi_x = 0.001, h_x^2 = 0.4, \kappa_x = 0.3, \kappa_y = 0.5, \alpha = 0$
For each threshold, the mean number of instruments used (IVs), the within groups and between group variances, their ratio (between/within) for both IVW-based and corrected effects are reported.

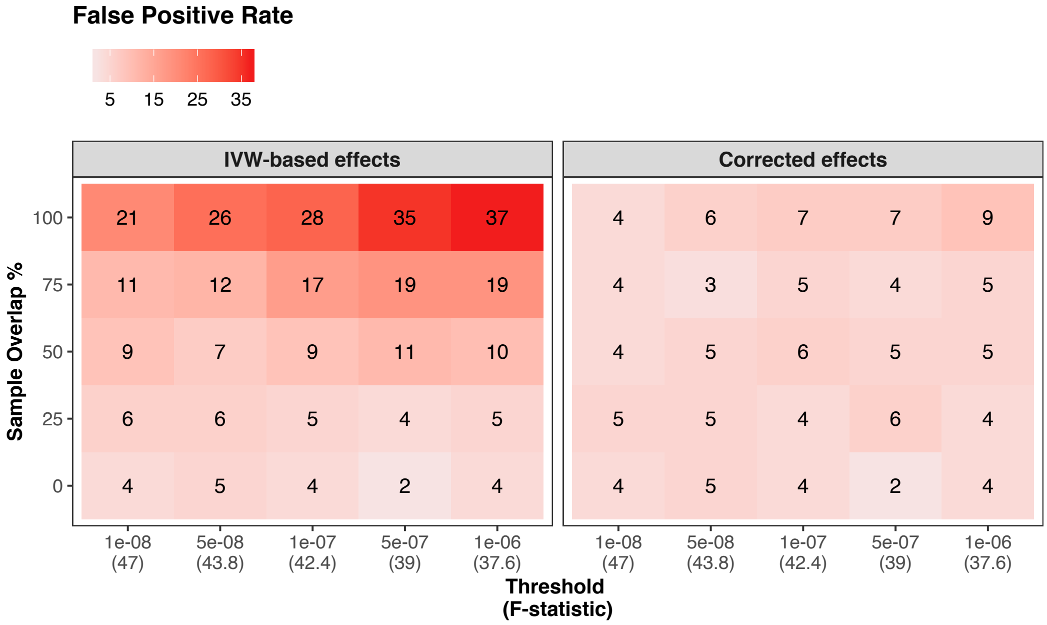


**Figure S8: False positive rate in the absence of a causal effect**
Panel A) shows the mean number of false positive signals for IVW-based effect estimates for each overlap and threshold obtained from 100 simulations. Panel B) shows the mean number of false positive signals for corrected effect estimates for each overlap and threshold obtained from 100 simulations

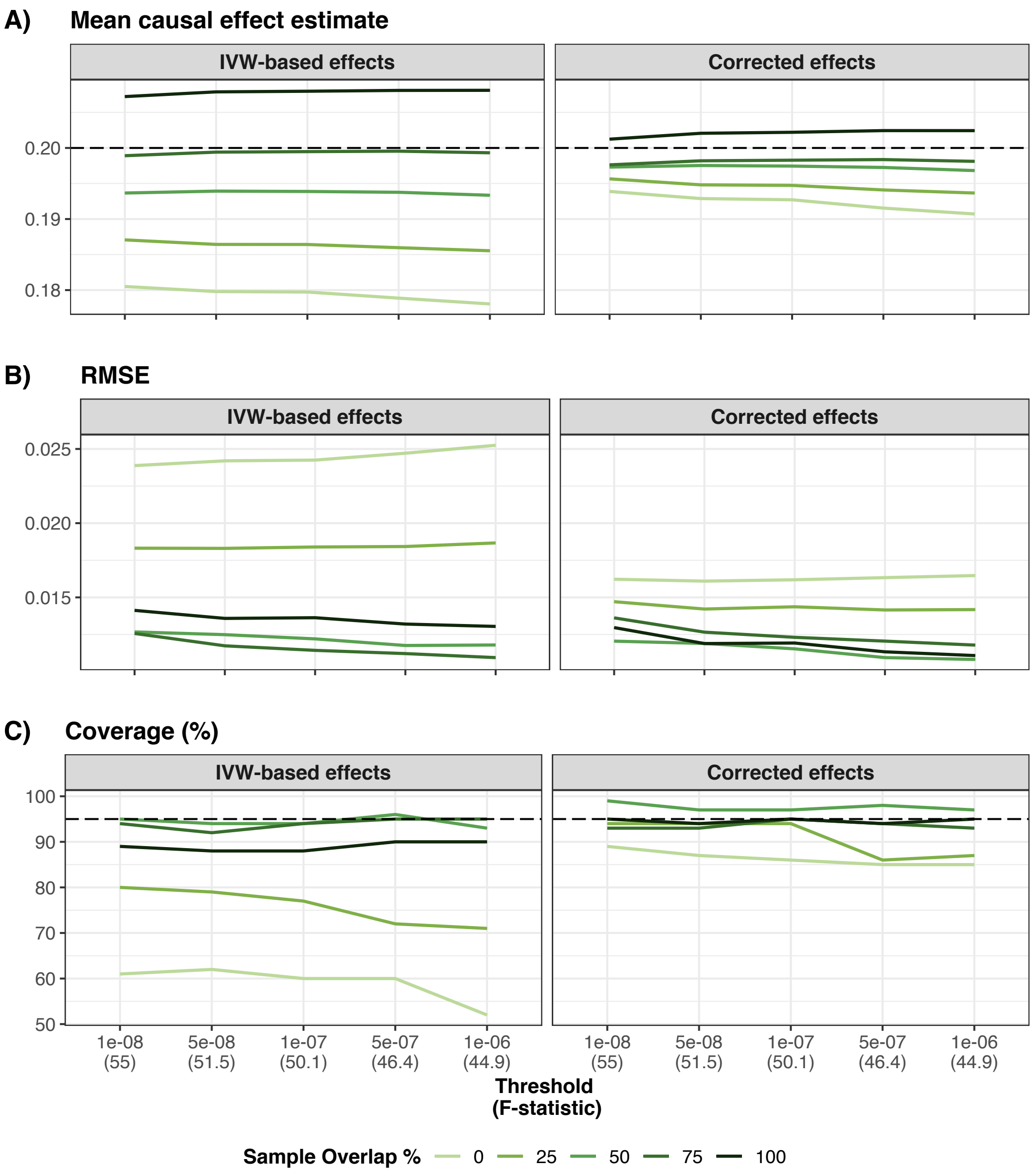


**Figure S9: Simulation results for a case-control design**
$n_A = n_B = 100,000, \pi_x = 0.001, h_x^2 = 0.4, \kappa_x = 0.3, \kappa_y = 0.5, \alpha = 0.2$, X is converted from the liability scale to the observed scale using a prevalence of 0.1
Panel A) shows the mean IVW-based and corrected effect for each overlap and threshold obtained from 100 simulations (the dashed line represents the true causal effect). Panel B) shows the mean RMSE obtained for IVW-based and corrected effect for each overlap and threshold. Panel C) shows the coverage of the 95% confidence interval for IVW-based and corrected effect for each overlap and threshold.

| | | IVW-based effects | | | Corrected effects | | |
|---|---|---|---|---|---|---|---|
| Threshold | IVs | Within groups | Between groups | Ratio | Within groups | Between groups | Ratio |
| 1e-08 | 126 | 0.000158 | 0.0107 | 67.6 | 0.000183 | 0.000746 | 4.07 |
| 5e-08 | 148 | 0.000143 | 0.0120 | 84.0 | 0.000164 | 0.001228 | 7.48 |
| 1e-07 | 158 | 0.000140 | 0.0121 | 86.4 | 0.000162 | 0.001312 | 8.12 |
| 5e-07 | 187 | 0.000129 | 0.0130 | 100.5 | 0.000149 | 0.001737 | 11.65 |
| 1e-06 | 202 | 0.000124 | 0.0137 | 110.4 | 0.000142 | 0.001986 | 13.95 |

**Table S13: Analysis of variance for a case-control design**
$n_A = n_B = 100,000, \pi_x = 0.001, h_x^2 = 0.4, \kappa_x = 0.3, \kappa_y = 0.5, \alpha = 0.2$, X is converted from the liability scale to the observed scale using a prevalence of 0.1
For each threshold, the mean number of instruments used (IVs), the within groups and between group variances, their ratio (between/within) for both IVW-based and corrected effects are reported.

| | | IVW-based effects | | | Corrected effects | | |
|---|---|---|---|---|---|---|---|
| Threshold | IVs | Within groups | Between groups | Ratio | Within groups | Between groups | Ratio |
| 1e-08 | 54.4 | 0.000457 | 0.0499 | 109 | 0.000695 | 0.00212 | 3.04 |
| 5e-08 | 76.6 | 0.000351 | 0.0479 | 137 | 0.000537 | 0.00168 | 3.14 |
| 1e-07 | 89.3 | 0.000310 | 0.0536 | 173 | 0.000476 | 0.00321 | 6.74 |
| 5e-07 | 129.6 | 0.000246 | 0.0565 | 229 | 0.000378 | 0.00424 | 11.23 |
| 1e-06 | 152.7 | 0.000210 | 0.0588 | 280 | 0.000325 | 0.00497 | 15.28 |

**Table S15: Analysis of variance for realistic settings**
$n_A = n_B = 100,000, \pi_x = 0.005, h_x^2 = 0.2, \kappa_x = 0.3, \kappa_y = 0.5, \alpha = 0.1$
For each threshold, the mean number of instruments used (IVs), the within groups and between group variances, their ratio (between/within) for both IVW-based and corrected effects are reported.

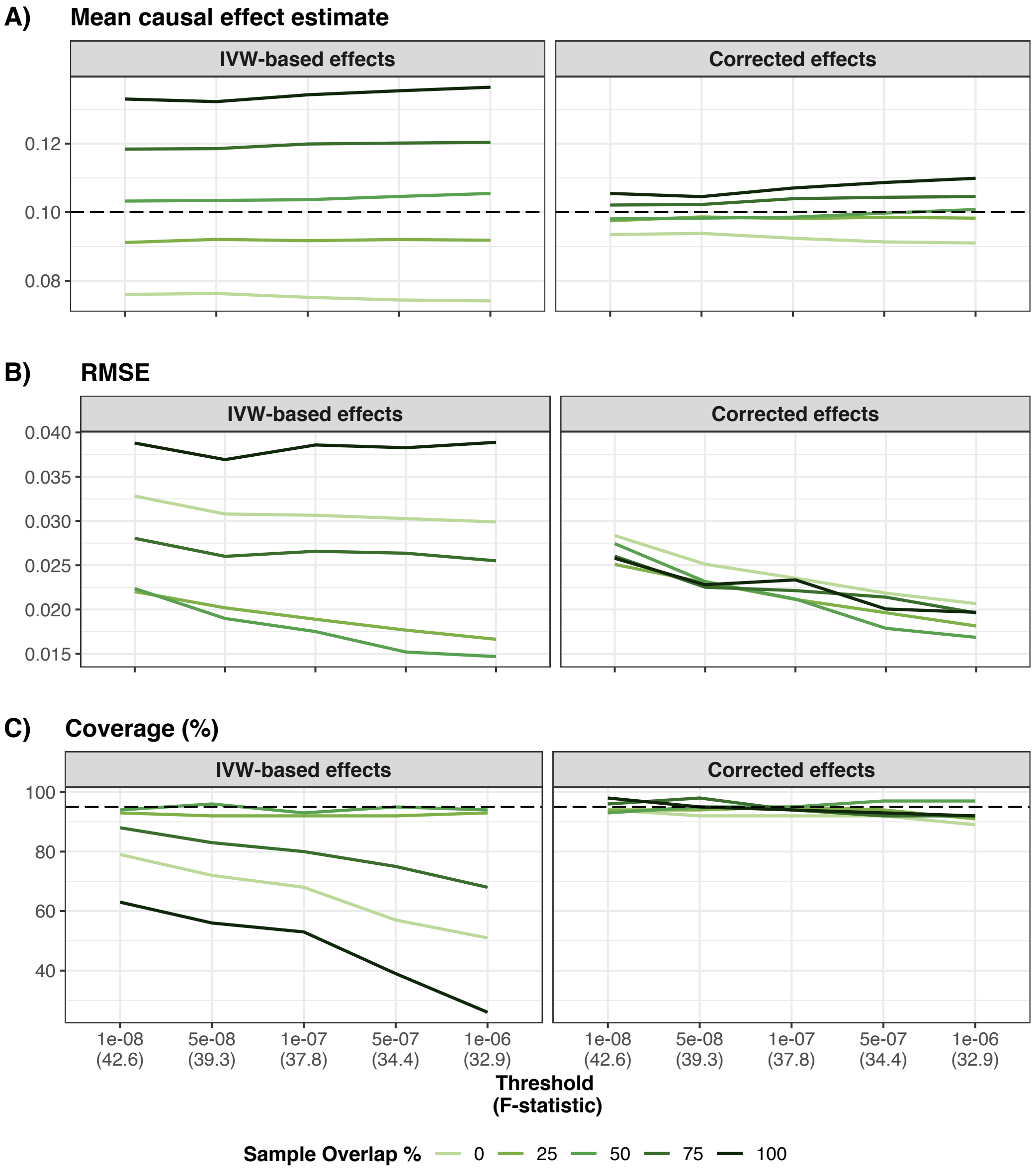


**Figure S10: Simulation results for realistic settings**
$n_A = n_B = 100,000, \pi_x = 0.005, h_x^2 = 0.2, \kappa_x = 0.3, \kappa_y = 0.5, \alpha = 0.1$
Panel A) shows the mean IVW-based and corrected effect for each overlap and threshold obtained from 100 simulations (the dashed line represents the true causal effect). Panel B) shows the mean RMSE obtained for IVW-based and corrected effect for each overlap and threshold. Panel C) shows the coverage of the 95% confidence interval for IVW-based and corrected effect for each overlap and threshold.

| Effect (sample) | IVW-based (non-overlapping samples) | Corrected (fully overlapping samples) |
|---|---|---|
| bias | -0.02937318 | 0.008976098 |
| variance | 0.00037905 | 0.000110344 |
| RMSE | 0.03518584 | 0.013777182 |
| coverage | 72 | 88 |
| power | 94 | 100 |

**Table S16: Increasing sample size and power by allowing and correcting for sample overlap**

fully overlapping samples: $n_A = n_B = 360,000, \pi_x = 0.01, h_x^2 = 0.15, \kappa_x = 0.3, \kappa_y = 0.5, \alpha = 0.1$
non-overlapping samples: $n_A = n_B = 180,000, \pi_x = 0.01, h_x^2 = 0.15, \kappa_x = 0.3, \kappa_y = 0.5, \alpha = 0.1$
For a threshold of 5e-8, we estimated the IVW-based effect obtained when splitting the sample into two non-overlapping halves and the corrected effect obtained using fully overlapping samples. We calculated the bias, the variance, the RMSE, the coverage and the power for both estimates.

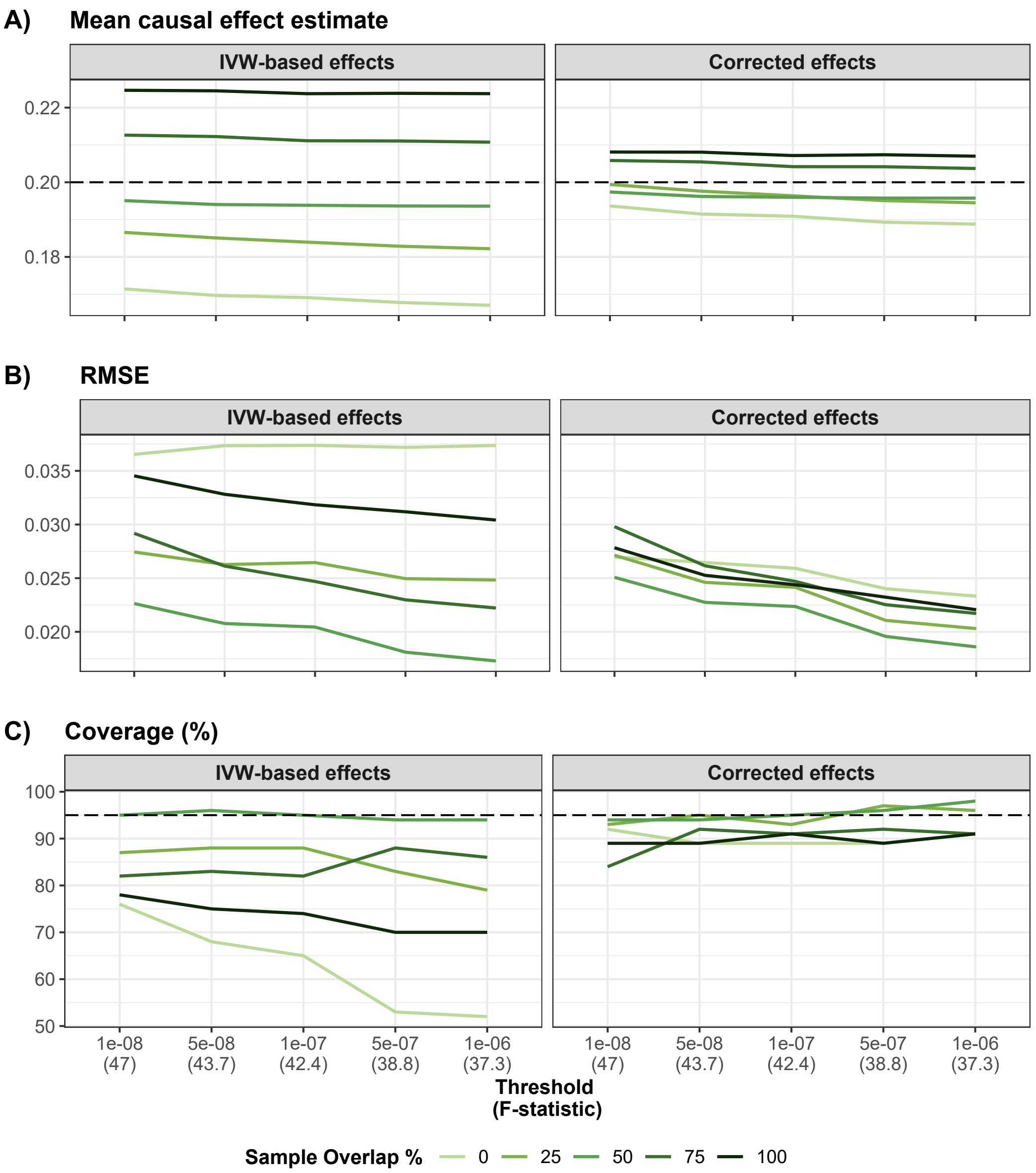


**Figure S11: Simulation results in presence of uncorrelated pleiotropy**
$n_A = n_B = 20,000, \pi_x = 0.001, h_x^2 = 0.4, \kappa_x = 0.3, \kappa_y = 0.5, \alpha = 0.2, \pi_y = 0.002, h_y^2 = 0.3,$ 60% of the causal SNPs for X were also causal for Y
Panel A) shows the mean IVW-based and corrected effect for each overlap and threshold obtained from 100 simulations (the dashed line represents the true causal effect). Panel B) shows the mean RMSE obtained for IVW-based and corrected effect for each overlap and threshold. Panel C) shows the coverage of the 95% confidence interval for IVW-based and corrected effect for each overlap and threshold.

| | | IVW-based effects | | | Corrected effects | | |
|---|---|---|---|---|---|---|---|
| Threshold | IVs | Within groups | Between groups | Ratio | Within groups | Between groups | Ratio |
| 1e-08 | 50.4 | 0.000577 | 0.0443 | 76.7 | 0.000729 | 0.00358 | 4.92 |
| 5e-08 | 63.4 | 0.000474 | 0.0472 | 99.4 | 0.000598 | 0.00468 | 7.83 |
| 1e-07 | 69.7 | 0.000445 | 0.0468 | 105.2 | 0.000561 | 0.00437 | 7.79 |
| 5e-07 | 90.5 | 0.000358 | 0.0494 | 137.7 | 0.000449 | 0.00536 | 11.93 |
| 1e-06 | 101.0 | 0.000324 | 0.0505 | 155.7 | 0.000409 | 0.00538 | 13.16 |

**Table S18: Analysis of variance in presence of uncorrelated pleiotropy**
$n_A = n_B = 20,000, \pi_x = 0.001, h_x^2 = 0.4, \kappa_x = 0.3, \kappa_y = 0.5, \alpha = 0.2, \pi_y = 0.002, h_y^2 = 0.3,$
60% of the causal SNPs for X were also causal for Y
For each threshold, the mean number of instruments used (IVs), the within groups and between group variances, their ratio (between/within) for both IVW-based and corrected effects are reported.

| | | IVW-based effects | | | Corrected effects | | |
|---|---|---|---|---|---|---|---|
| Threshold | IVs | Within groups | Between groups | Ratio | Within groups | Between groups | Ratio |
| 1e-08 | 54.0 | 0.000579 | 0.0788 | 136 | 0.000728 | 0.00577 | 7.93 |
| 5e-08 | 68.8 | 0.000500 | 0.0786 | 157 | 0.000626 | 0.00629 | 10.05 |
| 1e-07 | 76.0 | 0.000468 | 0.0801 | 171 | 0.000589 | 0.00686 | 11.66 |
| 5e-07 | 97.3 | 0.000408 | 0.0875 | 215 | 0.000515 | 0.00950 | 18.47 |
| 1e-06 | 108.7 | 0.000384 | 0.0935 | 244 | 0.000487 | 0.01129 | 23.20 |

**Table S20: Analysis of variance in presence of moderate correlated pleiotropy**
$n_A = n_B = 20,000, \pi_x = 0.001, h_x^2 = 0.4, \kappa_x = 0.3, \kappa_y = 0.5, \alpha = 0.2, \pi_u = 0.0001, h_u^2 = 0.2, q_x = 0.4, q_y = 0.3$
For each threshold, the mean number of instruments used (IVs), the within groups and between group variances, their ratio (between/within) for both IVW-based and corrected effects are reported.

| | | IVW-based effects | | | Corrected effects | | |
|---|---|---|---|---|---|---|---|
| Threshold | IVs | Within groups | Between groups | Ratio | Within groups | Between groups | Ratio |
| 1e-08 | 54.7 | 0.000952 | 0.182 | 191 | 0.001234 | 0.0102 | 8.3 |
| 5e-08 | 69.1 | 0.000879 | 0.195 | 221 | 0.001147 | 0.0140 | 12.2 |
| 1e-07 | 76.8 | 0.000774 | 0.198 | 255 | 0.001010 | 0.0150 | 14.9 |
| 5e-07 | 97.9 | 0.000736 | 0.216 | 294 | 0.000965 | 0.0198 | 20.5 |
| 1e-06 | 109.3 | 0.000657 | 0.228 | 346 | 0.000863 | 0.0220 | 25.5 |

**Table S22: Analysis of variance in presence of strong correlated pleiotropy**
$n_A = n_B = 20,000, \pi_x = 0.001, h_x^2 = 0.4, \kappa_x = 0.3, \kappa_y = 0.5, \alpha = 0.2, \pi_u = 0.0005, h_u^2 = 0.3, q_x = 0.5, q_y = 0.7$
For each threshold, the mean number of instruments used (IVs), the within groups and between group variances, their ratio (between/within) for both IVW-based and corrected effects are reported.

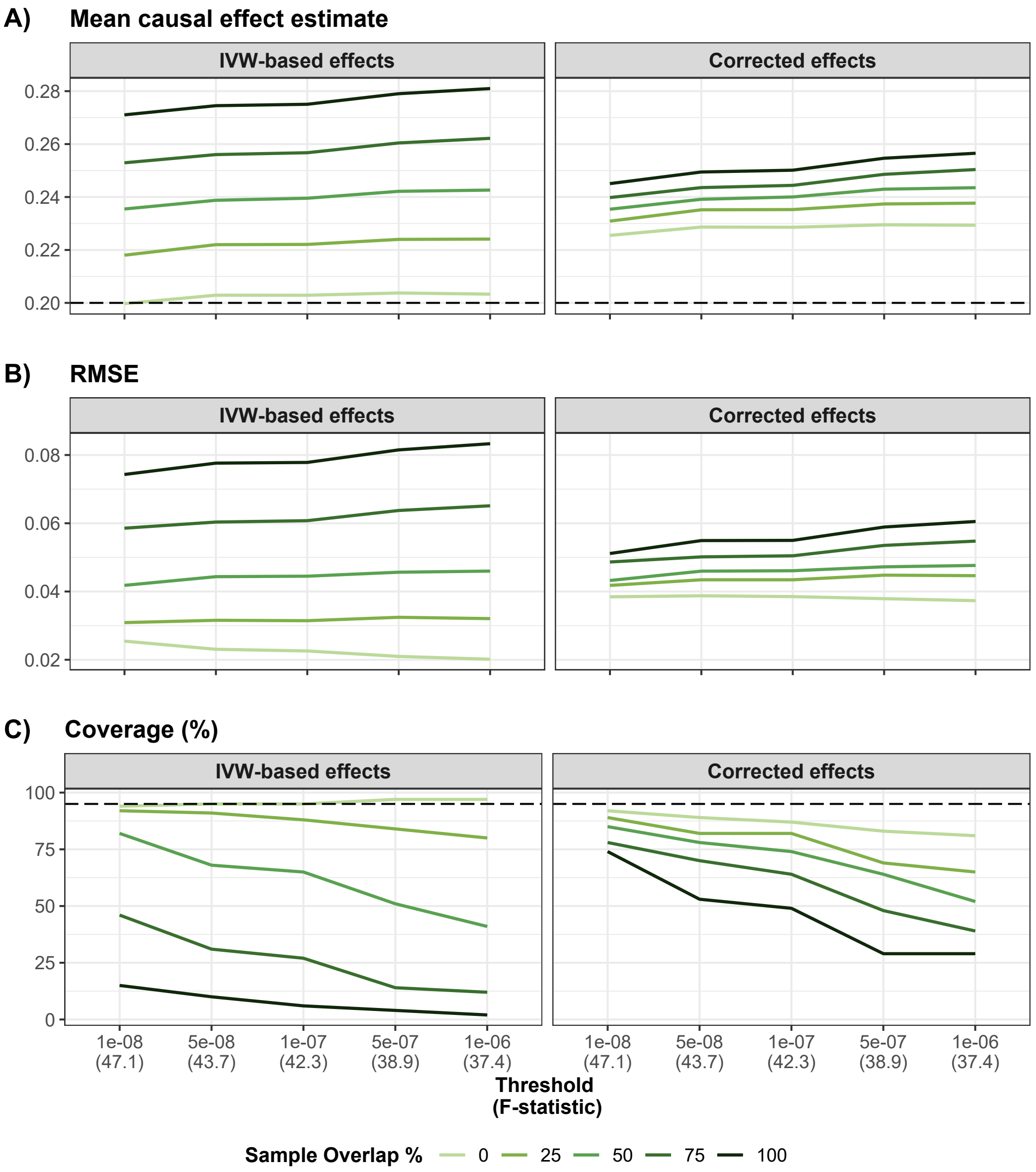


**Figure S12: Simulation results in presence of moderate correlated pleiotropy**
$n_A = n_B = 20,000, \pi_x = 0.001, h_x^2 = 0.4, \kappa_x = 0.3, \kappa_y = 0.5, \alpha = 0.2, \pi_u = 0.0001, h_u^2 = 0.2, q_x = 0.4, q_y = 0.3$
Panel A) shows the mean IVW-based and corrected effect for each overlap and threshold obtained from 100 simulations (the dashed line represents the true causal effect). Panel B) shows the mean RMSE obtained for IVW-based and corrected effect for each overlap and threshold. Panel C) shows the coverage of the 95% confidence interval for IVW-based and corrected effect for each overlap and threshold.

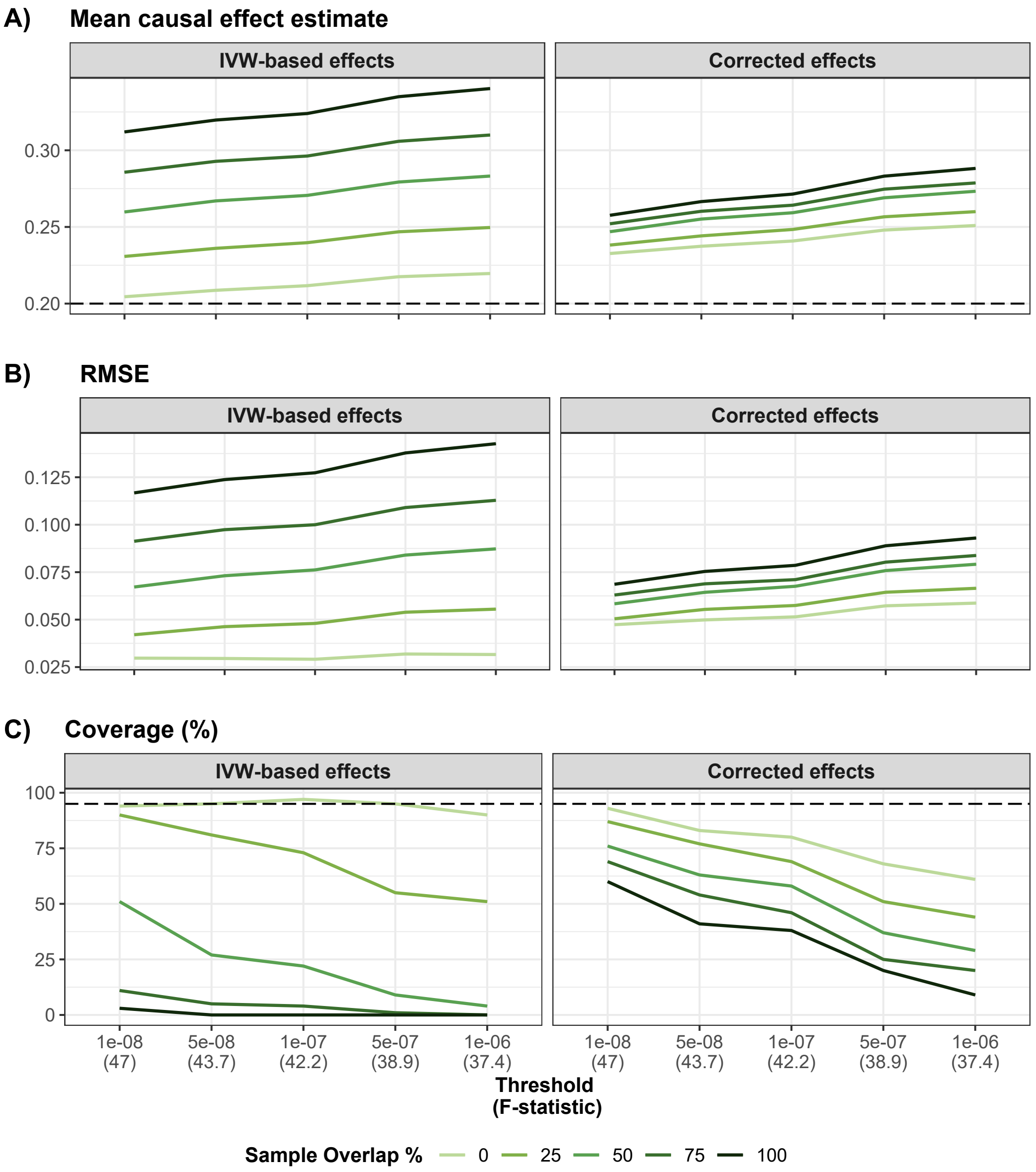


**Figure S13: Simulation results in presence of strong correlated pleiotropy**
$n_A = n_B = 20,000, \pi_x = 0.001, h_x^2 = 0.4, \kappa_x = 0.3, \kappa_y = 0.5, \alpha = 0.2, \pi_u = 0.0005, h_u^2 = 0.3, q_x = 0.5, q_y = 0.7$
Panel A) shows the mean IVW-based and corrected effect for each overlap and threshold obtained from 100 simulations (the dashed line represents the true causal effect). Panel B) shows the mean RMSE obtained for IVW-based and corrected effect for each overlap and threshold. Panel C) shows the coverage of the 95% confidence interval for IVW-based and corrected effect for each overlap and threshold.

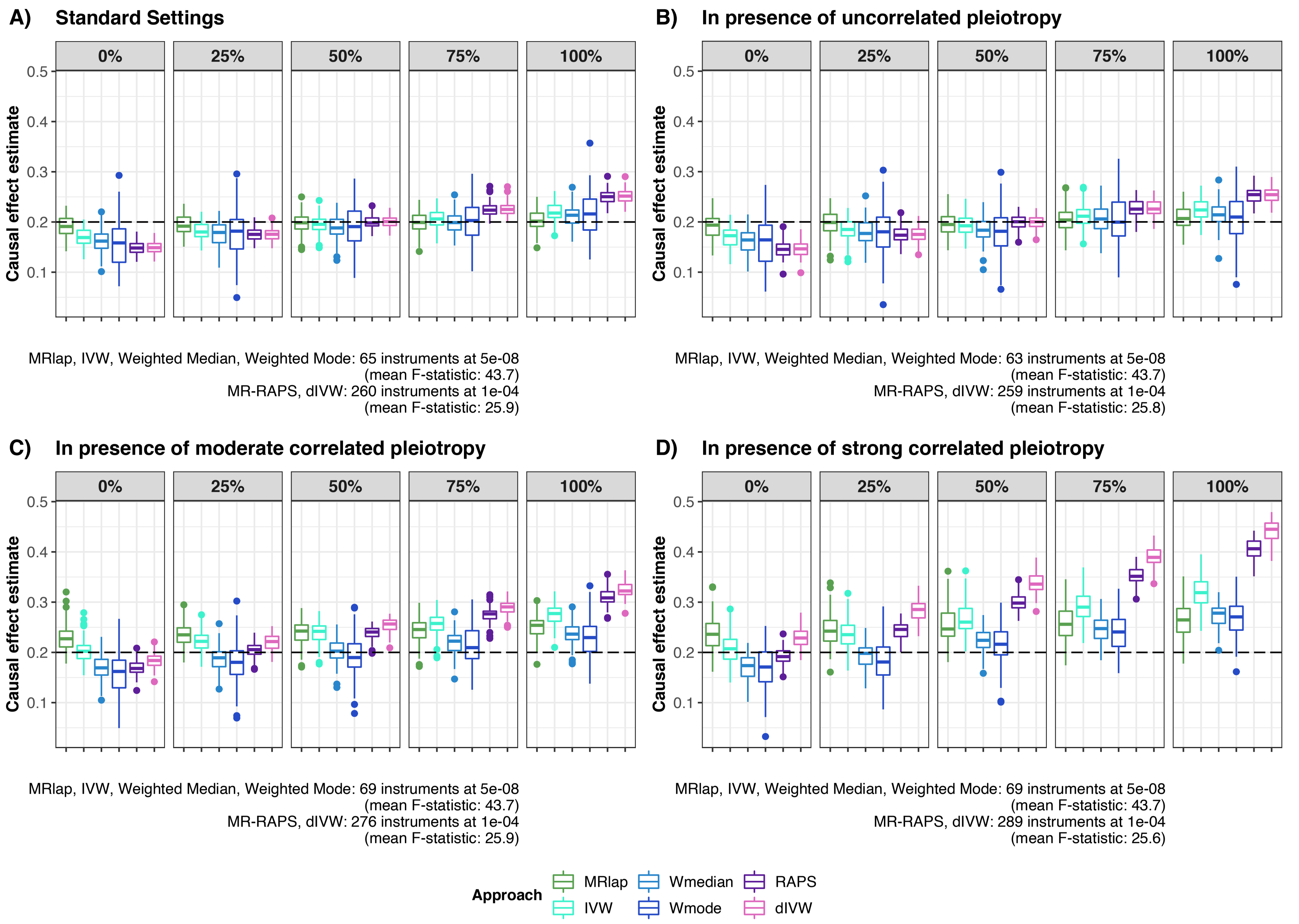


**Figure S14: Comparison of different MR approaches (less stringent threshold for MR-RAPS and dIVW)**
Causal effects estimates were obtained from 100 simulations using 6 different methods (MRlap in green, IVW in turquoise blue, Weighted Median in light blue, Weighted Mode in dark blue, MR-RAPS in purple and dIVW in pink). The dashed line represents the true causal effect. Panel A) shows results for the standard settings scenario (no pleiotropy). Panel B) shows results in presence of uncorrelated pleiotropy. Panel C) shows results in presence of moderate correlated pleiotropy. Panel D) shows results in presence of strong correlated pleiotropy. The average number of instruments and mean F-statistic for the different approaches (at 5e-08 / 1e-04) are indicated for each scenario.

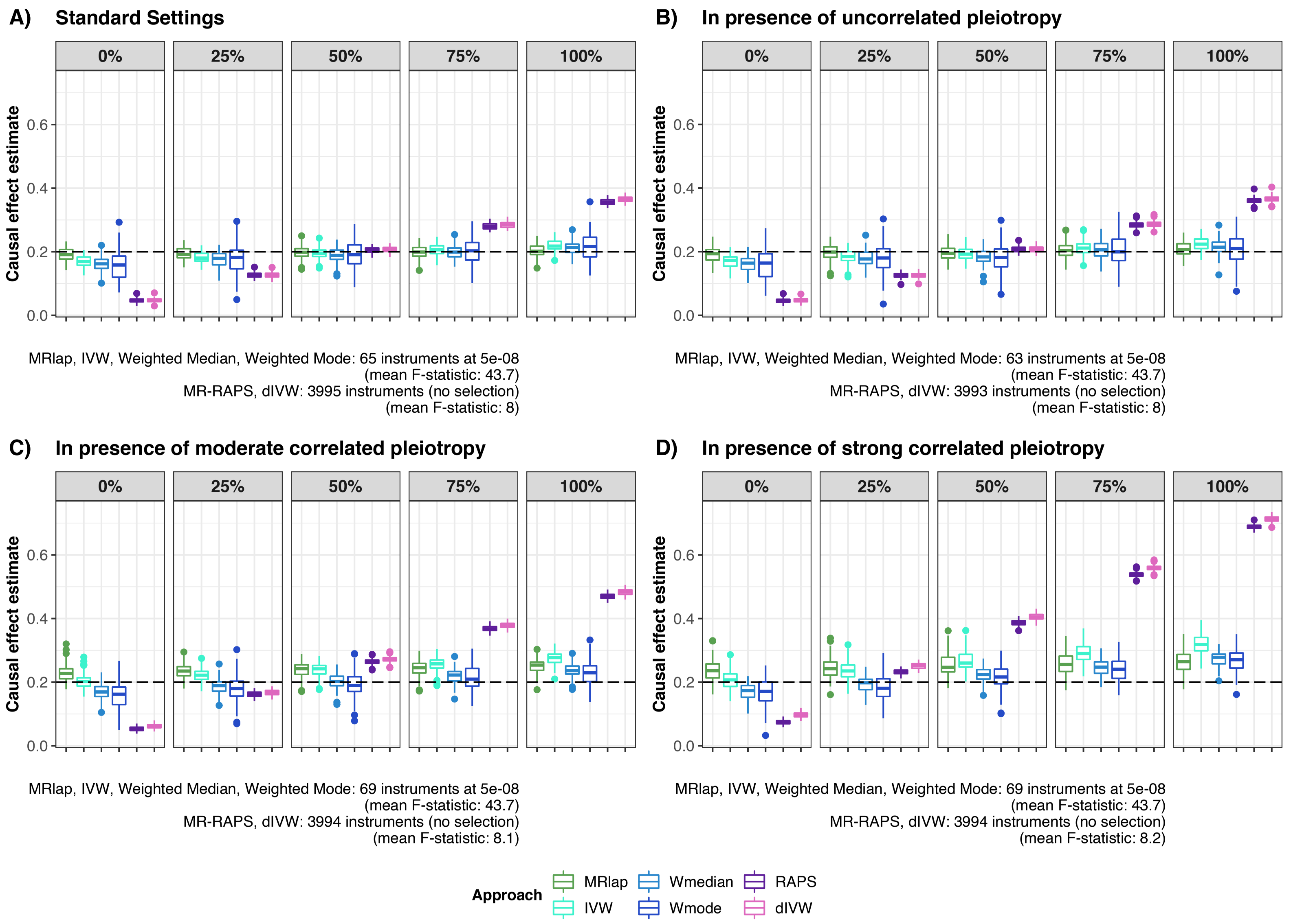


**Figure S15: Comparison of different MR approaches (no selection for MR-RAPS and dIVW)**
Causal effects estimates were obtained from 100 simulations using 5 different approaches (MRlap in green, IVW in turquoise blue, Weighted Median in light blue, Weighted Mode in dark blue, MR-RAPS in purple and dIVW in pink). The dashed line represents the true causal effect. Panel A) shows results for the standard settings scenario. Panel B) shows results in presence of uncorrelated pleiotropy. Panel C) shows results in presence of moderate correlated pleiotropy. Panel D) shows results in presence of strong correlated pleiotropy. The average number of instruments and mean F-statistic for the different approaches (at 5e-08 / without selection) are indicated for each scenario.

| | | IVW-based effects | | | Corrected effects | | |
|---|---|---|---|---|---|---|---|
| Threshold | IVs | Within groups | Between groups | Ratio | Within groups | Between groups | Ratio |
| 1e-08 | 27.4 | 0.001220 | 0.0103 | 8.48 | 0.00215 | 0.011240 | 5.231 |
| 5e-08 | 37.9 | 0.001068 | 0.0162 | 15.22 | 0.00195 | 0.007090 | 3.644 |
| 1e-07 | 44.3 | 0.001023 | 0.0193 | 18.86 | 0.00189 | 0.005630 | 2.976 |
| 5e-07 | 65.3 | 0.000717 | 0.0278 | 38.76 | 0.00139 | 0.003070 | 2.206 |
| 1e-06 | 79.0 | 0.000656 | 0.0368 | 56.00 | 0.00130 | 0.000758 | 0.583 |

**Table S26: Analysis of variance for the effect of BMI on SBP**
For each threshold, the mean number of instruments used (IVs), the within groups and between group variances, their ratio (between/within) for both IVW-based and corrected effects are reported.
*Since only results for a threshold of 5e-8 are discussed in the paper, results for all other thresholds have been greyed out.*

| | | IVW-based effects | | | Corrected effects | | |
|---|---|---|---|---|---|---|---|
| Threshold | IVs | Within groups | Between groups | Ratio | Within groups | Between groups | Ratio |
| 1e-08 | 27.7 | 0.00223 | 0.0114 | 5.09 | 0.00387 | 0.01013 | 2.615 |
| 5e-08 | 38.2 | 0.00191 | 0.0201 | 10.53 | 0.00345 | 0.00474 | 1.374 |
| 1e-07 | 44.3 | 0.00177 | 0.0223 | 12.60 | 0.00325 | 0.00423 | 1.300 |
| 5e-07 | 66.1 | 0.00150 | 0.0335 | 22.33 | 0.00289 | 0.00127 | 0.439 |
| 1e-06 | 79.3 | 0.00133 | 0.0436 | 32.86 | 0.00262 | 0.00120 | 0.460 |

**Table S28: Analysis of variance for the effect of BMI on smoking**
For each threshold, the mean number of instruments used (IVs), the within groups and between group variances, their ratio (between/within) for both IVW-based and corrected effects are reported.
*Since only results for a threshold of 5e-8 are discussed in the paper, results for all other thresholds have been greyed out.*

| | | IVW-based effects | | | Corrected effects | | |
|---|---|---|---|---|---|---|---|
| Threshold | IVs | Within groups | Between groups | Ratio | Within groups | Between groups | Ratio |
| 1e-08 | 28.4 | 0.000807 | 0.0100 | 12.4 | 0.001421 | 0.001576 | 1.109 |
| 5e-08 | 38.7 | 0.000746 | 0.0144 | 19.3 | 0.001356 | 0.000986 | 0.727 |
| 1e-07 | 44.9 | 0.000724 | 0.0161 | 22.2 | 0.001340 | 0.000876 | 0.653 |
| 5e-07 | 66.3 | 0.000592 | 0.0209 | 35.2 | 0.001147 | 0.000492 | 0.428 |
| 1e-06 | 79.2 | 0.000481 | 0.0228 | 47.4 | 0.000947 | 0.000467 | 0.493 |

**Table S30: Analysis of variance for the effect of BMI on alcohol**
For each threshold, the mean number of instruments used (IVs), the within groups and between group variances, their ratio (between/within) for both IVW-based and corrected effects are reported.
*Since only results for a threshold of 5e-8 are discussed in the paper, results for all other thresholds have been greyed out.*

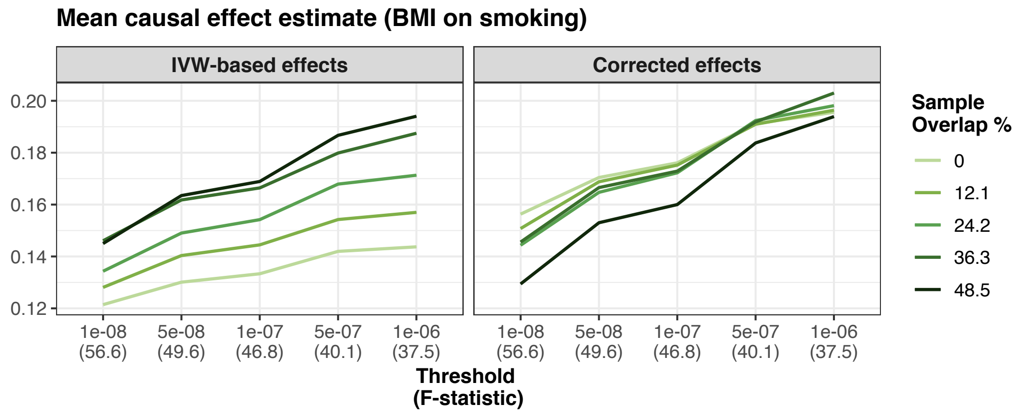


**Figure S16: Effect of BMI on smoking**
This figure shows the mean IVW-based and corrected effect for each overlap and threshold obtained from 100 different sampled datasets.

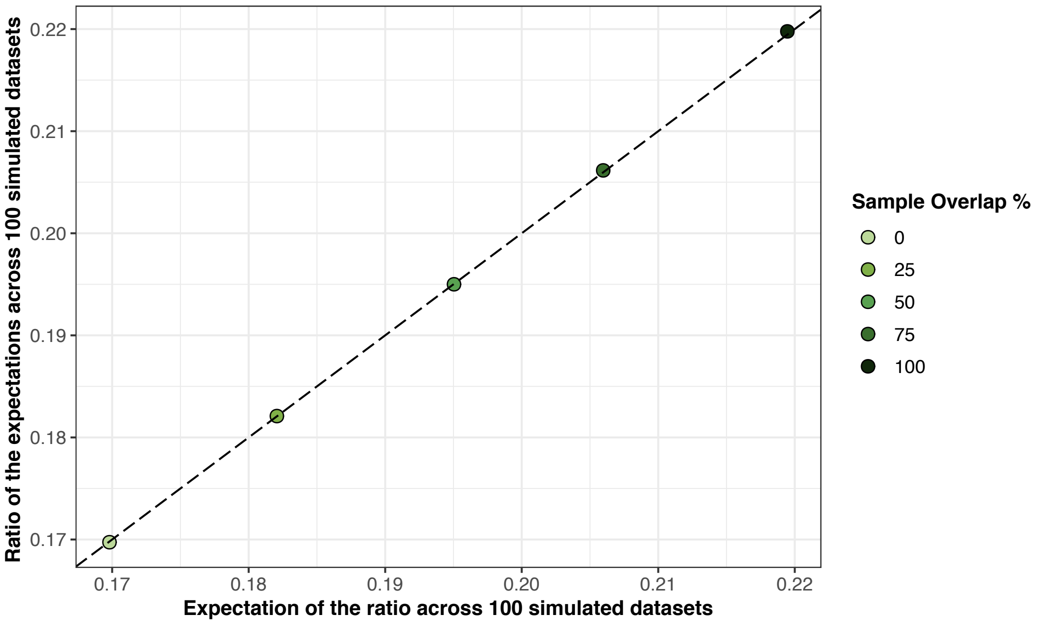


**Figure S17: Ratio of the expectation (simulated data)**
The IVW ratio, as well as the numerator and the denominator from (S16) were estimated for 100 datasets simulated using standard settings .
$n_A = n_B = 20,000, \pi_x = 0.001, h_x^2 = 0.4, \kappa_x = 0.3, \kappa_y = 0.5, \alpha = 0.2$ We reported the mean ratio (expectation of the ratio) estimated across the 100 simulated datasets and the ratio between the mean numerator / denominator standard deviation (ratio of the expectations) across the 100 simulated datasets, for different overlaps and a threshold of $5e-08$. The dashed line corresponds to the identity line.

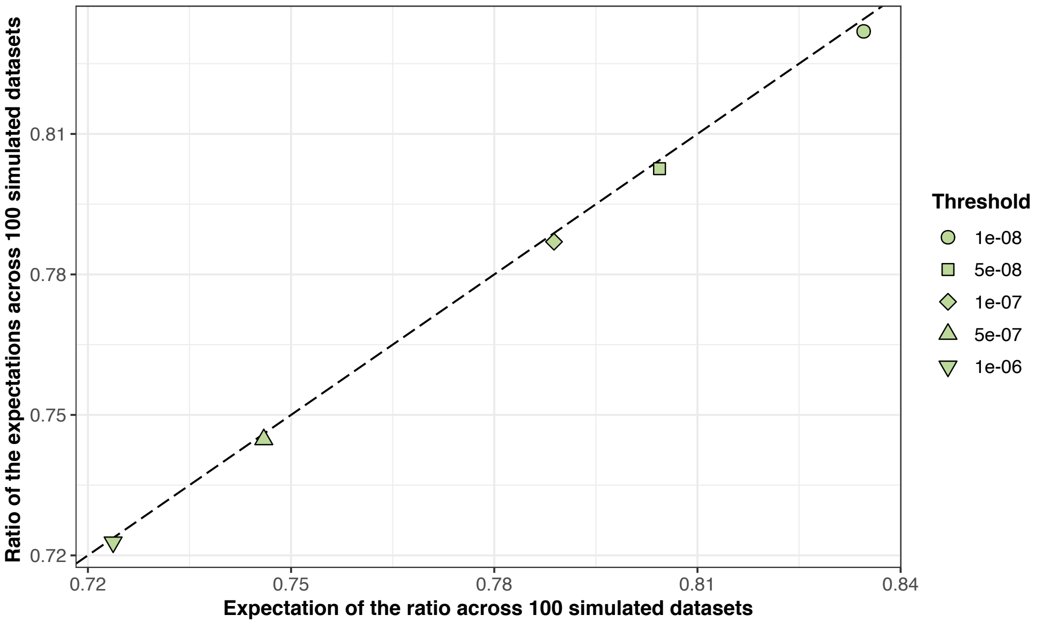


**Figure S18: Ratio of the expectation (BMI-BMI data)**
The IVW ratio, as well as the numerator and the denominator from (S16) were estimated for 100 sampled datasets from the BMI-BMI analysis .
We reported the mean ratio (expectation of the ratio) estimated across the 100 sampled datasets and the ratio between the mean numerator / denominator standard deviation (ratio of the expectations) across the 100 sampled datasets, for non-overlapping sampled and different thresholds. The dashed line corresponds to the identity line.

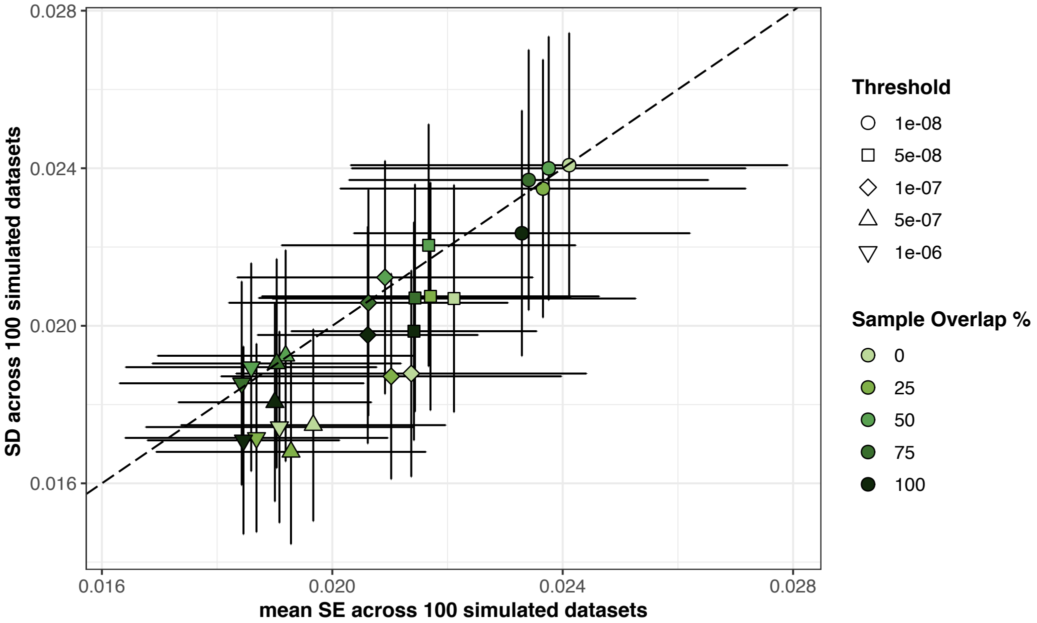


**Figure S19: Standard error estimation for the corrected effects**
The corrected effect standard error was estimated (as described in Supplementary Section C) for 100 datasets simulated using standard settings .
$n_A = n_B = 20,000, \pi_x = 0.001, h_x^2 = 0.4, \kappa_x = 0.3, \kappa_y = 0.5, \alpha = 0.2$ We reported the mean standard error (SE) estimated across the 100 simulated datasets and the observed standard deviation (SD) across the 100 simulated datasets, as well as their 95% confidence intervals, for different overlaps and thresholds. The dashed line corresponds to the identity line.
